# A cellular basis for the mammalian nocturnal-diurnal switch

**DOI:** 10.1101/2023.06.22.546020

**Authors:** Andrew D. Beale, Matthew J. Christmas, Nina M. Rzechorzek, Andrei Mihut, Aiwei Zeng, Christopher Ellis, Nathan R. James, Nicola J. Smyllie, Violetta Pilorz, Rose Richardson, Mads F. Bertelsen, Shaline V. Fazal, Zanna Voysey, Kevin Moreau, Jerry Pelletier, Priya Crosby, Sew Y Peak-Chew, Rachel S. Edgar, Madeline A. Lancaster, Roelof A. Hut, John S. O’Neill

## Abstract

Early mammals were nocturnal until the Cretaceous-Paleogene extinction enabled diurnal niche expansion. Diurnality evolved multiple times independently, but the mechanisms driving this shift remain unclear. We identify a conserved cell-intrinsic signal inversion that facilitates the transition from nocturnality to diurnality. Diurnal and nocturnal mammalian cells respond oppositely to temperature and osmotic cycles, mirroring species’ activity patterns. Cells exhibit differential global responses to temperature changes, including the phosphoproteome and protein synthesis. mTOR signaling is identified as a central mediator of this inversion, with diurnal mammals converging on modifications to mTOR and WNK pathways during evolution. Reducing mTOR activity induces nocturnal-to-diurnal shifting at cellular, tissue, and organismal levels. Therefore, the mTOR pathway is a cellular nexus that integrates energetic state and environmental signals to determine activity niche.

## Main text

Early mammals were nocturnal (night active) until extinction of the diurnal (day active) dinosaurs facilitated a rapid expansion into daytime niches (*1–3*). Diurnality subsequently arose multiple times, independently, from diverse and distant nocturnal lineages (*1*, *3*). No mechanistic basis for the switch between nocturnality and diurnality is known, though evidently some change in the relationship between internal circadian clocks and external daily rhythms is required (*4*, *5*).

Despite the 76 million years that separate nocturnal mice and diurnal humans from their common ancestor (*6*), the same cell-autonomous circadian clock mechanism operates in both mouse and human cells (*7*). Daily rhythms of gene expression, proteome renewal, and myriad cellular functions depend on cell-intrinsic ∼24h oscillations in the production of PERIOD (PER) proteins (*7*, *8*); where the changing activity of PER over time effectively determines the biological time-of-day (*9*). Similarly, the hypothalamic suprachiasmatic nucleus (SCN) performs an equivalent function in diurnal and nocturnal mammals, receiving light input directly from the eyes to generate an internal representation of solar time (*5*, *10–19*). However, unlike the SCN, PER oscillations in peripheral cells and tissues are oppositely organized between diurnal and nocturnal mammals (*18*, *20*, *21*), and instead vary with daily systemic signals that habitually coincide with the transition from resting, fasting and lower body temperature to activity, feeding and higher body temperature (*22*), rather than external solar time. Thus, excepting the SCN, the major behavioral and physiological daily rhythms in mammals are set to opposite times of day between nocturnal and diurnal mammals (Fig. 1A), suggesting a switch downstream of the SCN (*4*). How diurnal mammals integrate the same environmental cues to achieve an inversion of organismal and cellular physiology compared with nocturnal mammals is an open question whose answer is critical for understanding the internal synchrony that is pivotal for long-term health (*23–26*).

**Fig. 1.**
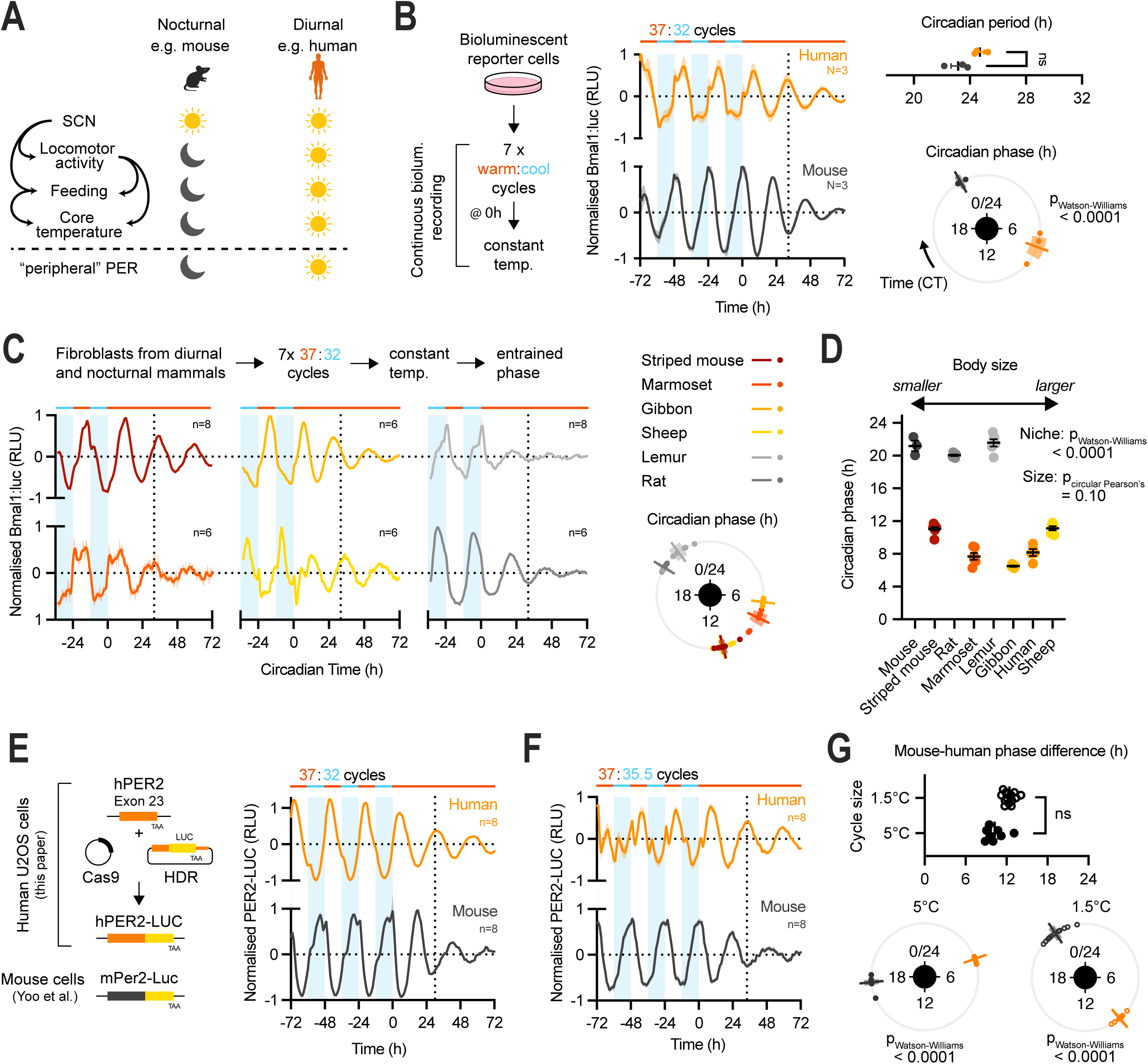
Entrainment to temperature is a cellular correlate of behavioral temporal niche. **(A)** With respect to the external day-night cycle, organismal and cellular physiology differ between nocturnal mice and diurnal humans, despite these species having the same cell-autonomous circadian clock mechanism. The retinorecipient hypothalamic SCN functions very similarly in diurnal and nocturnal mammals, with neuronal firing and PER oscillations peaking in the daytime, allowing the SCN to serve as an internal representation of the day-night cycle. However, outside of the SCN, organismal behaviour, physiology and cellular activity, including oscillations of PER, are oppositely organized between diurnal and nocturnal mammals and instead vary with daily systemic signals that consistently coincide with the transition from rest/fast to activity/feeding rather than external solar time. This suggests a switch, downstream of the SCN, that controls the appearance of diurnality. **(B)** After entrainment in 7 x 37°C (red) and 32°C (blue) 12h:12h cycles (data for final 3 cycles shown) which emulate the daily body temperature rhythm, mouse (grey) and human (orange) primary fibroblasts (derived from N = 3 individuals) are set to opposing phases when released into constant 37°C (red). Bioluminescence from *Bmal1:luc* was recorded under cycling and constant temperature portions of the experiment, and signal was detrended and normalized to aid visualization of circadian phase. A dotted line illustrates the inverse phases under constant conditions. Circadian period varies between individuals, as previously reported (*159*), but does not significantly differ between the species (Student’s t-test). However, circadian phase, defined as the time of the peak of *Bmal1:luc* relative to the last transition to 37°C, significantly differs (Watson-Williams test). **(C)** Primary fibroblasts from striped mouse (*Rhabdomys pumilio*), marmoset (*Callithrix jacchus*), gibbon (*Hylobates lar*), sheep (*Ovis aries*), rat (*Rattus norvegicus*) and lemur (*Lemur catta*) were cultured under 37°C and 32°C 12h:12h for 7 cycles (data for final 1.5 cycles shown) before release into constant 37 °C. Bioluminescence from *Bmal1:luc* was recorded throughout (n=6-8). A dotted line at t= 32h illustrates the different phases under constant conditions. **(D)** Circadian phase of the *Bmal1:luc* rhythm in constant conditions for the 8 mammalian species from (B) and (C). Two distinct clusters of opposing circadian phases of entrainment are found in mammalian cells. Phase was significantly different between temporal niche (p_Watson-Williams_ < 0.0001) and did not correlate with body size rank (π_circular Pearson’s_ = −0.25; p = 0.10). Statistics: Temporal niche, Watson-Williams; Ranked body size, circular Pearson’s. **(E)** Human (U2OS) and mouse (fibroblasts, immortalized) expressing PER2-LUC from its endogenous locus, entrain to cycles of temperature (12h 37°C: 12h 32°C, data for 3 cycles shown) with opposing phases (n=8). Note the high amplitude rhythms during temperature cycles that damp upon entry into constant temperature condition, and the different phase of entrainment between the two different reporters, *Bmal1:luc* (B) and PER2-LUC (E). **(F)** Human (U2OS) and mouse (fibroblasts, immortalized) PER2-LUC cells were synchronized by medium change at t=-186h and subject to temperature cycles (12h 37°C: 12h 35.5°C, data for final 3 cycles shown)) set 6h out of alignment to original phase. Human and mouse cells re-entrain to the new timing cue within 7 days, and exhibit a stable and opposite entrained phase in constant conditions. **(G)** Circadian phase for mouse (grey) and human (orange), calculated from (D) and (E), is given as peak of PER2-LUC expression relative to the last transition to 37°C, and significantly differs between mouse and human at each cycle magnitude (Watson-Williams test). Phase difference between mouse and human at cycle magnitudes of 5°C (solid circles) and 1.5°C (open circles) is not significantly different. Statistics: Phase difference, t-test; circular phase, Watson-Williams.

At the cellular level, acute stimulation of PER and/or global protein synthesis elicits similar shifts in the timing (or phase) of subsequent PER oscillations in both mouse and human cells (*9*, *27–29*). Physiologically, daily PER oscillations in cells throughout the body are synchronized and amplified by behavioral patterns of feed/fast, rest/activity, light/dark and stress exposure acting *via* specific systemic signals (*29–35*), a process known as circadian entrainment (*36*, *37*). Hormonal entrainment by insulin signaling (*22*, *29*, *38*, *39*) and glucocorticoids (*30*, *31*) which signal patterns of feed/fast and light/dark respectively, occurs by similar mechanisms in human and mouse (*29*, *40*), and should reinforce the differential behavioral patterns that drive the daily release of these hormones. Cellular clocks throughout the body and brain can also be synchronized by daily rhythms in body temperature that associate with locomotor/feeding thermogenesis during wakeful activity and increased cooling *via* peripheral vasodilation during sleep/rest (*41–48*). Whether temperature-mediated timing cues act comparably on cells from diurnal and nocturnal mammals has not been investigated, however.

Circadian synchronization by temperature is typically weaker than hormonal stimulation, with heat shock pathways (*46*, *47*), cold shock proteins (*49*, *50*), cdc-like kinases (*51*) and upstream open reading frames (uORFs) in PER mRNA (*52*) having each been independently proposed to communicate temperature change to the cellular clock by a range of transcriptional and post-transcriptional mechanisms (*53–55*). As these pathways are evolutionarily conserved, circadian responses to temperature change are assumed to operate analogously in mouse and humans and other mammals. Mouse and human biology can differ markedly, however, beyond obvious developmental differences (*56*). For example, mouse and human cells exhibit profoundly different biochemical reaction rates (*57–60*).

Here, we show that the mTOR signaling pathway activity and downstream sensitivity of bulk protein synthesis to temperature is a fundamental difference between nocturnal mice and diurnal humans, with profound physiological consequences that include the nocturnal-diurnal switch. We identify cell-autonomous differences between nocturnal and diurnal mammals in their response to thermal and osmotic challenge by specific (PER2 protein synthesis) and general (global phosphorylation and translation rate) mechanisms. We recapitulate temporal niche selection *in vitro* and reveal its cellular and molecular bases as a thermodynamic, not kinetic, effect. Finally, we test the functional consequences of modifying protein synthesis rates on temporal niche *in vivo*, and pinpoint mTOR activity as a signaling nexus that integrates bioenergetic and thermodynamic cues into the cellular clock.

## Results

### Cellular circadian rhythms of nocturnal and diurnal mammals are differentially entrained by daily temperature cycles

Daily temperature cycles can synchronize circadian clocks in cultured mammalian cells and modulate the timing of clock protein activity *in vivo* (*44*, *46*, *47*, *61*). Most systemic timing cues elicit very similar effects on the circadian clocks of mouse and human cells when delivered *in vitro* and reflect the activation of cell-autonomous response pathways ((*29*, *40*); fig. S1, A and B). We were therefore surprised to find that, using a conventional clock reporter (*Bmal1:luciferase*, *Bmal1:luc*), primary fibroblasts from multiple mice and humans consistently entrained oppositely to 12h:12h 37°C:32°C temperature cycles that mimic daily body temperature rhythms (Fig. 1B, fig. S1C, (*43*, *44*, *47*, *62*)). This is evident from bioluminescence waveforms that rapidly become antiphasic to each other during the temperature cycle and subsequently persist at constant temperature, and suggests a fundamental difference in the way cells from the two species respond to temperature.

This differential synchronization by temperature mirrors the opposite temporal niches habitually occupied by mice and humans, so we sought to test the generality of our findings using primary fibroblasts from a range of diurnal and nocturnal mammals. In natural environments, humans, gibbons, marmosets (*63*), sheep (*64*) and striped mice (*65*) occupy diurnal niches whereas rats, mice and many species of lemurs (*63*, *66*) are typically nocturnal. Remarkably, we found that, after temperature cycles, nocturnal representatives consistently entrained oppositely to cells from diurnal species (Fig. 1C) with no significant difference in circadian period between temporal niche (fig. S1, D and E). The difference in phase did not associate with body size (Fig. 1D).

To validate our findings and facilitate deeper mechanistic investigation, we repeated these experiments using an alternative reporter, PER2-LUCIFERASE (PER2-LUC aka PER2::LUC). PER2-LUC is a well-established, reliable reporter of the molecular clock in mammalian cells and tissues (*67*, *68*), since resultant bioluminescence correlates directly with the nascent production of endogenous PER protein (*69*). We compared fibroblasts from PER2-LUC mice (*67*) with human PER2-LUC knock-in U2OS cells generated using CRISPR-Cas9 (fig. S1, F-G). Again, we found mouse and human cells quickly developed oppositely phased rhythms under a 5°C daily temperature cycle that were maintained under constant conditions (Fig. 1E, fig. S1, H and I).

To confirm our findings were not attributable to any thermal stress response we repeated these experiments with a smaller 1.5°C temperature cycle. Again, we observed that human PER2-LUC showed an inverted phase relative to mouse PER2-LUC rhythms (Fig 1, F and G), with the only difference being that the absolute phases relative to the temperature cycle differed (Fig. 1G). This is consistent the theory that the phase of entrainment varies with the strength of stimulus (*36*). From these observations, we infer the existence of a cell-intrinsic signal inverter when diurnal mammalian cells are compared with nocturnal cells. We considered that understanding this signal inverter might provide insight into the nature of the mammalian nocturnal-diurnal switch.

### Diurnal cellular clocks are buffered against temperature change

During entrainment *in vivo*, the phase of cellular clocks is adjusted during each day by systemic signals in a fashion that varies with the magnitude of each stimulus and the relative biological times (circadian phase) at which they are received (*70*, *71*). Having found that daily temperature cycles, in the absence of other synchronizing cues, elicited opposite effects on diurnal vs nocturnal cellular clocks, we sought to elucidate the mechanism of signal inversion by using thermal challenge as a tool. Single temperature shifts are sufficient to adjust cellular clock phase (*46*, *52*); we therefore asked whether differential synchronization by temperature cycles between mouse and human cells is due to differences in their response to the same temperature shift. As such, mouse and human PER2-LUC cells were subjected to a single temperature increase or decrease at different circadian phases (fig. S2, A and B). From the phase response curves (fig. S2, C and D), it was evident the circadian clock was indeed differentially sensitive to single temperature steps between mouse and human cells. Whilst the qualitative response to temperature change was similar, over most of the circadian cycle, human cells showed greater advances for temperature increase whereas mouse cells showed greater delays for temperature decrease. Under a daily temperature cycle, this is sufficient to result in opposite entrainment (fig. S2E) and superficially explains the cellular phenomenon but not its underlying mechanism.

Acute changes in PER protein production shift the phase of cellular clocks *in vitro* and *in vivo* (*9*, *27*, *29*). We therefore asked whether differential sensitivity of mammalian cellular clocks to temperature change was reflected at the level of PER synthesis (Fig. 2A). We drew on understanding of firefly luciferase enzyme kinetics (*69*, *72*, *73*) to deconvolve the acute response to a rapid 5°C temperature increase (Fig. 2B) into two components. First, change in the baseline due to change in catalytic turnover of luciferase, which was not different between mouse and human cells (Fig. 2C); second, the change in total and peak luminescence that reflects the induction of PER2-LUC protein synthesis, which occurred more rapidly and produced more nascent PER2 in mouse than human cells (Fig. 2D). At lower luciferin concentrations, which reflect steady-state PER2-luciferase concentration (*69*), the luciferase signal did not change with temperature over these short timescales (fig. S3, A and B), suggesting it is the synthesis of the PER2 protein which is responding to the temperature change with a different magnitude in mouse than human cells. Over several cycles then, in principle, species-specific differences in the thermal sensitivity of PER protein production could function cumulatively to invert cellular clock timing. We therefore asked by what mechanisms temperature-dependent translation of PER2 might differ between species. We considered this must either occur by mechanisms that selectively regulate PER or by more general mechanisms that include changes in PER expression and activity.

**Fig. 2.**
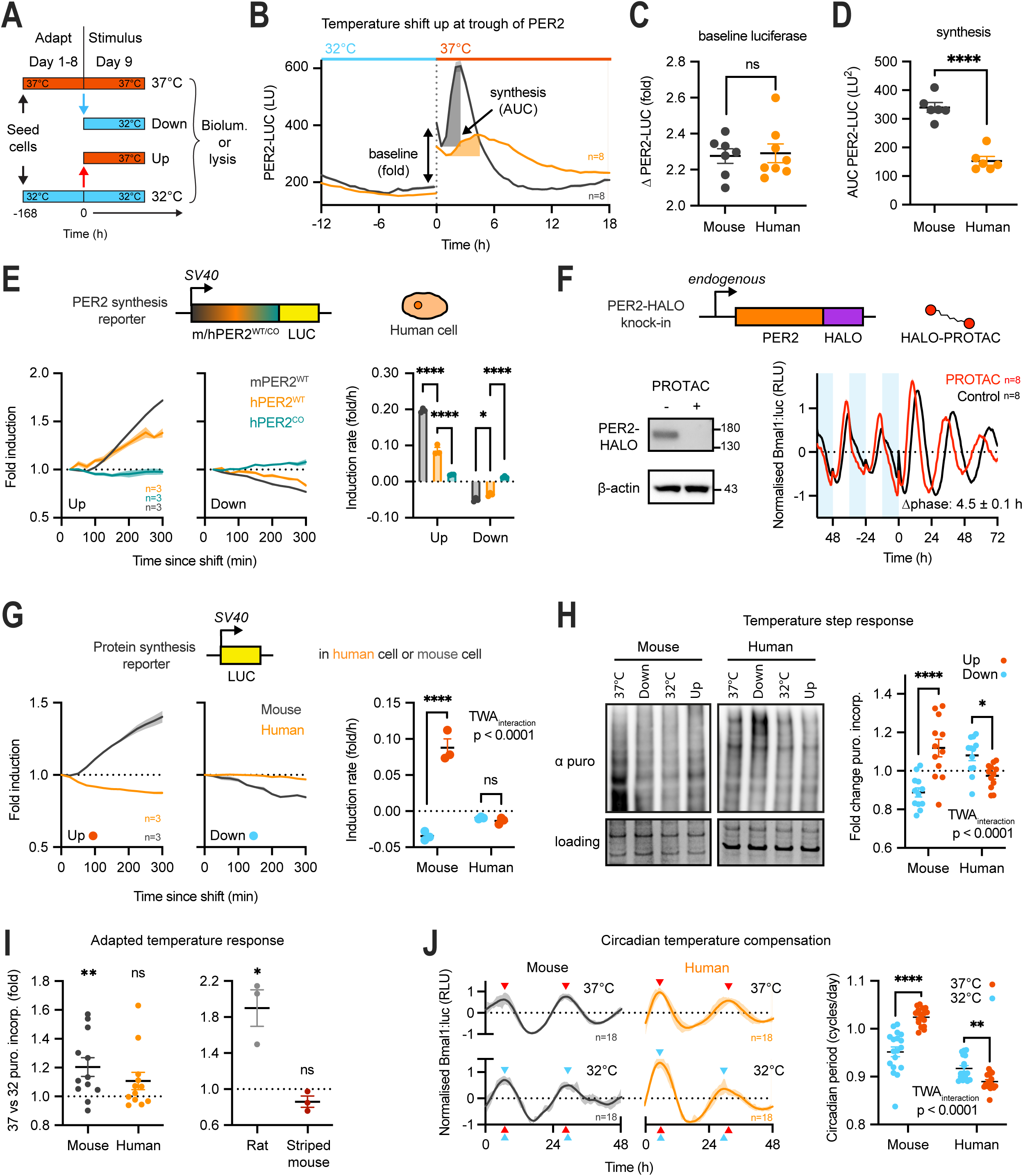
Differential response to temperature is both a specific property of PER2 and global translation. **(A)** Schematic of acute temperature shift and long-term temperature adaptation experiments. Cells are maintained at the indicated temperature (37°C or 32°C) for >6 days before temperature shifts (5°C shifts) occur. **(B)** Raw bioluminescence data (arbitrary luminescence units, LU) showing preceding 24h and ensuing 36h of cells exposed to a temperature shift up from 32°C to 37°C at 0h (trough of PER2). Change in baseline, Δ baseline, due to temperature dependence of ^app^K_m_ of luciferase is indicated by arrow, and integration of luciferase signal, ϕ synthesis, due to synthesis of new PER2-LUC is indicated by shading. **(C)** and **(D)** Quantification of Δ baseline, (C), and ϕ synthesis, (D), after a temperature shift up at 0 h in mouse and human lines. Mean ± SEM and individual points presented throughout. Statistics: Student’s t-test, unpaired. **(E)** (Left) U2OS cells stably expressing constitutive PER2-LUC fusions – human PER2-LUC (orange), mouse PER2-LUC (grey) or codon-optimized human PER2-LUC (teal) – were kept for 3 days at constant temperature of either 32°C or 37°C before shifting temperature at t = 0 min. Fold induction of luminescence is shown relative to t = 0 min. (Right) Rate of induction was quantified as the gradient of the straight line fit from non-linear regression (Prism), given as fold/hour. Induction rates were compared between reporters. Statistics: TWA followed by Šídák’s post-hoc test. **(F)** U2OS cells expressing HaloTag from the endogenous PER2 locus were treated with 1 µM HaloPROTAC3 or DMSO control. (Left) Treatments were applied to tagged cells for 24h before lysis and immunoblotting. Anti-Halotag antibody was used to detect the presence of the fusion PER2-HALO protein, anti β-actin serves as a loading control. (Right) PER2-HaloTag U2OS cells were cultured under temperature cycles ± 1 µM HaloPROTAC3 (n=4 each condition). At 0h cells were kept in constant temperature. Bioluminescence from *Bmal1:luc* was recorded throughout. Difference in phase of entrainment between control and PROTAC treatment is given, mean ± SEM. PROTAC treatment elicits an advance in entrained phase to temperature relative to control. **(G)** Human cells (U2OS) or mouse cells (NIH 3t3) stably expressing constitutive LUC as a reporter of protein synthesis were exposed to the same temperature conditions as (D). (Right) Rate of induction of constitutive LUC calculated as (D). **(H)** At 1.5h after temperature step, 10 µg/ml puromycin was added to the cells which were lysed 30 min later. Fold change puromycin incorporation was calculated by comparing incorporation in the stepped condition vs incorporation in the control condition. (Left) Representative immunoblot of one of three biological replicates of mouse or human cells exposed to the four conditions (constant 37°C, shift down from 37°C to 32°C, constant 32°C, shift up from 32°C to 37°C). Anti-puromycin (top) and coomassie loading control (bottom). (Right) Fold change puromycin incorporation in each direction temperature shift, N=12. Statistics: TWA mixed-effects model, followed by Šídák’s post-hoc test. Interaction species x temperature F(1, 44) = 30.47, p < 0.0001. **(I)** Fold change protein synthesis rate in cells adapted to constant 37°C vs constant 32°C (from Fig. 2H and fig. S6C) in (left) biological replicate primary fibroblast cells of mouse (N=12) or human (N=12), and (right) rat (n=3) or striped mouse (n=3) fibroblasts. Statistics: one sample t-test, H_0_ = 1. **(J)** Free-running period at constant 37°C (red) or 32 °C (blue) in biological replicate fibroblast cells of mouse (N=3, n=6) or human (N=3, n=6) expressing Bmal1:luc. Statistics: TWA mixed-effects model. Interaction species x temperature, F(1, 4) = 23.68, p = 0.0082, Šídák’s post-hoc test reported; (right) TWA. Interaction species x temperature F(1, 20) = 170.8, p < 0.0001, Šídák’s post-hoc test reported.

Compared with the clear mouse/human difference in PER2 translation and consistent with previous reports, we found no evidence for equivalent differences in the acute transcriptional response of *Per2* to temperature change (fig. S3, C and D (Miyake *et al*, 2023)). This suggests signal inversion occurs post-transcriptionally. As such, we note *Per2* mRNA contains a temperature-responsive upstream open reading frame (uORF) that modulates translation of the PER2 protein to temperature increases in the physiological range (*52*, *74*). The *Per2* uORF is highly conserved among nocturnal and diurnal mammals however (fig. S3E and (*52*), and so not an attractive candidate for species-specific differences in PER2 protein synthesis. In contrast, mouse or human PER2-LUC ORF expressed constitutively in mouse or human cells recapitulated the acute response of endogenous PER2-LUC to temperature change (Fig. 2E, fig. S4, A and B), suggesting differential thermal sensitivity of PER2 translation is largely intrinsic to the coding region without requiring 5’- or 3’-UTR regulation. Rare codon usage and RNA secondary structure are common mechanisms of translational regulation that affect the synthesis of many proteins (*75–79*), including those with circadian function such as PER2 (*80*, *81*). Consistent with this we found that, compared with wild type, codon-optimized hPER2 (hPER2-CO, fig. S4C) with less predicted mRNA structure (fig. S4D) showed minimal sensitivity to temperature change (Fig. 2E).

On the other hand, we identified only modest differences in codon usage and predicted mRNA structure between mouse and human PER2 (fig. S4, C and D), with mouse PER2 slightly more structured than human. Furthermore, PER1 in both species was highly similar (fig. S4, E and F), so we wanted to distinguish whether PER2 itself was essential for the signal inverter in diurnal cells, or else simply associated with it. To directly test the contribution of PER2 to circadian synchronization by temperature cycles, we used CRISPR-edited cells where endogenous HALO-tagged PER2 could be acutely depleted using HALO-PROTAC3 (Fig. 2F, and fig. S5). Critically, when PER2 was acutely depleted, we found significant but only modest differences in the effect of daily temperature cycles on the diurnal cellular clock (Fig. 2F), consistent with previous reports (*52*). Therefore, whilst species-specific differences in PER2 translation may contribute to differential effects of temperature, they cannot be the sole basis for cellular signal inversion. From these data, we do not discount differences in the individual contributions of many other proteins, such as PER1. However, an alternative hypothesis is that general diurnal/nocturnal differences in the temperature-dependence of the translational machinery underlie the observed PER2 translational differences. This hypothesis is informed by recent developmental studies, showing marked differences in global biochemical reactions between species, with humans exhibiting generally slower rates and more stable proteins than mice (*57–59*). We therefore asked whether broader differences in the translational response to temperature change might underpin our observations.

Using constitutively expressed luciferase as a reporter for bulk 5’-cap-dependent translation, we found that mouse cells were much more sensitive to temperature increase and decrease than human cells (Fig. 2G). Mouse cellular translation increased with temperature increase, and *vice versa*, as previously reported (*82*, *83*). By contrast, human cells showed an inverted response with reduced magnitude: reduced translation for temperature increase and no significant change for temperature decrease. This inverted response of protein synthesis to temperature was particularly stark over repeated temperature cycles (fig. S6A). We validated these findings by quantifying nascent polypeptide production with puromycin-labelling in primary fibroblasts (Fig. 2H, and fig. S6B). Again, we found that protein synthesis in human cells was more resistant to physiological temperature change compared to mouse cells. The differential effect of temperature on translation rate was also observed over longer timescales: after 1 week at constant 32°C or 37°C, mouse cell protein synthesis was clearly temperature-dependent, faster at the higher temperature, whereas human cells showed no significant difference between the two (Fig. 2I). Translation in nocturnal rat cells likewise showed differential long-term temperature sensitivity compared with cells from the similarly sized but diurnal striped mouse (Fig. 2I, fig. S6C).

Circadian rhythms exhibit the remarkable feature of temperature compensation, where, unlike most biological processes, the ∼24h period of oscillation is only modestly affected by a change in ambient temperature (Q_10_ of 0.8-1.2) (*84*, *85*). However, consistent with their increased translational sensitivity to temperature, the cellular circadian rhythms of nocturnal mammals showed an increased temperature dependence relative to diurnal mammals. Mouse and rat circadian rhythms ran at a significantly faster rate at 37°C than 32°C, whereas circadian rhythms in human and striped mouse cells ran significantly slower at the higher temperature (Fig 2J, fig. S6D). Taken together, this suggests that biochemical reactions are more sensitive to temperature and run faster at higher temperatures in nocturnal mammalian cells compared to diurnal species. This provides an additional insight into cellular signal inversion that is complementary to the acute differences in thermosensitivity described above: during daily temperature cycles nocturnal cellular clocks accelerate at the higher temperature, whereas diurnal ones tend to slow down.

### Global species differences in the response to temperature change involve mTORC1 and WNK1

What causes differences in the temperature sensitivity of the protein synthesis machinery at the molecular level? Protein synthesis is principally controlled by phosphorylation of proteins comprising the translational apparatus (*86*), including members of the cap-binding complex eIF4F, 43S preinitiation complex, and the elongation factor eEF2 (*87*). To gain insight into differential responses to temperature, we performed quantitative (phospho)proteomics on biological replicates of primary mouse and human fibroblasts subjected to high or low temperature over either acute or extended time frames (Fig. 3A, fig. S7A). We reasoned that thermosensitive phosphosites could impart directionality to temperature signals, including those that collectively control translation rate. To identify potential thermosensitive phosphosites we focused our analysis on those where phosphorylation changed in proportion (fold change increases with temperature increase and *vice versa*) or in inverse proportion (fold change decreases with temperature increase and *vice versa*) with acute temperature change or longer-term temperature adaptation (Fig. 3, B and C and table S1).

**Fig. 3.**
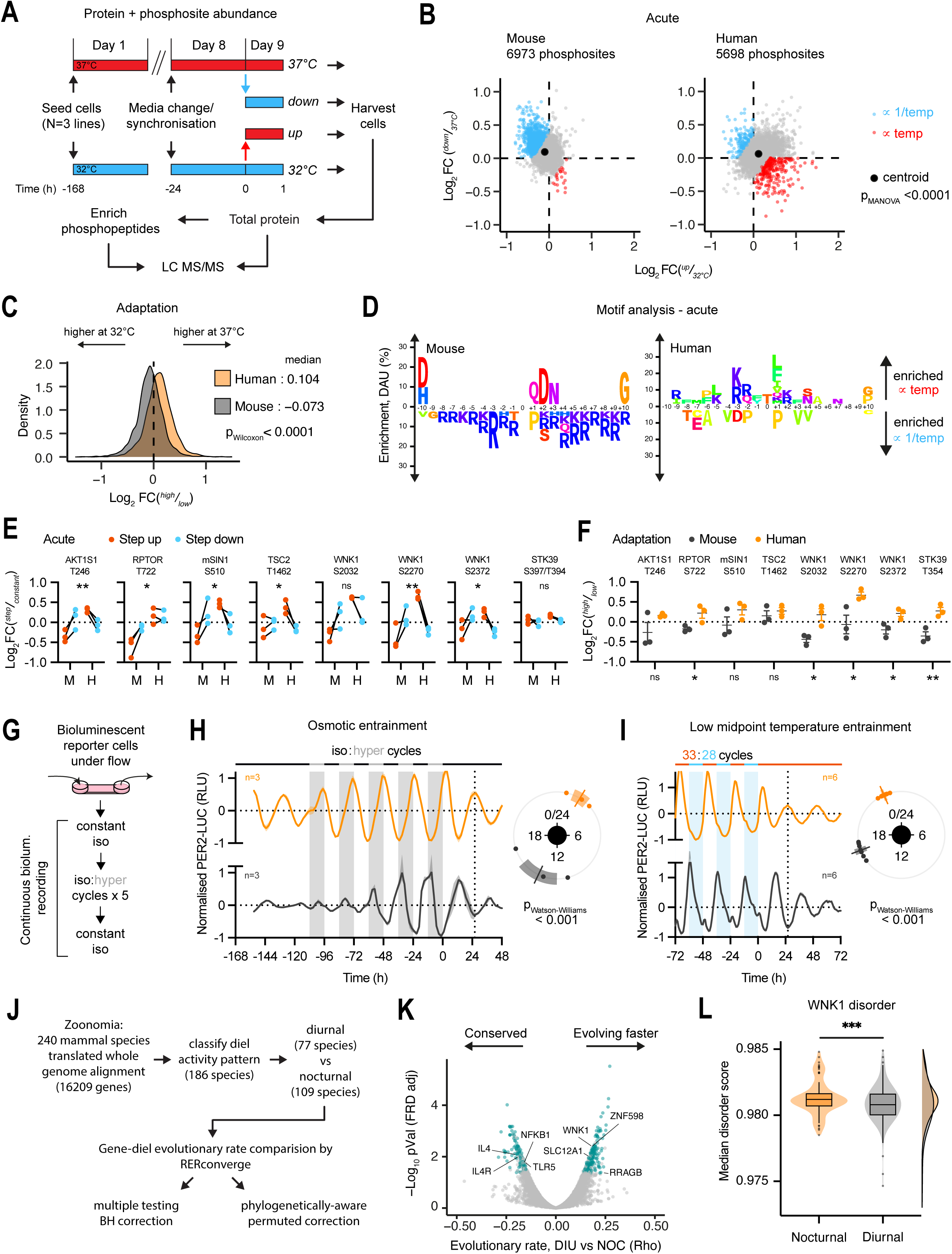
Differential mTOR pathway activity as the basis of the nocturnal-diurnal switch. **(A)** Schematic of temperature shift experiment for (phospho)proteomics. Human and mouse primary fibroblasts were kept for 1 week in constant temperature of either 32°C (blue) or 37°C (red). At t = −24h, cells were treated with 100 nM dexamethasone to synchronize and t = 0h cells either shifted up, down or kept at the constant temperature as a control (either 32°C or 37°C). Cells were lysed 1h later and quantitative proteomics (TMT-MS/MS) was performed to analyze the (phospho)proteome. For each phosphosite or peptide, fold change upon a temperature step is calculated by dividing (phospho)peptide signal of the shifted condition by the constant condition from which they were shifted. Long-term adaptation to temperature was examined by calculation of fold change (phospho)peptide signal from constant 37°C by signal at constant 32°C. **(B)** Phosphoproteomics matrices for mouse (6973 phosphopeptides) and human (5698 phosphopeptides). Matrix shows fold changes upon shift up (x-axis) and shift down (y-axis) for each phosphopeptide. Phosphopeptides are classified where fold change significantly increases with increasing temperature and decreases with decreasing temperature (proportional, ∝ temp, red) or significantly decreases with increasing temperature and increases with decreasing temperature (inversely proportional, ∝ 1/temp, blue). Phosphopeptides that do not change significantly, or change significantly but in a single direction, are shown in grey. Total phosphoproteome fold changes were compared by MANOVA, and centroids, representing the average direction of the phosphoproteome response to temperature shift, are plotted: mouse centroid (−0.028, 0.022); human centroid (0.047, 0.026). **(C)** Probability density distribution of fold change upon temperature adaptation for every detected phosphosite. Mouse (grey) and human (orange). Statistics: Mann-Whitney, p-value shown. **(D)** Motif analysis was performed on phosphopeptides that changed proportionally (∝ temp) or inversely proportionally (∝ 1/temp). Sequence logos showing enriched AA residues with significant differential AA usage (DAU) are shown for phosphopeptides that change proportionally (above) and inverse proportionally (below) for mouse (left) and human (right). Sequence logos are centered around the phosphoacceptor at position 0. Sequence logos showing under-represented AA residues (i.e. depleted) are shown in fig. S7. **(E and F)** Fold change of the abundance of significantly changing phosphosites of mTOR pathway members and WNK1 in human and mouse cells under acute shift (E) and adaptation conditions (F), extracted from (B) and (C). **(G)** Schematic of microfluidic-based entrainment of cells to repeated cycles of osmolarity. **(H)** Human (U2OS) and mouse (fibroblasts, immortalized) expressing PER2-LUC, were cultured under flow in isosmotic media (isosmotic relative to standard culture media) for 60h before exposure to cycles of osmolarity (12h iso-osmotic: 12h +50 mOsm) for 5 complete cycles and subsequent release back into isosmotic media for a final 48h. (H) Circadian phase of human and mouse cells relative to the final transition into isosmotic media. Circadian phase was compared with the Watson-Williams test. **(I)** Human (U2OS) and mouse (fibroblasts, immortalized) PER2-LUC cells entrain to cycles of temperature at below physiological levels (12h 33°C: 12h 28°C) with opposing phases. (Right) Circadian phase of human and mouse cells relative to the final transition into 33°C constant conditions. Circadian phase was compared with the Watson-Williams test. **(J)** Analysis pipeline for relative evolutionary rate of genes in nocturnal and diurnal mammals from the Zoonomia database. **(K)** Correlation values (Rho) between relative evolutionary rates of 16209 genes and phenotype, comparing 186 mammalian species classified as diurnal or nocturnal species, plotted against significance (-log_10_ p-value). Gene with significantly different evolutionary rates between diurnal and nocturnal are colored teal; a selection of genes are labelled. **(L)** WNK1 protein disorder was calculated per residue per species using Metapredict v3 (*160*) on amino acid sequences and alignments from the Zoonomia resource (*103*, *104*). The median disorder score of WNK1 for each species was compared by activity pattern (diurnal vs nocturnal).

We first noted the clear directional bias to the acute temperature response of the phosphoproteome (Fig. 3B). This directional bias in phosphoproteome response of the two species matches their directional bias in phase response (Fig. S2). A similar directional bias was observed in the temperature-adapted phosphoproteome (Fig. 3C), reflecting widespread differences in their homeostatic mechanisms of ambient thermo-adaption. Conversely, protein abundances were much less sensitive to acute or longer-term temperature change in both species (fig. S7, B to D and table S1).

There was, however, little overlap between mouse and human cells in the identity of temperature-dependent phosphosites and of the proteins to which they belong (fig. S7E). The pathways previously identified as regulators of circadian temperature response, HSF1 signaling (*46*, *47*) *via* HSP70 and HSP90, or the RNA binding proteins CIRBP and RBM3 (*49*, *50*) had similar proteomic responses to acute or long-term temperature change between mouse and human cells (fig. S7F). This aligns with the expected strong evolutionary conservation of the cellular response to temperature (*88–90*), but not with a role in a cell-intrinsic circadian signal inverter. Therefore, to examine alternative regulators, we performed motif analysis for amino acids surrounding the phosphoacceptor (S/T/Y) to identify the kinases and/or phosphatases that drive the observed phosphoproteomic differences upon temperature changes. We observed differences between mouse and human cells in both the direction and magnitude of response: in mouse, basic residues were highly enriched for inversely proportional phosphorylations; in humans this trend was reversed, apparent only in the −2/−3 positions, and with smaller magnitude (Fig. 3D, fig. S7G).

Basic residue motifs are recognized by diverse basophilic kinases, including those of the AGC family as well as With No Lysine/K (WNK) kinases (*91*, *92*). The AGC family include key regulators and effectors in the PI3K-AKT-mTOR pathway – the major pathway for control of protein synthesis, macromolecular crowding and cellular metabolism, whose activity reflects the integration of many different metabolic and extracellular signals to function as a ‘metabolic rheostat’ (*93–98*). WNK kinases are the master sensor/effectors of the WNK-OSR1/SPAK-SLC12A pathway that maintains intracellular water balance. Amongst the relatively small number of overlapping proteins in mouse and human cells that showed a difference in the direction of their phosphorylation response to temperature change (acute, <3%; adaptation, <1%; fig. S7E), a clear differential pattern was observed in key regulatory sites of WNK1 and mTOR pathway components (Fig. 3, E and F) consistent with enrichment for basic motifs in opposite directions. WNK1 and mTOR are ubiquitous essential proteins that function as major determinants of translation and cellular homeostasis more generally, and whose activities are coordinately and circadian-regulated in cultured cells and *in vivo* (*99–101*). We hypothesized that nocturnal-diurnal differences in their response to perturbation underlies the phenotypic switch.

### Convergent evolution of diurnal response to thermodynamic perturbation

Within concentrated macromolecule solutions like the cytosol, modest changes in temperature elicit large changes in the total thermodynamic potential energy of water. Water potential deviates from the linear relationship described by Van’t Hoff’s equation in both magnitude and direction as more water molecules are constrained within macromolecule hydration layers at lower temperatures with proportionally less ‘free’ molecules in bulk solvent, reducing the potential energy to perform work in the cell (*102*). Thermosensitivity can therefore be imparted to biological systems through either direct kinetic effects or components that respond to changes in solvent thermodynamics. These are easily distinguished by testing whether an equivalent change in water potential can mimic a temperature shift. For example, increasing external osmolarity would phenocopy decreasing temperature as ‘free’ water moves out of the cell by osmosis, reducing the intracellular water potential. WNK and mTOR signaling pathways are both sensitive to changes in water potential, for example, phosphorylation at OXSR1-S339 and AKT1-T450 scale directly with extracellular osmolarity and inversely with temperature (*102*). We therefore considered whether circadian entrainment to temperature change might occur by a thermodynamic mechanism and predicted, then demonstrated, that mouse and human cells differentially entrain to daily cycles in extracellular osmolarity (Fig. 3, G and H; fig. S8, A to D). Conversely, a nocturnal-diurnal switch that relies on kinetic effects would be sensitive to absolute temperature. We subjected mouse and human cells to temperature cycles with the same 5°C amplitude but a lower mid-point of 30.5°C compared to 34.5°C and they continue to entrain to opposing phases (Fig. 3I).

Collectively, these results support a model where WNK and mTOR pathways form an intrinsic nocturnal-diurnal switch by virtue of species-specific differences in their response to thermodynamic changes in the intracellular environment. Diurnality evolved several times, likely acting through complementary changes at many genetic loci that were assumed to differ between diurnal lineages. However, if changes in WNK and mTOR activity are an efficient evolutionary means to select for diurnal phenotypes then convergent evolution should be detected by comparative genomics. We therefore mined the Zoonomia comparative genomics resource of placental mammals (*103*, *104*) to ask whether members of these pathways are amongst those genes that evolved particularly quickly in the genomes of diurnal mammals relative to nocturnal mammals. Of the 242 species analyzed, 77 and 109 were categorized as definitively diurnal and nocturnal, respectively, based on prior literature (Fig. 3J and table S2). After restricting the analysis to ubiquitously expressed genes – excluding tissue-specific genes such as olfactory receptors (fig. S8, E and F) – to identify candidates that could contribute to our observed cellular phenotype, we found WNK1, RRAGB, a core regulator of mTOR complex 1 (mTORC1) activity (*105*), and translational quality control factor ZNF598 (*106*) were among the genes that have evolved significantly faster in diurnal mammals (Fig. 3K). Faster evolutionary rates in an additional key regulator of mTORC, TSC2, and a second paralogue of WNK1, WNK4, correlated with diurnality, but lay just outside our phylogeny-corrected significance threshold suggesting they are evolving faster in only a subset of related diurnal mammals (table S3).

The emergence of diurnality in mammals converges on mTOR and WNK pathway modifications, but how might these variations mechanistically lead to differential sensitivity to solvent thermodynamics? mTOR has many components and regulators that might impart ‘water responsiveness’, so we focused on WNK1. WNK1 autophosphorylation and activation is acutely sensitive to water potential (*100*, *102*, *107*). In cells, increased macromolecular crowding and the resultant decrease in solvent availability and potential energy drive WNK condensation, mediated by its intrinsically disordered C-terminal tail (*100*, *102*, *107–110*), an ensemble property of multiple sequence features rather than individual amino acid residues. Hydration of disordered regions has greater impact upon water potential than for compact structures, therefore they have increased likelihood of participating in compensatory biomolecular condensation to restore water equilibrium upon macromolecular crowding, thermal or osmotic challenge (*102*). We therefore predicted that diurnal WNK1 would contain less intrinsic disorder compared to nocturnal WNK1 as this would reduce the probability of (de)condensation upon temperature-driven changes in water potential and therefore the thermal sensitivity of WNK1 activity, reflecting the lower responsiveness of diurnal species to temperature change. We detected a significant difference in disorder between diurnal and nocturnal WNK1, the former tending towards less disorder as predicted and reflecting the lower responsiveness of diurnal species to temperature change (Fig. 3L, fig. S8G).

Comparative genomics therefore confirmed our hypothesis that the diurnal/nocturnal switch arose convergently and independently through multiple complementary mutations that act together to alter the cellular sensitivity (e.g. WNK pathway) and responsiveness (e.g. mTOR pathway) to perturbation of cellular thermodynamic equilibria by modulating the favorability of key macromolecular interactions. This differentially affects circadian phase *via* a combination of specific (PER synthesis) and more general mechanisms (basophilic kinase activity, bulk translation) that ultimately renders human circadian clocks more robust to thermal and osmotic perturbation than those in mice. Under repeated daily thermodynamic perturbations, this results in entrainment to opposing phases (fig. S2E). Ultimately, our results strongly suggest that cellular clocks respond to crowding-related changes in macromolecular hydration and supramolecular assembly rather than changes in solute kinetic energy, as was implicitly assumed.

### Perturbation of mTOR activity and translational initiation makes nocturnal cells behave like diurnal cells

This hypothesis makes a simple testable prediction: decreasing the basal protein synthesis rate by inhibiting mTOR activity should attenuate the capacity of nocturnal cellular clocks to respond to thermal challenge more than diurnal clocks, rendering them more diurnal-like by reducing the relative magnitude of temperature-dependent differences in translation between the two. Whereas a panel of small molecule inhibitors of proteins and kinases previously implicated in circadian post-translational regulation revealed only modest effects on entrained phase under daily temperature cycles (fig. S9, A to D), inhibitors targeting the mTOR signaling pathway showed large effects on entrained phase (Fig. 4A and fig. S9, E and F), with selective mTOR inhibition by INK128 showing the largest effect. When mTOR activity is reduced, mouse cells showed significant phase delays under daily temperature cycles, whereas human cells were relatively unaffected (Fig. 4A). INK128 treatment of fibroblasts from another nocturnal mammal (rat) and diurnal mammal (striped mouse) gave comparable entrainment phenotypes, demonstrating the conservation of the role of this pathway in temperature signaling (Fig. 4B). Growth factor signaling acts *via* mTORC1 to control protein synthesis rates (*93*, *94*), and can be manipulated in cell culture by changing serum concentration. In lower serum concentrations, mouse cellular rhythms were delayed by up to 6h under daily temperature cycles compared with high serum control conditions, whereas human cells were not (Fig. 4C). In all cases, suppression of mTOR activity makes cells from nocturnal mammals behave more like cells from diurnal mammals, with PER2-LUC peaks selectively shifting towards the early warm portion of the temperature cycle, whereas the contrary was not true for diurnal cells.

**Fig. 4.**
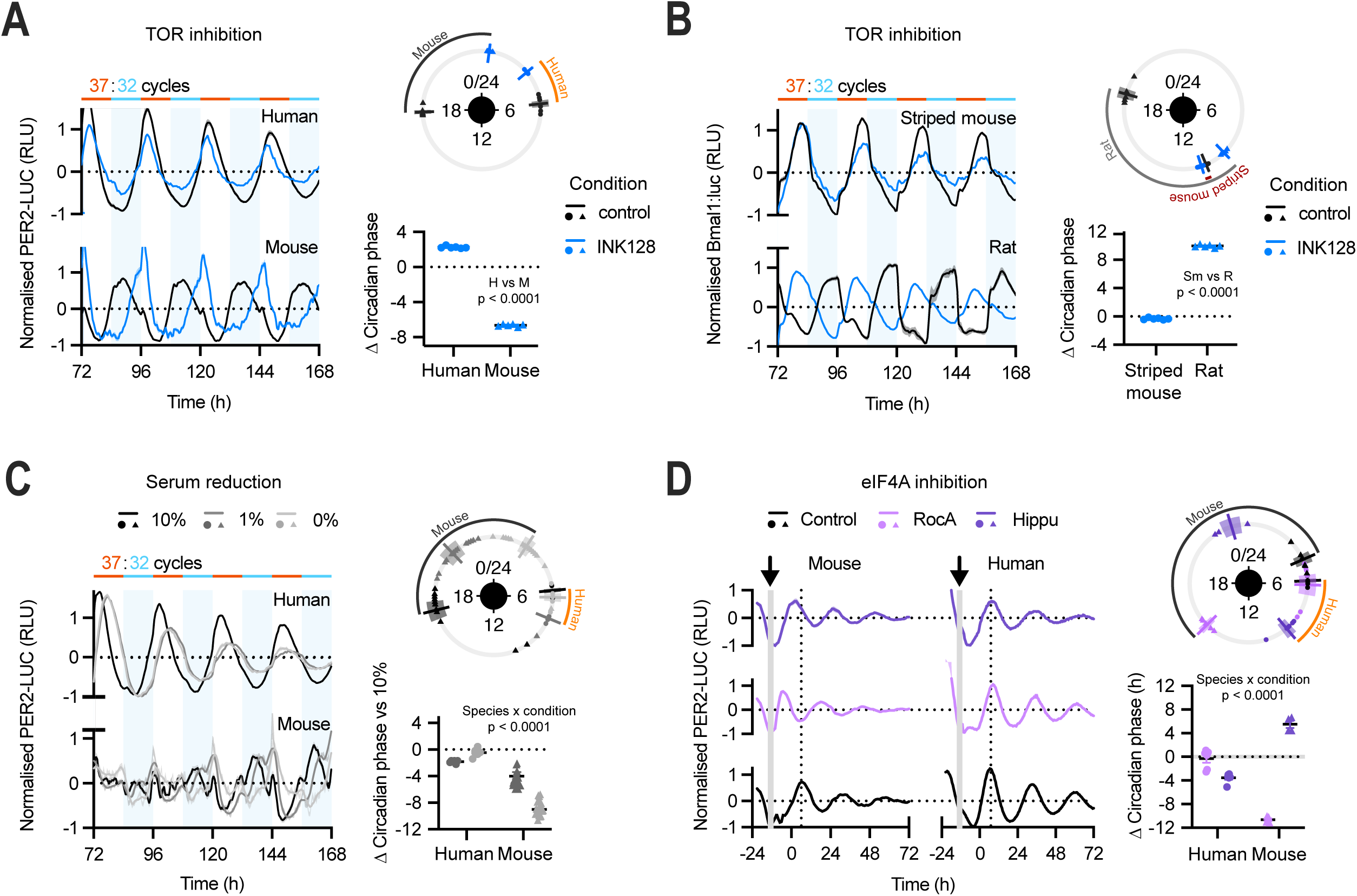
Manipulation of mTOR pathway activity alters phase of entrainment. **(A and B)** Human U2OS and mouse fibroblasts expressing PER2-LUC (A) and striped mouse and rat fibroblasts expressing *Bmal1:luc* (B) were entrained in 7 x 12h:12h 37°C:32°C temperature cycles in the presence of mTORC1/2 inhibitor (1 µM INK128) or control, then left to free-run at 37°C. Bioluminescence from PER2-LUC was recorded throughout, detrended and normalized to aid visualisation of circadian phase during cycling conditions. Circadian phase under control (black) and INK128 (blue) conditions is shown for human and striped mice (circles) and mouse and rat (triangles) as circle plots, and change in phase, relative to control, is shown below. Statistics: Phase_control_ vs phase_treatment_ two-way ANOVA followed by Šídák’s post-hoc test, n=4-6 each condition. **(C)** Human and mouse PER2-LUC fibroblasts, cultured in decreasing concentrations of serum, were entrained to 7 days of temperature cycles before transfer to constant conditions. Dashed line at the peak of 10% serum control is shown for illustration purposes. (Right) Circadian phase under control (black) and 1% serum (medium grey) and 0% serum (light grey) conditions is shown for human (circles) and mouse (triangles). Colored lines indicate human (orange) and mouse phases (grey). Change in phase, relative to 10% serum control, is shown below. Statistics: Two-way ANOVA followed by Šídák’s post-hoc test, n=6 (human) or 19-24 (mouse). **(D)** Human and mouse PER2-LUC cells, kept in constant conditions, were treated with inhibitors of the cap-binding complex that target eIF4A (rocaglamide, rocA; hippuristanol) at the trough of PER2-LUC at t=-12h, indicated by an arrow. Dashed line at the peak of DMSO control is shown for illustration purposes. (Right) Change in phase, relative to control, in mouse and human after treatment. Statistics: Two-way ANOVA followed by Šídák’s post-hoc test, n=4 each condition.

mTOR inhibition was not sufficient to make nocturnal cells completely phenocopy diurnal cells, and several other essential genes showed faster evolution in diurnal than nocturnal species including translational regulators (table S3). We therefore assessed how robust human *versus* mouse cellular circadian rhythms are to acute perturbation of translation rate by pharmacological attenuation of 5’-cap-dependent translational initiation, independently of mTOR or temperature. Circadian clocks drive, and are driven/synchronized, by daily cycles of protein synthesis (*28*, *111*– *114*), amplified *in vivo* by daily timing cues such as insulin/IGF-1 signaling linked with feed/fast cycles, which act *via* the translational machinery (*29*). When treated at the same circadian phase, very clear and significant differences were observed in the magnitude and direction of circadian phase shifts between mouse and human cells in response to direct inhibitors of eIF4A, rocaglamide A (RocA) and hippuristanol (Fig. 4D). Again, the cellular clock in mouse cells was much more sensitive than in human cells, consistent with the idea that natural selection has led to increased resistance to translational perturbation in diurnal mammals.

### mTOR regulation of circadian phase is maintained from cells to tissues

The function of the mTOR pathway as a cellular signaling nexus for translational regulation is conserved across eukaryotes and essential in mammals. Our results strongly suggest that differences in mTOR regulation and activity constitute a major element of the nocturnal-diurnal switch. If so, mTOR inhibition should render circadian clocks in mouse tissues more diurnal in their response to daily temperature cycles, both *ex vivo* and *in vivo*.

To test this, we subjected tissue explants from adult PER2-LUC mice to daily temperature cycles ± mTOR inhibition (Fig. 5A). As expected (*46*, *115*), high amplitude PER2-LUC oscillations were observed in neuroendocrine (pituitary) and non-neuronal (lung, adrenal) tissues, with PER2 consistently peaking around the warm-to-cold transition (Fig. 5B), as in mouse fibroblasts (Fig. 1E). Reduction of mTOR activity by INK128 resulted in a significant phase shift, delaying the PER2-LUC peak by 8-12h to near the cold-to-warm transition (Fig. 5B), such that they now resembled human cells rather than mouse cells *in vitro* (Fig. 1E).

**Fig. 5.**
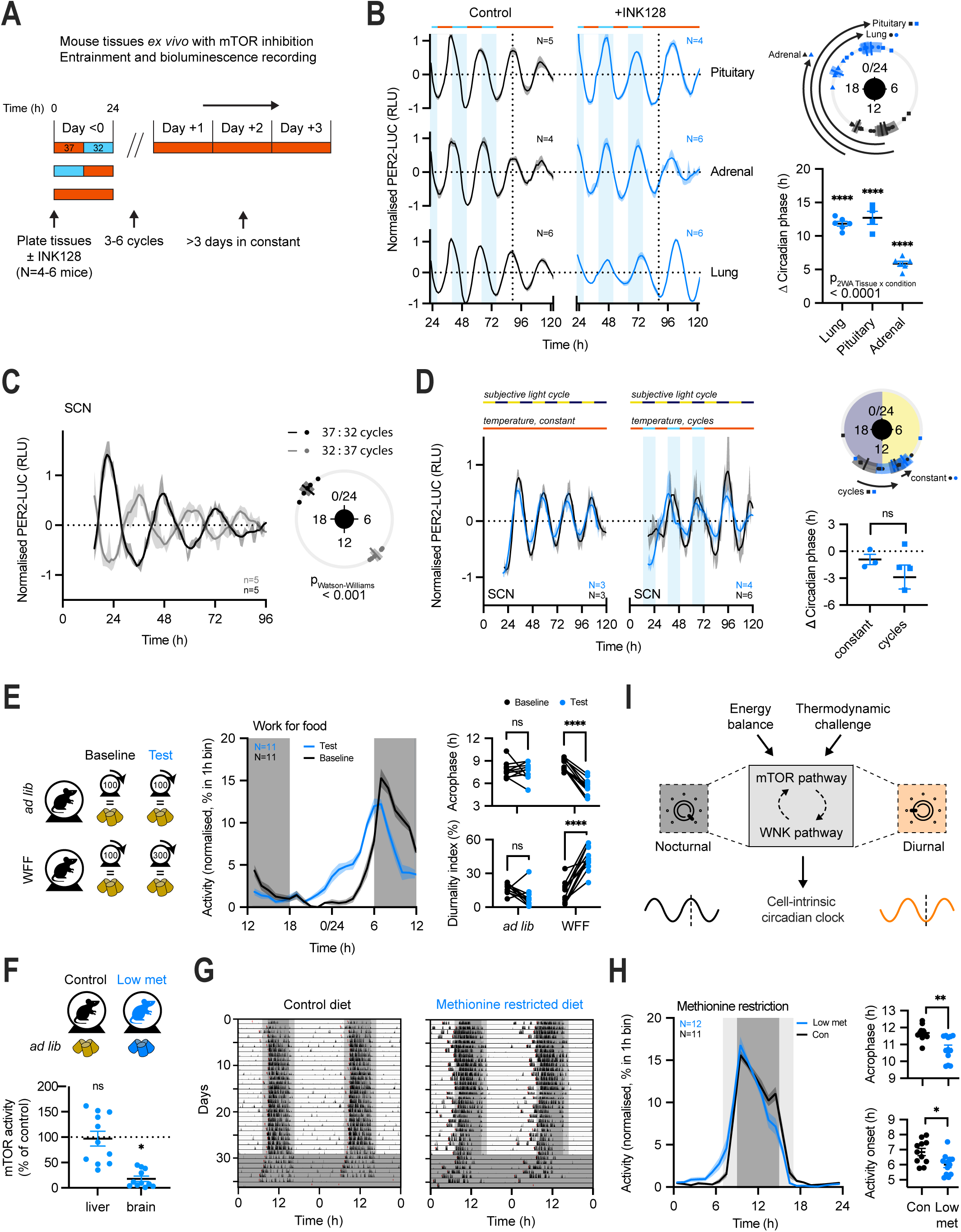
mTOR pathway activity underlies phase of entrainment in tissues and the nocturnal-diurnal switch *in vivo*. **(A)** Experiment schematic. Tissues were dissected from PER2-LUC mice and exposed to 3-6 temperature cycles coincident with or antiphasic to their previous activity patterns, or constant temperature. INK128 or control was added at the start of culture and luminescence from PER2-LUC was measured to compare circadian phase of entrainment. **(B)** PER2-LUC mouse pituitary (square), adrenal (triangle) or lung (circle) tissues were dissected from mice at the beginning of lights on (i.e. the start of the rest phase for nocturnal mice) at time = 0h. Tissue slices were cultured *ex vivo* in the presence of luciferin, treated with DMSO (black, left) or INK128 (blue, right), and exposed to cycles of temperature (12h 37°C: 12h 32°C) antiphasic to previous activity patterns. Days 2 and 3 of temperature cycles and 3 days of constant conditions are shown. (Right) Circadian phase of entrainment, calculated given relative to the final transition from 32°C to 37°C, for each tissue under each condition is shown, with arrows indicating the direction of the phase shift. Delta phase, relative to DMSO control. Statistics: TWA mixed-effects model, matched design. Interaction tissue x drug F(2, 24) = 4.28, p = 0.025; Šídák’s post-hoc test reported, n=5 each condition. **(C)** SCN tissue was dissected from PER2-LUC mice, sliced, and placed into *ex vivo* culture in the presence of luciferin. Slices were entrained for 7 days in antiphasic temperature cycles (12h: 12h 37 °C: 32°C, black line, or 32°C: 37°C, grey line) before release into constant 37 °C. (Right) Circadian phase of entrainment, relative to time = 0h. Statistics: Student’s t-test, unpaired, n=4-5. **(D)** SCN slices from PER2-LUC mice kept under LD cycles (subjective LD cycle shown) were kept at constant temperature (37 °C) or exposed for 3 days to temperature cycles (12h: 12h 37°C: 32°C) in antiphase to the LD cycle from which the mice were taken. *Ex vivo* culture started at 0h when slices were given either INK128 (blue) or DMSO (black), and bioluminescence was monitored throughout. PER2-LUC expression is plotted against circadian time, where 0h = the start of the subjective light period. Delta phase, relative to DMSO control. Student’s t-test, unpaired, n=3-5, not significant. **(E)** Experimental schematic for the Work for Food (WFF) paradigm as conducted by Hut et al. (*125*). Adult male mice were singly housed in cages equipped water *ad libitum* and running wheels under 12h: 12h light:dark cycles throughout. The number of food pellets given was controlled by the number of revolutions each mouse made on the running wheel. In the baseline portion of the experiment, one pellet was given per ∼100 revolutions. The number of revolutions per pellet was gradually increased in the WFF group until one pellet was given per ∼300 revolutions. In this way, a finely controlled organismal reduction in mTOR-dependent processes can be achieved (*128*). Running wheel activity and core body temperature from implanted temperature logger was recorded throughout. **(F)** (Left) Average proportion of total daily locomotor activity in 1h bins across the 24h day during the baseline and test period in the WFF group is shown. Activity in N=11 mice was averaged over 7 days of recording during each period of the experiment. (Right) Average acrophase of activity (top) and diurnality index (% of activity occurring in hours of full light, bottom) is shown for control (n=10) and WFF groups (n=11), with Holm-Šídák’s post-hoc test significance after Two-way ANOVA indicated. **(G to I)** WT mice (N=23, 12 male, 11 female) were kept in long day conditions (14h light: 4h twilight: 6h dark) for 1 weeks with *ad libitum* supply of food and water. After week 1, N=12 mice (6 male, 6 female) were switched onto an ad libitum supply of compositionally identical food with the exception that methionine content was reduced to 25% of control. N=11 mice (6 male, 5 female) were kept on *ad libitum* control diet containing 100% methionine. Mice were kept on diets for 4 weeks with continuous monitoring of wheel running activity. (G) mTOR activity in liver and brain taken from mice after methionine restriction for 4 weeks, vs mice on control diet as indicated by immunoblot. **(H)** Representative actogram of mice on control diet (left) and methionine restricted diet (right). **(I)** Average 24h activity was calculated for days 15-37 for all mice, binned into 1h bins and normalized. Light conditions are indicated relative to midday as t=0h with twilight (light grey) and darkness (dark grey) periods shaded. (Right) Average acrophase of activity (top) and diurnality index (% of activity occurring in hours of full light, bottom) is shown for control diet (con) and low methionine diet (low met) groups, with t-test (Welch’s correction) significance indicated. **(J)** Thermodynamic challenge and energy balance are integrated intracellularly by the mTOR pathway and WNK pathways, which feedback upon each other and regulate the cell-intrinsic circadian clock. Differential sensitivity of this feedback in nocturnal and diurnal cells (through multigenic modifications including WNK1, RRAGB and ZNF598), sets the cellular circadian clock response resulting in distinct circadian phases. Dial icon by *Colourcreatype* from Noun Project (CC BY 3.0).

The hypothalamic SCN of nocturnal mammals are remarkable for PER rhythms that are essentially opposite to almost all other tissues, in the same phase as SCN of diurnal mammals (Fig. 1A). This is consistent with the SCN’s conserved function in all mammals as a dedicated photic timekeeper, responsible for encoding and communicating anticipated photoperiod. Interneuronal coupling renders SCN PER rhythms more robust than other tissues and much more sensitive to photic cues than to systemic signals such as temperature (*29*, *46*, *116*). Adult SCN are sensitive to temperature, however, and explants stably entrain to the same daily temperature cycles employed throughout this study (Fig. 5C)(*116–118*). We found that after 7 daily temperature cycles, SCN entrained with a phase that was much later than other tissues, with PER2 peaking late in the cold phase (Fig. 5C) and at the end of the subjective day (Fig. 5D), reminiscent of the difference in circadian timing between SCN and other mouse tissues *in vivo*. Remarkably, the phase of SCN rhythms remained unaltered by mTOR inhibition (Fig. 5D). This mirrors the phenotype seen in cells and tissues from diurnal species, likely resulting from functional insensitivity to mTOR inhibition, which diminishes the responsiveness of the protein synthesis machinery to temperature changes. Indeed, the SCN is unaffected by abrupt changes in translation rate, which is conferred by network coupling (*112*, *119*). Since SCN activity in nocturnal mammals aligns with daytime, as for diurnal mammals (Fig 1A), mouse brain temperature rhythms would be expected to reinforce, rather than disrupt, the SCN’s established relationship with the light:dark cycle *in vivo*. Our findings support a model in which all mammalian SCN maintain an mTOR-insensitive representation of daytime, while the timing of behavior and physiology outside the SCN is governed by the interaction between cell-autonomous timekeeping and timing cues – such as temperature, osmolarity and growth factors – that regulate global and specific (PER) protein synthesis via mTOR.

### mTOR activity regulates nocturnal-to-diurnal behavioral switching

We have used a pharmacological inhibitor (INK128) that binds to the active site of mTOR (*120*) to demonstrate that, *in vitro*, modifying the basal activity of this pathway differentially alters cell-intrinsic responses in nocturnal vs diurnal mammals and thus mTOR pathway activity is implicated in the nocturnal-diurnal switch. However, diurnality/nocturnality is a behavioral phenotype in which timing of locomotor activity defines temporal niche classification. To truly demonstrate that mTOR pathway activity is implicated in the nocturnal-diurnal switch, we need to observe locomotor activity switching under organismal mTOR activity modification.

Under dietary-restricted conditions, such as those found in the wild, mTORC1 activity and protein synthesis is greatly reduced (*121–123*). Accordingly, mouse behavior becomes more diurnal than when fed *ad libitum* (*124*). We sought to replicate these observations in mice under laboratory conditions. Precise control of mice energy balance can be achieved using the Work for Food (WFF) paradigm (*125*, *126*), under which food is limited (Fig. 5E, fig. S10A) and mice lose significant body mass according to the negative energy balance imposed upon them (fig. S10B) (*127*, *128*). In these conditions, mTOR activity is significantly reduced in multiple brain and peripheral tissues (fig. S11). Compared with control conditions (food *ad libitum*) where mice are nocturnally active (fig. S10, C and D), under the negative energy balance conditions of WFF, mice apportion more of their activity to the daytime, like a diurnal mammal, (Fig. 5F, fig. S10, C to E, (*125*)), which is matched by advanced timing of core body temperature rhythms towards daylight hours (fig. S10, F-H).

WFF demonstrates that, without affecting the SCN ((*129*); fig. S11), it is possible for a nocturnal mouse to significantly alter locomotor activity timing while integrating the same environmental cues, manifesting diurnal behavior. This is consistent with the conserved role of daylight timing signaling in nocturnal and diurnal mammals (Fig. 1A). Differential gene expression analysis implicates a role of mTOR (fig. S11), though demonstrates that many other pathways are targeted by this extreme starvation treatment (*128*, *130*). To confirm that mTOR pathway activity is fundamental to the selection of locomotor activity timing, we targeted mTOR activity organismally via isocaloric modification of amino acid concentration in the diet. Unlike total caloric restriction, which acts largely independently of the mTOR pathway (*131*) and results in a self-imposed feed-fast cycle (*132*), amino acid restriction inhibits mTOR activity through amino acid sensing by the Rag-dependent signaling pathway (*105*, *133*, *134*). In cells, amino acid reduction altered cellular entrainment to temperature cycles and phenocopied pharmacological inhibition of mTOR (fig. S10I). Partial, brain-restricted, mTOR inhibition was achieved in mice fed *ad libitum* for four weeks with an isocaloric methionine restricted diet (Fig. 5G, fig. S10, J and K) with minimal weight loss (fig. S10L). Under these conditions, which permit reduction in mTOR activity without adverse consequences such as excessive weight loss which can confound measurement of locomotor activity, both the onset and peak of activity of mice on a methionine-restricted diet was significantly phase-advanced into the daylight hours relative to control (Fig. 5, H and I) with no change in locomotor period (fig. S10M). Taken together, these activity shifts in response to organismal modifications of mTOR activity are consistent with the cellular data, and support a molecular mechanism whereby the basal level of mTOR activity modulates the response to physiological entraining cues. Therefore, amongst several factors, nocturnal-to-diurnal switching involves convergent evolution for differential responsiveness of WNK and mTOR pathway signaling which can be recapitulated *in vitro* and *in vivo* (Fig. 5J).

## Discussion

Mammalian colonization of the daytime niche accelerated when its previous occupants, the dinosaurs, became extinct (*1*). Subsequently, mammals came to occupy all temporal niches, frequently switching between them as life history and environment dictates (*5*, *135*). The specific mechanism that permits this switch between nocturnality and diurnality was previously unknown. We investigated an apparent cell-intrinsic inversion of the molecular circadian clockwork to entrainment cues that alter intracellular water thermodynamics – temperature and osmolarity. This largely arises from differences in the basal activity and sensitivity of the mTOR pathway, with downstream consequences on protein synthesis. Members of this pathway have evolved more rapidly in diurnal compared to nocturnal mammals, and modulation of mTOR activity in cultured cells, tissues or *in vivo* is able to recapitulate the switch from nocturnal to diurnal circadian timing.

These critical differences in the cellular response to temperature mirror recent findings in developmental biology, where mammalian species show marked differences in global biochemical reaction rates which correlate with developmental tempo (*57–59*). Our analogous discovery of significant differences in global phosphorylation and protein synthesis between mice and humans led to mTOR kinase as a plausible and key component of a different phenomenon: the nocturnal-diurnal switch. We note that mTOR complexes 1 and 2 have several substrate effectors and are regulated by multiple different cell signaling systems (*136*). As part of the large and interlinked PI3K-AKT-mTOR pathway, mTOR regulates and is regulated by cellular crowding *via* WNK1 and the circadian response to osmolarity, amongst myriad other things (*97*, *107*, *137*, *138*). Therefore, we do not discount that differential temperature sensitivity of other cellular kinases, phosphatases, and signaling mechanisms, acting upstream, downstream or in parallel with mTOR, may also contribute to temporal niche switching. Moreover, that the re-organization of physiology under reduced mTOR activity via amino acid restriction or WFF requires several weeks and does not recapitulate diurnal behaviour (*125*, *130*, *139*). Therefore, we also do not discount roles for hypothalamic neuroplasticity, melatonin signal inversion or direct photic modulation of locomotor activity in temporal niche selection (*13*, *140–142*).

Ultimately though, any switching mechanism that arose evolutionarily must have a genetic basis. We demonstrate this through a genome-wide comparison of diurnal and nocturnal mammals, which provides complimentary genetic evidence for the importance of mTOR activity with key proteins, RRAGB and WNK1, having faster evolutionary rates in diurnal *versus* nocturnal mammals. We consider genetic mechanisms of diurnality may be broadly dispersed and polygenic, and to this end we have evidence for faster evolutionary rates in olfactory pathway genes (fig. S9) and phototransduction genes (*143*). Future work might be directed towards identifying the detailed molecular and structural differences between diurnal and nocturnal mammals in mTOR pathway components and its regulators.

At the whole organism level, our findings agree with the circadian thermo-energetics (CTE) hypothesis for conditional niche-switching in several different mammals (*4*). CTE states that nocturnal activity patterns for homeothermic mammals are more costly than diurnal patterns, since nocturnal animals have higher energy requirements to mitigate the greater heat loss of being active during the (cold) night (*129*, *144*, *145*). Diurnality arises as an energy saving measure when food availability is scarce, which outcompetes the extra predation pressure of being active by day (*146*, *147*). At the cellular level, these results support the bioenergetic hypothesis for circadian and other biological rhythms (*148–152*), where oscillations primarily function to minimize the high cost of maintaining protein homeostasis. In this context, the increased resistance to translational perturbation in cells from diurnal mammals is thus an energy saving measure, and will diminish the cellular challenge of conflicting timing cues. It would be interesting to investigate whether birds – which independently evolved diurnality, homeothermy and have a higher basal core body temperature than mammals (*153*), as well as marked heat stress and specialized thermoregulation during flight (*154*) – use the same mechanism.

Overall, our findings illustrate marked species differences in the cellular environment and global pathway activity which influences circadian phase in cells, tissues and *in vivo*. Our findings add to a growing literature demonstrating species-specific differences in molecular activity which map to a cellular or external phenotype (*57–60*, *155–157*). It is striking that many of these findings involve global regulation of protein turnover and the mTOR pathway, and integrate metabolic status with functional output (*157*, *158*).

## Acknowledgements

We thank Ken Wright for intellectual contributions in the genesis of the project; Rajesh Narasimamurthy and David Virshup for pilot experiments; Michael Hastings, Robert Lucas and Roger Barker for resources and discussion; Tim Eppley for valuable discussions on activity patterns in Lemurs; Joana Frankel for general discussion on diurnal models; Jo Menzies for discussion on luciferase kinetics; and Aymen al-Rawi for valuable contributions to proteomic analysis. We thank all past and current members of O’Neill lab and David Bechtold, for their valuable feedback. We thank biomedical services group staff at Medical Research Council (MRC) Ares facility and LMB facilities for assistance. This project is supported through a research collaboration between AstraZeneca UK Limited and the Medical Research Council, reference BSF38. At the time of writing, Nina Rzechorzek was undertaking an AZ-MRC Industry Partnership for Academic Clinicians, partly funded by AstraZeneca and the Medical Research Council (MC_EX_MR/Y013018/1).

For the purpose of open access, the MRC Laboratory of Molecular Biology applies a CC BY public copyright licence to any Author Accepted Manuscript version arising.

## Funding

Medical Research Council MC_UP_1201/4 (JSO)

Medical Research Council MR/S022023/1 and MC_EX_MR/S022023/1 (NMR)

Wellcome Trust Investigator Award to Robert Lucas, University of Manchester 210684/Z/18/Z (RR)

Royal Society - Wellcome Sir Henry Dale Fellowship 208790/Z/17/Z and UKRI Future Leaders Fellowship MR/Y017552/1 (RSE)

## Author contributions

Conceptualization: ADB, JSO, NMR

Formal Analysis: ADB, NMR, MJC, SYP-C, VP, RAH

Funding acquisition: NMR, JSO

Investigation: ADB, MJC, NMR, AZ, AM, CE, NJS, VP, NRJ, SYP-C, RAH

Methodology: ADB, JSO

Project administration: ADB, JSO

Resources: NJS, RR, MAL, MFB, SVF, ZV, JP, RSE

Software: ADB Supervision: JSO

Visualization: ADB

Writing – original draft: ADB

Writing – review & editing: ADB, JSO, NMR, AM, RSE, MAL, MJC, AZ, RAH, NJS

## Data and materials availability

All data are available in the main text and supplementary material. Code are available at https://github.com/andrewbeale. Materials are available upon request to JSO and are subject to materials transfer agreements.

## Supplementary Materials

Materials and Methods Figs. S1 to S11

Tables S1 to S3 References (*161–266*)

## References and Notes

1. M. P. Gerkema, W. I. L. Davies, R. G. Foster, M. Menaker, R. A. Hut, The nocturnal bottleneck and the evolution of activity patterns in mammals. Proc Royal Soc B Biological Sci 280, 20130508 (2013).

2. R. Refinetti, The diversity of temporal niches in mammals. Biol Rhythm Res 39, 173–192 (2008).

3. R. Maor, T. Dayan, H. Ferguson-Gow, K. E. Jones, Temporal niche expansion in mammals from a nocturnal ancestor after dinosaur extinction. Nat Ecol Evol 1, 1889–1895 (2017).

4. R. A. Hut, N. Kronfeld-Schor, V. van der Vinne, H. De la Iglesia, “Chapter 17. In search of a temporal niche: Environmental factors” in Progress in Brain Research, A. Kalsbeek, M. Merrow, T. Roenneberg, R. G. Foster, Eds. (Elsevier, 2012)vol. 199 of Progress in Brain Research, pp. 281–304.

5. D. R. van der Veen, S. J. Riede, P. D. Heideman, M. Hau, V. van der Vinne, R. A. Hut, Flexible clock systems: adjusting the temporal programme. Philosophical Transactions Royal Soc B Biological Sci 372, 20160254 (2017).

6. M. dos Reis, J. Inoue, M. Hasegawa, R. J. Asher, P. C. J. Donoghue, Z. Yang, Phylogenomic datasets provide both precision and accuracy in estimating the timescale of placental mammal phylogeny. Proc Royal Soc B Biological Sci 279, 3491–3500 (2012).

7. C. L. Partch, C. B. Green, J. S. Takahashi, Molecular architecture of the mammalian circadian clock. Trends Cell Biol 24, 90–99 (2014).

8. D. C. Wong, J. S. O’Neill, Non-transcriptional processes in circadian rhythm generation. Curr Opin Physiology 5, 117–132 (2018).

9. R. Chen, A. Schirmer, Y. Lee, H. Lee, V. Kumar, S.-H. Yoo, J. S. Takahashi, C. Lee, Rhythmic PER Abundance Defines a Critical Nodal Point for Negative Feedback within the Circadian Clock Mechanism. Mol Cell 36, 417–430 (2009).

10. W. J. Schwartz, S. M. Reppert, S. M. Eagan, M. C. Moore-Ede, In vivo metabolic activity of the suprachiasmatic nuclei: a comparative study. Brain Res 274, 184–187 (1983).

11. A. A. Nunez, A. Bult, T. L. McElhinny, L. Smale, Daily Rhythms of Fos Expression in Hypothalamic Targets of the Suprachiasmatic Nucleus in Diurnal and Nocturnal Rodents. J Biol Rhythm 14, 300–306 (1999).

12. D. M. Berson, F. A. Dunn, M. Takao, Phototransduction by Retinal Ganglion Cells That Set the Circadian Clock. Science 295, 1070–1073 (2002).

13. B. Bano-Otalora, M. J. Moye, T. Brown, R. J. Lucas, C. O. Diekman, M. D. Belle, Daily electrical activity in the master circadian clock of a diurnal mammal. Elife 10, e68179 (2021).

14. A. P. Patton, M. H. Hastings, The suprachiasmatic nucleus. Curr Biology Cb 28, R816–R822 (2018).

15. E. Challet, Minireview: Entrainment of the suprachiasmatic clockwork in diurnal and nocturnal mammals. Endocrinology 148, 5648–55 (2007).

16. N. Inagaki, S. Honma, D. Ono, Y. Tanahashi, K. Honma, Separate oscillating cell groups in mouse suprachiasmatic nucleus couple photoperiodically to the onset and end of daily activity. Proc National Acad Sci 104, 7664–7669 (2007).

17. R. Hut, E. V. der Zee, K. Jansen, M. Gerkema, S. Daan, Gradual reappearance of post-hibernation circadian rhythmicity correlates with numbers of vasopressin-containing neurons in the suprachiasmatic nuclei of European ground squirrels. J Comp Physiol B 172, 59–70 (2002).

18. L. S. Mure, H. D. Le, G. Benegiamo, M. W. Chang, L. Rios, N. Jillani, M. Ngotho, T. Kariuki, O. Dkhissi-Benyahya, H. M. Cooper, S. Panda, Diurnal transcriptome atlas of a primate across major neural and peripheral tissues. Science 359 (2018).

19. R. Cohen, N. Kronfeld-Schor, C. Ramanathan, A. Baumgras, L. Smale, The Substructure of the Suprachiasmatic Nucleus: Similarities between Nocturnal and Diurnal Spiny Mice. Brain Behav. Evol. 75, 9–22 (2010).

20. C. Ramanathan, A. Stowie, L. Smale, A. A. Nunez, Phase preference for the display of activity is associated with the phase of extra-suprachiasmatic nucleus oscillators within and between species. Neuroscience 170, 758–772 (2010).

21. C. M. Lambert, D. R. Weaver, Peripheral Gene Expression Rhythms in a Diurnal Rodent. J Biol Rhythm 21, 77–79 (2006).

22. P. K. Jha, E. Challet, A. Kalsbeek, Circadian rhythms in glucose and lipid metabolism in nocturnal and diurnal mammals. Mol Cell Endocrinol 418, 74–88 (2015).

23. G. D. M. Potter, D. J. Skene, J. Arendt, J. E. Cade, P. J. Grant, L. J. Hardie, Circadian Rhythm and Sleep Disruption: Causes, Metabolic Consequences, and Countermeasures. Endocr Rev 37, 584–608 (2016).

24. C. Dibner, U. Schibler, Circadian timing of metabolism in animal models and humans. J Intern Med 277, 513–527 (2015).

25. J. Bass, J. S. Takahashi, Circadian Integration of Metabolism and Energetics. Science 330, 1349–1354 (2010).

26. C. Scheiermann, J. Gibbs, L. Ince, A. Loudon, Clocking in to immunity. Nat Rev Immunol 18, 423–437 (2018).

27. M. D’Alessandro, S. Beesley, J. K. Kim, R. Chen, E. Abich, W. Cheng, P. Yi, J. S. Takahashi, C. Lee, A tunable artificial circadian clock in clock-defective mice. Nat Commun 6, 8587 (2015).

28. K. A. Feeney, L. L. Hansen, M. Putker, C. Olivares-Yañez, J. Day, L. J. Eades, L. F. Larrondo, N. P. Hoyle, J. S. O’Neill, G. van Ooijen, Daily magnesium fluxes regulate cellular timekeeping and energy balance. Nature 532, 375–379 (2016).

29. P. Crosby, R. Hamnett, M. Putker, N. P. Hoyle, M. Reed, C. J. Karam, E. S. Maywood, A. Stangherlin, J. E. Chesham, E. A. Hayter, L. Rosenbrier-Ribeiro, P. Newham, H. Clevers, D. A. Bechtold, J. S. O’Neill, Insulin/IGF-1 Drives PERIOD Synthesis to Entrain Circadian Rhythms with Feeding Time. Cell 177, 896–909.e20 (2019).

30. A. R. M. Buijs, J. Wortel, J. J. V. Heerikhuize, M. G. P. Feenstra, G. J. T. Horst, H. J. Romijn, Kalsbeek, Anatomical and functional demonstration of a multisynaptic suprachiasmatic nucleus adrenal (cortex) pathway. Eur J Neurosci 11, 1535–1544 (1999).

31. A. Balsalobre, S. A. Brown, L. Marcacci, F. Tronche, C. Kellendonk, H. M. Reichardt, G. Schütz, U. Schibler, Resetting of Circadian Time in Peripheral Tissues by Glucocorticoid Signaling. Science 289, 2344–2347 (2000).

32. S. M. Ota, X. Kong, R. Hut, D. Suchecki, P. Meerlo, The impact of stress and stress hormones on endogenous clocks and circadian rhythms. Frontiers Neuroendocrinol. 63, 100931 (2021).

33. X. Kong, S. M. Ota, D. Suchecki, A. Lan, A. I. Peereboom, R. A. Hut, P. Meerlo, Chronic Social Defeat Stress Shifts Peripheral Circadian Clocks in Male Mice in a Tissue-Specific and Time-of-Day Dependent Fashion. J. Biological Rhythm. 37, 164–176 (2022).

34. X. Kong, M. Luxwolda, R. A. Hut, P. Meerlo, Adrenalectomy prevents the effects of social defeat stress on PER2 rhythms in some peripheral tissues in male mice. Horm. Behav. 150, 105326 (2023).

35. A. Mihut, J. S. O’Neill, C. L. Partch, P. Crosby, PERspectives on circadian cell biology. Philos. Trans. B 380, 20230483 (2025).

36. T. Roenneberg, S. Daan, M. Merrow, The Art of Entrainment. J Biol Rhythm 18, 183–194 (2003).

37. C. H. Johnson, J. A. Elliott, R. Foster, Entrainment of Circadian Programs. Chronobiol Int 20, 741–774 (2003).

38. F. Damiola, N. L. Minh, N. Preitner, B. Kornmann, F. Fleury-Olela, U. Schibler, Restricted feeding uncouples circadian oscillators in peripheral tissues from the central pacemaker in the suprachiasmatic nucleus. Gene Dev 14, 2950–2961 (2000).

39. K.-A. Stokkan, S. Yamazaki, H. Tei, Y. Sakaki, M. Menaker, Entrainment of the Circadian Clock in the Liver by Feeding. Science 291, 490–493 (2001).

40. Y. Isojima, M. Nakajima, H. Ukai, H. Fujishima, R. G. Yamada, K. Masumoto, R. Kiuchi, M. Ishida, M. Ukai-Tadenuma, Y. Minami, R. Kito, K. Nakao, W. Kishimoto, S.-H. Yoo, K. Shimomura, T. Takao, A. Takano, T. Kojima, K. Nagai, Y. Sakaki, J. S. Takahashi, H. R. Ueda, CKIε/δ-dependent phosphorylation is a temperature-insensitive, period-determining process in the mammalian circadian clock. Proc National Acad Sci 106, 15744–15749 (2009).

41. R. Refinetti, M. Menaker, The circadian rhythm of body temperature. Physiol Behav 51, 613–637 (1992).

42. R. Refinetti, M. Menaker, The circadian rhythm of body temperature of normal and tau-mutant golden hamsters. J Therm Biol 17, 129–133 (1992).

43. K. Krauchi, A. Wirz-Justice, Circadian rhythm of heat production, heart rate, and skin and core temperature under unmasking conditions in men. Am J Physiology-regulatory Integr Comp Physiology 267, R819–R829 (1994).

44. S. A. Brown, G. Zumbrunn, F. Fleury-Olela, N. Preitner, U. Schibler, Rhythms of mammalian body temperature can sustain peripheral circadian clocks. Curr Biol 12, 1574--1583 (2002).

45. L. M. Prolo, J. S. Takahashi, E. D. Herzog, Circadian Rhythm Generation and Entrainment in Astrocytes. J Neurosci 25, 404–408 (2005).

46. E. D. Buhr, S.-H. Yoo, J. S. Takahashi, Temperature as a Universal Resetting Cue for Mammalian Circadian Oscillators. Science 330, 379–385 (2010).

47. C. Saini, J. Morf, M. Stratmann, P. Gos, U. Schibler, Simulated body temperature rhythms reveal the phase-shifting behavior and plasticity of mammalian circadian oscillators. Gene Dev 26, 567--580 (2012).

48. A. D. Beale, E. A. Hayter, P. Crosby, U. K. Valekunja, R. S. Edgar, J. E. Chesham, E. S. Maywood, F. H. Labeed, A. B. Reddy, K. P. Wright, K. S. Lilley, D. A. Bechtold, M. H. Hastings, J. S. O’Neill, Mechanisms and physiological function of daily haemoglobin oxidation rhythms in red blood cells. EMBO J., e114164 (2023).

49. J. Morf, G. Rey, K. Schneider, M. Stratmann, J. Fujita, F. Naef, U. Schibler, Cold-inducible RNA-binding protein modulates circadian gene expression posttranscriptionally. Science 338, 379--383 (2012).

50. Y. Liu, W. Hu, Y. Murakawa, J. Yin, G. Wang, M. Landthaler, J. Yan, Cold-induced RNA-binding proteins regulate circadian gene expression by controlling alternative polyadenylation. Sci Rep-uk 3, 1--11 (2013).

51. T. Haltenhof, A. Kotte, F. D. Bortoli, S. Schiefer, S. Meinke, A.-K. Emmerichs, K. K. Petermann, B. Timmermann, P. Imhof, A. Franz, B. Loll, M. C. Wahl, M. Preußner, F. Heyd, A Conserved Kinase-Based Body-Temperature Sensor Globally Controls Alternative Splicing and Gene Expression. Mol Cell 78, 57–69.e4 (2020).

52. T. Miyake, Y. Inoue, X. Shao, T. Seta, Y. Aoki, K. T. N. Pham, Y. Shichino, J. Sasaki, T. Sasaki, M. Ikawa, Y. Yamaguchi, H. Okamura, S. Iwasaki, M. Doi, Minimal upstream open reading frame of Per2 mediates phase fitness of the circadian clock to day/night physiological body temperature rhythm. Cell Reports, 112157 (2023).

53. B. Kornmann, O. Schaad, H. Bujard, J. S. Takahashi, U. Schibler, System-driven and oscillator-dependent circadian transcription in mice with a conditionally active liver clock. Plos Biol 5, 0179--0189 (2007).

54. H. Reinke, C. Saini, F. Fleury-Olela, C. Dibner, I. J. Benjamin, U. Schibler, Differential display of DNA-binding proteins reveals heat-shock factor 1 as a circadian transcription factor. Gene Dev 22, 331--345 (2008).

55. I. Gotic, S. Omidi, F. Fleury-Olela, N. Molina, F. Naef, U. Schibler, Temperature regulates splicing efficiency of the cold-inducible RNA-binding protein gene Cirbp. Gene Dev 30, 2005– 2017 (2016).

56. M. Ebisuya, J. Briscoe, What does time mean in development? Dev Camb Engl 145, dev164368 (2018).

57. M. Matsuda, H. Hayashi, J. Garcia-Ojalvo, K. Yoshioka-Kobayashi, R. Kageyama, Y. Yamanaka, M. Ikeya, J. Toguchida, C. Alev, M. Ebisuya, Species-specific segmentation clock periods are due to differential biochemical reaction speeds. Science 369, 1450–1455 (2020).

58. T. Rayon, D. Stamataki, R. Perez-Carrasco, L. Garcia-Perez, C. Barrington, M. Melchionda, K. Exelby, J. Lazaro, V. L. J. Tybulewicz, E. M. C. Fisher, J. Briscoe, Species-specific pace of development is associated with differences in protein stability. Science 369 (2020).

59. J. Lázaro, M. Costanzo, M. Sanaki-Matsumiya, C. Girardot, M. Hayashi, K. Hayashi, S. Diecke, T. B. Hildebrandt, G. Lazzari, J. Wu, S. Petkov, R. Behr, V. Trivedi, M. Matsuda, M. Ebisuya, A stem cell zoo uncovers intracellular scaling of developmental tempo across mammals. Cell Stem Cell 30, 938–949.e7 (2023).

60. M. Matsuda, J. Lázaro, M. Ebisuya, Metabolic activities are selective modulators for individual segmentation clock processes. Nat. Commun. 16, 845 (2025).

61. J. S. O’Neill, A. B. Reddy, Circadian clocks in human red blood cells. Nature 469, 498–503 (2011).

62. N. M. Rzechorzek, M. J. Thrippleton, F. M. Chappell, G. Mair, A. Ercole, M. Cabeleira, The CENTER-TBI High Resolution ICU (HR ICU) Sub-Study Participants and Investigators, J. Rhodes, I. Marshall, J. S. O’Neill, A daily temperature rhythm in the human brain predicts survival after brain injury. Brain 145, 2031–2048 (2022).

63. L. Santini, D. Rojas, G. Donati, Evolving through day and night: origin and diversification of activity pattern in modern primates. Behav Ecol 26, 789–796 (2015).

64. V. E. Mendel, G. V. Raghavan, A study of diurnal temperature patterns in sheep. J Physiology 174, 206–216 (1964).

65. D. M. Schumann, H. M. Cooper, M. D. Hofmeyr, N. C. Bennett, Circadian rhythm of locomotor activity in the four-striped field mouse, Rhabdomys pumilio: A diurnal African rodent. Physiol. Behav. 85, 231–239 (2005).

66. G. Donati, L. Santini, J. Razafindramanana, L. Boitani, S. Borgognini-Tarli, (Un-)expected nocturnal activity in “Diurnal” Lemur catta supports cathemerality as one of the key adaptations of the lemurid radiation. Am J Phys Anthropol 150, 99–106 (2013).

67. S.-H. Yoo, S. Yamazaki, P. L. Lowrey, K. Shimomura, C. H. Ko, E. D. Buhr, S. M. Siepka, H.-K. Hong, W. J. Oh, O. J. Yoo, M. Menaker, J. S. Takahashi, PERIOD2::LUCIFERASE real-time reporting of circadian dynamics reveals persistent circadian oscillations in mouse peripheral tissues. P Natl Acad Sci Usa 101, 5339–5346 (2004).

68. J. Park, K. Lee, H. Kim, H. Shin, C. Lee, Endogenous circadian reporters reveal functional differences of PERIOD paralogs and the significance of PERIOD:CK1 stable interaction. P Natl Acad Sci Usa 120, e2212255120 (2023).

69. K. A. Feeney, M. Putker, M. Brancaccio, J. S. O’Neill, In-depth Characterization of Firefly Luciferase as a Reporter of Circadian Gene Expression in Mammalian Cells. J Biol Rhythm 31, 540–550 (2016).

70. G. Manella, D. Aizik, R. Aviram, M. Golik, G. Asher, Circa-SCOPE: high-throughput live single-cell imaging method for analysis of circadian clock resetting. Nat Commun 12, 5903 (2021).

71. G. Manella, N. Bolshette, M. Golik, G. Asher, Input integration by the circadian clock exhibits nonadditivity and fold-change detection. Proc National Acad Sci 119 (2022).

72. O. Gandelman, I. Allue, K. Bowers, P. Cobbold, Cytoplasmic factors that affect the intensity and stability of bioluminescence from firefly luciferase in living mammalian cells. J. Biolumin. Chemilumin. 9, 363–371 (1994).

73. B. R. Branchini, R. A. Magyar, K. M. Marcantonio, K. J. Newberry, J. G. Stroh, L. K. Hinz, M. H. Murtiashaw, Identification of a Firefly Luciferase Active Site Peptide Using a Benzophenone-based Photooxidation Reagent*. J. Biol. Chem. 272, 19359–19364 (1997).

74. M. Doi, H. Shimatani, Y. Atobe, I. Murai, H. Hayashi, Y. Takahashi, J.-M. Fustin, Y. Yamaguchi, H. Kiyonari, N. Koike, K. Yagita, C. Lee, M. Abe, K. Sakimura, H. Okamura, Non-coding cis-element of Period2 is essential for maintaining organismal circadian behaviour and body temperature rhythmicity. Nat Commun 10, 2563 (2019).

75. C.-H. Yu, Y. Dang, Z. Zhou, C. Wu, F. Zhao, M. S. Sachs, Y. Liu, Codon Usage Influences the Local Rate of Translation Elongation to Regulate Co-translational Protein Folding. Mol Cell 59, 744–754 (2015).

76. V. Presnyak, N. Alhusaini, Y.-H. Chen, S. Martin, N. Morris, N. Kline, S. Olson, D. Weinberg, K. E. Baker, B. R. Graveley, J. Coller, Codon Optimality Is a Major Determinant of mRNA Stability. Cell 160, 1111–1124 (2015).

77. Q. Wu, S. G. Medina, G. Kushawah, M. L. DeVore, L. A. Castellano, J. M. Hand, M. Wright, A. A. Bazzini, Translation affects mRNA stability in a codon-dependent manner in human cells. Elife 8, e45396 (2019).

78. Y. Liu, Q. Yang, F. Zhao, Synonymous but not Silent: The Codon Usage Code for Gene Expression and Protein Folding. Annu Rev Biochem 90, 1–27 (2021).

79. M. Verma, J. Choi, K. A. Cottrell, Z. Lavagnino, E. N. Thomas, S. Pavlovic-Djuranovic, P. Szczesny, D. W. Piston, H. S. Zaher, J. D. Puglisi, S. Djuranovic, A short translational ramp determines the efficiency of protein synthesis. Nat Commun 10, 5774 (2019).

80. J. Fu, K. A. Murphy, M. Zhou, Y. H. Li, V. H. Lam, C. A. Tabuloc, J. C. Chiu, Y. Liu, Codon usage affects the structure and function of the Drosophila circadian clock protein PERIOD. Gene Dev 30, 1761–1775 (2016).

81. M. Zhou, J. Guo, J. Cha, M. Chae, S. Chen, J. M. Barral, M. S. Sachs, Y. Liu, Non-optimal codon usage affects expression, structure and function of clock protein FRQ. Nature 495, 111– 115 (2013).

82. N. Craig, Effect of reduced temperatures on protein synthesis in mouse L cells. Cell 4, 329– 335 (1975).

83. N. Craig, Regulation of translation in rabbit reticulocytes and mouse L-cells; comparison of the effects of temperature. J. Cell. Physiol. 87, 157–166 (1976).

84. C. S. Pittendrigh, On temperature independence in the clock system controlling emergence time in Drosophila. Proc National Acad Sci 40, 1018–1029 (1954).

85. R. Narasimamurthy, D. M. Virshup, Molecular Mechanisms Regulating Temperature Compensation of the Circadian Clock. Front Neurol 8, 1--5 (2017).

86. C. G. Proud, Phosphorylation and Signal Transduction Pathways in Translational Control. Csh Perspect Biol 11, a033050 (2019).

87. N. R. James, J. S. O’Neill, Circadian Control of Protein Synthesis. BioEssays, doi: 10.1002/bies.202300158 (2024).

88. X. Liu, P. C. C. Liu, N. Santoro, D. J. Thiele, Conservation of a stress response: human heat shock transcription factors functionally substitute for yeast HSF. The EMBO J. 16, 6466–6477 (1997).

89. M. E. LLeonart, A new generation of proto-oncogenes: Cold-inducible RNA binding proteins. Biochimica et Biophys. Acta BBA - Rev. Cancer 1805, 43–52 (2010).

90. M. B. Al-Fageeh, C. M. Smales, Control and regulation of the cellular responses to cold shock: the responses in yeast and mammalian systems. Biochem. J. 397, 247–259 (2006).

91. H. L. Rust, P. R. Thompson, Kinase Consensus Sequences: A Breeding Ground for Crosstalk. Acs Chem Biol 6, 881–892 (2011).

92. J. L. Johnson, T. M. Yaron, E. M. Huntsman, A. Kerelsky, J. Song, A. Regev, T.-Y. Lin, K. Liberatore, D. M. Cizin, B. M. Cohen, N. Vasan, Y. Ma, K. Krismer, J. T. Robles, B. van de Kooij, A. E. van Vlimmeren, N. Andrée-Busch, N. F. Käufer, M. V. Dorovkov, A. G. Ryazanov, Y. Takagi, E. R. Kastenhuber, M. D. Goncalves, B. D. Hopkins, O. Elemento, D. J. Taatjes, A. Maucuer, A. Yamashita, A. Degterev, M. Uduman, J. Lu, S. D. Landry, B. Zhang, I. Cossentino, R. Linding, J. Blenis, P. V. Hornbeck, B. E. Turk, M. B. Yaffe, L. C. Cantley, An atlas of substrate specificities for the human serine/threonine kinome. Nature, 1–8 (2023).

93. C. C. Dibble, B. D. Manning, Signal integration by mTORC1 coordinates nutrient input with biosynthetic output. Nat Cell Biol 15, 555–564 (2013).

94. C. C. Thoreen, L. Chantranupong, H. R. Keys, T. Wang, N. S. Gray, D. M. Sabatini, A unifying model for mTORC1-mediated regulation of mRNA translation. Nature 485, 109–113 (2012).

95. A. J. Valvezan, B. D. Manning, Molecular logic of mTORC1 signalling as a metabolic rheostat. Nat. Metab. 1, 321–333 (2019).

96. L. J. Holt, M. Delarue, Macromolecular crowding: Sensing without a sensor. Curr. Opin. Cell Biol. 85, 102269 (2023).

97. M. Delarue, G. P. Brittingham, S. Pfeffer, I. V. Surovtsev, S. Pinglay, K. J. Kennedy, M. Schaffer, J. I. Gutierrez, D. Sang, G. Poterewicz, J. K. Chung, J. M. Plitzko, J. T. Groves, C. Jacobs-Wagner, B. D. Engel, L. J. Holt, mTORC1 Controls Phase Separation and the Biophysical Properties of the Cytoplasm by Tuning Crowding. Cell 174, 338–349.e20 (2018).

98. Y. Xie, T. Shu, T. Liu, M.-C. Spindler, J. Mahamid, G. M. Hocky, D. Gresham, L. J. Holt, Polysome collapse and RNA condensation fluidize the cytoplasm. Mol. Cell 84, 2698–2716.e9 (2024).

99. J. Kim, H. Kim, K. Hwang, J. S. Chang, K. Park, S. Cha, I. D. Kong, WNK1 kinase is essential for insulin-stimulated GLUT4 trafficking in skeletal muscle. FEBS Open Bio 8, 1866– 1874 (2018).

100. A. Stangherlin, J. L. Watson, D. C. S. Wong, S. Barbiero, A. Zeng, E. Seinkmane, S. P. Chew, A. D. Beale, E. A. Hayter, A. Guna, A. J. Inglis, M. Putker, E. Bartolami, S. Matile, N. Lequeux, T. Pons, J. Day, G. van Ooijen, R. M. Voorhees, D. A. Bechtold, E. Derivery, R. S. Edgar, P. Newham, J. S. O’Neill, Compensatory ion transport buffers daily protein rhythms to regulate osmotic balance and cellular physiology. Nat Commun 12, 6035 (2021).

101. E. Pracucci, R. T. Graham, L. Alberio, G. Nardi, O. Cozzolino, V. Pillai, G. Pasquini, L. Saieva, D. Walsh, S. Landi, J. Zhang, A. J. Trevelyan, G.-M. Ratto, Daily rhythm in cortical chloride homeostasis underpins functional changes in visual cortex excitability. Nat. Commun. 14, 7108 (2023).

102. J. L. Watson, E. Seinkmane, C. T. Styles, A. Mihut, L. K. Krüger, K. E. McNally, V. J. Planelles-Herrero, M. Dudek, P. M. McCall, S. Barbiero, M. V. Oever, S. Y. Peak-Chew, B. T. Porebski, A. Zeng, N. M. Rzechorzek, D. C. S. Wong, A. D. Beale, A. Stangherlin, M. Riggi, J. Iwasa, J. Morf, C. Miliotis, A. Guna, A. J. Inglis, J. Brugués, R. M. Voorhees, J. E. Chambers, Q.-J. Meng, J. S. O’Neill, R. S. Edgar, E. Derivery, Macromolecular condensation buffers intracellular water potential. Nature, 1–11 (2023).

103. D. P. Genereux, A. Serres, J. Armstrong, J. Johnson, V. D. Marinescu, E. Murén, D. Juan, G. Bejerano, N. R. Casewell, L. G. Chemnick, J. Damas, F. D. Palma, M. Diekhans, I. T. Fiddes, M. Garber, V. N. Gladyshev, L. Goodman, W. Haerty, M. L. Houck, R. Hubley, T. Kivioja, K.-P. Koepfli, L. F. K. Kuderna, E. S. Lander, J. R. S. Meadows, W. J. Murphy, W. Nash, H. J. Noh, M. Nweeia, A. R. Pfenning, K. S. Pollard, D. A. Ray, B. Shapiro, A. F. A. Smit, M. S. Springer, C. C. Steiner, R. Swofford, J. Taipale, E. C. Teeling, J. Turner-Maier, J. Alfoldi, B. Birren, O. A. Ryder, H. A. Lewin, B. Paten, T. Marques-Bonet, K. Lindblad-Toh, E. K. Karlsson, A comparative genomics multitool for scientific discovery and conservation. Nature 587, 240–245 (2020).

104. M. J. Christmas, I. M. Kaplow, D. P. Genereux, M. X. Dong, G. M. Hughes, X. Li, P. F. Sullivan, A. G. Hindle, G. Andrews, J. C. Armstrong, M. Bianchi, A. M. Breit, M. Diekhans, C. Fanter, N. M. Foley, D. B. Goodman, L. Goodman, K. C. Keough, B. Kirilenko, A. Kowalczyk, C. Lawless, A. L. Lind, J. R. S. Meadows, L. R. Moreira, R. W. Redlich, L. Ryan, R. Swofford, A. Valenzuela, F. Wagner, O. Wallerman, A. R. Brown, J. Damas, K. Fan, J. Gatesy, J. Grimshaw, J. Johnson, S. V. Kozyrev, A. J. Lawler, V. D. Marinescu, K. M. Morrill, A. Osmanski, N. S. Paulat, B. N. Phan, S. K. Reilly, D. E. Schäffer, C. Steiner, M. A. Supple, A. P. Wilder, M. E. Wirthlin, J. R. Xue, Z. Consortium§, B. W. Birren, S. Gazal, R. M. Hubley, K.-P. Koepfli, T. Marques-Bonet, W. K. Meyer, M. Nweeia, P. C. Sabeti, B. Shapiro, A. F. A. Smit, M. S. Springer, E. C. Teeling, Z. Weng, M. Hiller, D. L. Levesque, H. A. Lewin, W. J. Murphy, A. Navarro, B. Paten, K. S. Pollard, D. A. Ray, I. Ruf, O. A. Ryder, A. R. Pfenning, K. Lindblad-Toh, E. K. Karlsson, G. Andrews, J. C. Armstrong, M. Bianchi, B. W. Birren, K. R. Bredemeyer, A. M. Breit, M. J. Christmas, H. Clawson, J. Damas, F. D. Palma, M. Diekhans, M. X. Dong, E. Eizirik, K. Fan, C. Fanter, N. M. Foley, K. Forsberg-Nilsson, C. J. Garcia, J. Gatesy, S. Gazal, D. P. Genereux, L. Goodman, J. Grimshaw, M. K. Halsey, A. J. Harris, G. Hickey, M. Hiller, A. G. Hindle, R. M. Hubley, G. M. Hughes, J. Johnson, D. Juan, I. M. Kaplow, E. K. Karlsson, K. C. Keough, B. Kirilenko, K.-P. Koepfli, J. M. Korstian, A. Kowalczyk, S. V. Kozyrev, A. J. Lawler, C. Lawless, T. Lehmann, D. L. Levesque, H. A. Lewin, X. Li, A. Lind, K. Lindblad-Toh, A. Mackay-Smith, V. D. Marinescu, T. Marques-Bonet, V. C. Mason, J. R. S. Meadows, W. K. Meyer, J. E. Moore, L. R. Moreira, D. D. Moreno-Santillan, K. M. Morrill, G. Muntané, W. J. Murphy, A. Navarro, M. Nweeia, S. Ortmann, A. Osmanski, B. Paten, N. S. Paulat, A. R. Pfenning, B. N. Phan, K. S. Pollard, H. E. Pratt, D. A. Ray, S. K. Reilly, J. R. Rosen, I. Ruf, L. Ryan, O. A. Ryder, P. C. Sabeti, D. E. Schäffer, A. Serres, B. Shapiro, A. F. A. Smit, M. Springer, C. Srinivasan, C. Steiner, J. M. Storer, K. A. M. Sullivan, P. F. Sullivan, E. Sundström, M. A. Supple, R. Swofford, J.-E. Talbot, E. Teeling, J. Turner-Maier, A. Valenzuela, F. Wagner, O. Wallerman, C. Wang, J. Wang, Z. Weng, A. P. Wilder, M. E. Wirthlin, J. R. Xue, X. Zhang, Evolutionary constraint and innovation across hundreds of placental mammals. Science 380, eabn3943 (2023).

105. A. Acharya, C. Demetriades, mTORC1 activity licenses its own release from the lysosomal surface. Mol. Cell 84, 4385–4400.e7 (2024).

106. S. Juszkiewicz, V. Chandrasekaran, Z. Lin, S. Kraatz, V. Ramakrishnan, R. S. Hegde, ZNF598 Is a Quality Control Sensor of Collided Ribosomes. Mol. Cell 72, 469–481.e7 (2018).

107. C. R. Boyd-Shiwarski, D. J. Shiwarski, S. E. Griffiths, R. T. Beacham, L. Norrell, D. E. Morrison, J. Wang, J. Mann, W. Tennant, E. N. Anderson, J. Franks, M. Calderon, K. A. Connolly, M. U. Cheema, C. J. Weaver, L. J. Nkashama, C. C. Weckerly, K. E. Querry, U. B. Pandey, C. J. Donnelly, D. Sun, A. R. Rodan, A. R. Subramanya, WNK kinases sense molecular crowding and rescue cell volume via phase separation. Cell 185, 4488–4506.e20 (2022).

108. A. R. Subramanya, C. R. Boyd-Shiwarski, Molecular Crowding: Physiologic Sensing and Control. Annu. Rev. Physiol. 86, 429–452 (2024).

109. C. R. Boyd-Shiwarski, D. J. Shiwarski, A. R. Subramanya, A New Phase for WNK Kinase Signaling Complexes as Biomolecular Condensates. Physiology 39, 269–287 (2024).

110. R. Akella, J. M. Humphreys, K. Sekulski, H. He, M. Durbacz, S. Chakravarthy, J. Liwocha, Z. J. Mohammed, C. A. Brautigam, E. J. Goldsmith, Osmosensing by WNK Kinases. Mol. Biol. Cell 32, 1614–1623 (2021).

111. S. B. Khalsa, D. Whitmore, G. D. Block, Stopping the circadian pacemaker with inhibitors of protein synthesis. Proc National Acad Sci 89, 10862–10866 (1992).

112. S. Yamaguchi, H. Isejima, T. Matsuo, R. Okura, K. Yagita, M. Kobayashi, H. Okamura, Synchronization of Cellular Clocks in the Suprachiasmatic Nucleus. Science 302, 1408–1412 (2003).

113. A. Schröder-Lorenz, L. Rensing, Circadian changes in protein-synthesis rate and protein phosphorylation in cell-free extracts of Gonyaulax polyedra. Planta 170, 7–13 (1987).

114. Y. Zhuang, Z. Li, S. Xiong, C. Sun, B. Li, S. A. Wu, J. Lyu, X. Shi, L. Yang, Y. Chen, Z. Bao, X. Li, C. Sun, Y. Chen, H. Deng, T. Li, Q. Wu, L. Qi, Y. Huang, X. Yang, Y. Lin, Circadian clocks are modulated by compartmentalized oscillating translation. Cell 186, 3245–3260.e23 (2023).

115. M. Abe, E. D. Herzog, S. Yamazaki, M. Straume, H. Tei, Y. Sakaki, M. Menaker, G. D. Block, Circadian Rhythms in Isolated Brain Regions. J Neurosci 22, 350–356 (2002).

116. U. Abraham, A. E. Granada, P. O. Westermark, M. Heine, A. Kramer, H. Herzel, Coupling governs entrainment range of circadian clocks. Mol Syst Biol 6, 438 (2010).

117. N. F. Ruby, D. E. Burns, H. C. Heller, Circadian Rhythms in the Suprachiasmatic Nucleus are Temperature-Compensated and Phase-Shifted by Heat Pulses In Vitro. J Neurosci 19, 8630– 8636 (1999).

118. V. Pilorz, P. S. Cunningham, A. Jackson, A. C. West, T. T. Wager, A. S. I. Loudon, D. A. Bechtold, A novel mechanism controlling resetting speed of the circadian clock to environmental stimuli. Curr Biology Cb 24, 766–73 (2013).

119. S. Nishide, D. Ono, Y. Yamada, S. Honma, K. Honma, De novo synthesis of PERIOD initiates circadian oscillation in cultured mouse suprachiasmatic nucleus after prolonged inhibition of protein synthesis by cycloheximide. Eur J Neurosci 35, 291–299 (2012).

120. A. C. Hsieh, Y. Liu, M. P. Edlind, N. T. Ingolia, M. R. Janes, A. Sher, E. Y. Shi, C. R. Stumpf, C. Christensen, M. J. Bonham, S. Wang, P. Ren, M. Martin, K. Jessen, M. E. Feldman, J. S. Weissman, K. M. Shokat, C. Rommel, D. Ruggero, The translational landscape of mTOR signalling steers cancer initiation and metastasis. Nature 485, 55–61 (2012).

121. M. A. McNurlan, A. M. Tomkins, P. J. Garlick, The effect of starvation on the rate of protein synthesis in rat liver and small intestine. Biochem J 178, 373–9 (1979).

122. F. Yoshizawa, T. Nagasawa, N. Nishizawa, R. Funabiki, Protein Synthesis and Degradation Change Rapidly in Response to Food Intake in Muscle of Food-Deprived Mice,. J Nutrition 127, 1156–1159 (1997).

123. T. Naito, A. Kuma, N. Mizushima, Differential Contribution of Insulin and Amino Acids to the mTORC1-Autophagy Pathway in the Liver and Muscle*. J. Biol. Chem. 288, 21074–21081 (2013).

124. S. Daan, K. Spoelstra, U. Albrecht, I. Schmutz, M. Daan, B. Daan, F. Rienks, I. Poletaeva, G. Dell’Omo, A. Vyssotski, H.-P. Lipp, Lab Mice in the Field: Unorthodox Daily Activity and Effects of a Dysfunctional Circadian Clock Allele. J Biol Rhythm 26, 118–129 (2011).

125. R. A. Hut, V. Pilorz, A. S. Boerema, A. M. Strijkstra, S. Daan, Working for Food Shifts Nocturnal Mouse Activity into the Day. Plos One 6, e17527 (2011).

126. G. Perrigo, Breeding and feeding strategies in deer mice and house mice when females are challenged to work for their food. Anim Behav 35, 1298–1316 (1987).

127. A. L. van Rosmalen, S. J. Riede, V. Pilorz, T. Adage, A. J. W. Scheurink, V. van der Vinne, R. Hut, “Nocturnal and Diurnal Behavior Assessed by the ‘Work-for-Food’ Protocol in Small Rodents” in Circadian Clocks (2022)Neuromethods, pp. 187–216.

128. L. van Rosmalen, J. Zhu, G. Maier, E. G. Gacasan, T. Lin, E. Zhemchuzhnikova, V. Rothenberg, S. Razu, S. Deota, R. K. Ramasamy, R. L. Sah, A. D. McCulloch, R. A. Hut, S. Panda, Multi-organ transcriptome atlas of a mouse model of relative energy deficiency in sport. Cell Metab. 36, 2015–2037.e6 (2024).

129. V. van der Vinne, S. J. Riede, J. A. Gorter, W. G. Eijer, M. T. Sellix, M. Menaker, S. Daan, V. Pilorz, R. A. Hut, Cold and hunger induce diurnality in a nocturnal mammal. Proc National Acad Sci 111, 15256–15260 (2014).

130. L. van Rosmalen, S. Deota, G. Maier, H. D. Le, T. Lin, R. K. Ramasamy, R. A. Hut, S. Panda, Energy balance drives diurnal and nocturnal brain transcriptome rhythms. Cell Rep. 43, 113951–113951 (2024).

131. H. H. Pak, A. N. Grossberg, R. R. Sanderfoot, R. Babygirija, C. L. Green, M. Koller, M. Dzieciatkowska, D. A. Paredes, D. W. Lamming, Non-canonical metabolic and molecular effects of calorie restriction are revealed by varying temporal conditions. Cell Rep. 43, 114663 (2024).

132. V. A. Acosta-Rodríguez, M. H. M. de Groot, F. Rijo-Ferreira, C. B. Green, J. S. Takahashi, Mice under Caloric Restriction Self-Impose a Temporal Restriction of Food Intake as Revealed by an Automated Feeder System. Cell Metab. 26, 267–277.e2 (2017).

133. R. E. Lawrence, K. F. Cho, R. Rappold, A. Thrun, M. Tofaute, D. J. Kim, O. Moldavski, J. H. Hurley, R. Zoncu, A nutrient-induced affinity switch controls mTORC1 activation by its Rag GTPase–Ragulator lysosomal scaffold. Nat. Cell Biol. 20, 1052–1063 (2018).

134. X. Gu, J. M. Orozco, R. A. Saxton, K. J. Condon, G. Y. Liu, P. A. Krawczyk, S. M. Scaria, J. W. Harper, S. P. Gygi, D. M. Sabatini, SAMTOR is an S-adenosylmethionine sensor for the mTORC1 pathway. Science 358, 813–818 (2017).

135. K. Devarajan, M. Fidino, Z. J. Farris, S. A. Adalsteinsson, G. Andrade-Ponce, J. L. Angstmann, W. Anthonysamy, J. Aquino, A. Asefa, B. Avila, L. L. Bailey, L. M. de S. Barbosa, M. de F. Barreto, O. Barton, C. E. Bates, M. G. Beltrão, T. Bird, E. G. Biro, F. Bisi, D. Bohórquez, M. Boyce, J. S. Brashares, G. Bullington, P. Burns, J. Burr, A. R. Butler, K. L. Calhoun, T. T. Cao, N. Casado, J. C. Cepeda-Duque, J. D. Cepek, A. G. Chiarello, M. Collins, P. Cordeiro-Estrela, S. Costa, G. Cremonesi, B. Cristescu, P. Cruz, A. C. F. de Albuquerque, C. D. Angelo, C. B. de Campos, L. M. M. de Sena, M. D. Bitetti, D. de M. Dias, D. Diefenbach, T. S. Doherty, T. P. dos Santos, G. T. Duarte, T. M. Eppley, J. Erb, C. F. Esteves, B. Evans, M. L. M. Falcão, H. Fernandes-Ferreira, J. R. Fieberg, L. C. F. de S. Filho, J. Fisher, M.-J. Fortin, G. A. Gale, T. Gallo, L. S. Ganoe, R. Garcia-Anleu, K. M. Gaynor, T. A. Gelmi-Candusso, P. N. Gichuru, Q. Gomez, A. M. Green, L. N. Guimarães, J. D. Haight, L. R. Harris, Z. D. Hawn, J. Heiman, H. Q. Hoang, S. Huebner, F. Iannarilli, M. E. Iezzi, J. S. Ivan, K. J. Jaspers, M. J. Jordan, J. Kamilar, M. Kane, M. H. Karimi, M. Kelly, M. T. Kohl, W. P. KuvleskyJr., A. Ladle, R. N. Larson, Q. T. Le, D. Le, V. S. Le, E. W. Lehrer, P. E. Lendrum, J. Lewis, A. Link, D. J. Lizcano, J. V. Lombardi, R. Long, E. López-Tello, C. Lugarini, D. Lugo, P. MacKay, M. Madadi, R. A. Magalhães, S. B. Magle, L. H. R. D. Maia, S. Mandujano, T. Marchenkova, P. H. Marinho, L. Marker, J. M. Pardo, A. Martinoli, R. L. Massara, J. Masseloux, D. Matiukhina, A. Mayer, L. Mazariegos, M. R. McClung, A. McInturff, D. McPhail, A. Mertl, C. R. Middaugh, D. Miller, D. Mills, D. Miquelle, V. Miritis, R. J. Moll, P. Molnár, R. A. Montgomery, T. L. Morelli, A. Mortelliti, R. I. Mueller, A. S. Mukhacheva, K. Mullen, A. Murphy, V. Nepomuceno, D. Ngoprasert, A. Nguyen, T. V. Nguyen, V. T. Nguyen, H. A. N. Quang, R. Nipko, A. C. C. Nobre, J. Northrup, M. A. Owen, A. P. Paglia, M. S. Palmer, G. Palomo-Munoz, L. E. Pardo, C. Parks, A. M. de O. Paschoal, B. Patterson, A. Paviolo, L. Pejchar, M. E. Pendergast, H. L. Perotto-Baldivieso, T. Petrov, M. K. P. Poisson, D. J. Polli, M. Pourmirzai, A. Reebin, K. R. Remine, L. Rich, C. S. Richardson, F. Robino, D. G. Rocha, F. L. Rocha, F. H. G. Rodrigues, A. T. Rohnke, T. J. Ryan, C. M. Salsbury, H. A. Sander, N. M. da C. Santos-Cavalcante, C. H. Sekercioglu, I. Seryodkin, D. H. Setiawan, S. Shadloo, M. Shahhosseini, G. Shannon, C. J. Shier, G. B. Smith, T. Snyder, R. Sollmann, K. L. Sparks, K. Sribuarod, C. C. St. Clair, T. Stankowich, R. Steinmetz, C. J. Stevenson, S. Sunarto, T. D. Surasinghe, S. V. Sutyrina, R. R. Swaisgood, A. Taktehrani, K. Thapa, M. Thorton, A. Tilker, M. W. Tobler, V. B. Tran, J. Tucker, R. C. V. Horn, J. S. Vargas-Soto, K. L. Velásquez-C, J. Venter, E. M. Venticinque, S. Verschueren, E. Wampole, D. J. Watchorn, O. R. Wearn, K. C. B. Weiss, A. Welschen, F. A. Widodo, J. Williamson, A. Wilting, G. Wittemyer, A. Zavaleta, A. J. Zellmer, B. D. Gerber, When the wild things are: Defining mammalian diel activity and plasticity. Sci. Adv. 11, eado3843 (2025).

136. S. Battaglioni, D. Benjamin, M. Wälchli, T. Maier, M. N. Hall, mTOR substrate phosphorylation in growth control. Cell 185, 1814–1836 (2022).

137. M. Dudek, D. R. J. Pathiranage, B. Bano-Otalora, A. Paszek, N. Rogers, C. F. Gonçalves, C. Lawless, D. Wang, Z. Luo, L. Yang, F. Guilak, J. A. Hoyland, Q.-J. Meng, Mechanical loading and hyperosmolarity as a daily resetting cue for skeletal circadian clocks. Nat. Commun. 14, 7237 (2023).

138. H. Yoshitane, K. Imamura, T. Okubo, Y. Otobe, S. Kawakami, S. Ito, T. Takumi, K. Hattori, I. Naguro, H. Ichijo, Y. Fukada, mTOR-AKT Signaling in Cellular Clock Resetting Triggered by Osmotic Stress. Antioxid. Redox Signal. 37, 631–646 (2022).

139. B. Saer, G. Taylor, A. Hayes, B. Ananthasubramaniam, J.-M. Fustin, Dietary methionine/choline deficiency affects behavioural and molecular circadian rhythms. bioRxiv, 2024.01.04.574014 (2024).

140. S. E. Doyle, T. Yoshikawa, H. Hillson, M. Menaker, Retinal pathways influence temporal niche. Proc. National Acad. Sci. 105, 13133–13138 (2008).

141. R. A. Schoonderwoerd, P. T. Gutiérrez, R. Blommers, A. W. Beurden, T. C. J. J. Coenen, N. J. Klett, S. H. Michel, J. H. Meijer, Inhibitory responses to retinohypothalamic tract stimulation in the circadian clock of the diurnal rodent Rhabdomys pumilio. FASEB J. 36, e22415 (2022).

142. R. Caputo, R. A. Schoonderwoerd, A. Ramkisoensing, J. A. M. Janse, H. C. van Diepen, S. Raison, P. Pévet, N. A. V. Derks, D. Sage-Ciocca, T. Deboer, E. Challet, J. H. Meijer, Physical activity increases neuronal activity in the circadian clock of diurnal Arvicanthis ansorgei. bioRxiv, 2022.05.31.493966 (2024).

143. R. Richardson, C. Y. Feigin, B. Bano-Otalora, M. R. Johnson, A. E. Allen, J. Park, R. J. McDowell, S. A. Mereby, I.-H. Lin, R. J. Lucas, R. Mallarino, The genomic basis of temporal niche evolution in a diurnal rodent. Curr. Biol. 33, 3289–3298.e6 (2023).

144. V. van der Vinne, J. A. Gorter, S. J. Riede, R. A. Hut, Diurnality as an energy-saving strategy: energetic consequences of temporal niche switching in small mammals. J. Exp. Biology 218, 2585–2593 (2015).

145. S. J. Riede, V. van der Vinne, R. A. Hut, The flexible clock: predictive and reactive homeostasis, energy balance and the circadian regulation of sleep–wake timing. J Exp Biol 220, 738–749 (2017).

146. V. van der Vinne, P. Tachinardi, S. J. Riede, J. Akkerman, J. Scheepe, S. Daan, R. A. Hut, Maximising survival by shifting the daily timing of activity. Ecol Lett 22, 2097–2102 (2019).

147. N. M. Weyer, A. Fuller, A. J. Haw, L. C. R. Meyer, D. Mitchell, M. Picker, B. Rey, R. S. Hetem, Increased Diurnal Activity Is Indicative of Energy Deficit in a Nocturnal Mammal, the Aardvark. Front. Physiol. 11, 637 (2020).

148. A. Stangherlin, E. Seinkmane, J. S. O’Neill, Understanding circadian regulation of mammalian cell function, protein homeostasis, and metabolism. Curr Opin Syst Biology 28, 100391 (2021).

149. D. C. S. Wong, E. Seinkmane, A. Zeng, A. Stangherlin, N. M. Rzechorzek, A. D. Beale, J. Day, M. Reed, S. Y. Peak-Chew, C. T. Styles, R. S. Edgar, M. Putker, J. S. O’Neill, CRYPTOCHROMES promote daily protein homeostasis. Embo J 41, e2021108883 (2022).

150. E. Seinkmane, A. Edmondson, S. Y. Peak-Chew, A. Zeng, N. M. Rzechorzek, N. R. James, J. West, J. Munns, D. C. Wong, A. D. Beale, J. S. O’Neill, Circadian regulation of macromolecular complex turnover and proteome renewal. EMBO J. 43, 2813–2833 (2024).

151. H. C. Causton, K. A. Feeney, C. A. Ziegler, J. S. O’Neill, Metabolic Cycles in Yeast Share Features Conserved among Circadian Rhythms. Curr Biol 25, 1056–1062 (2015).

152. J. S. O’Neill, N. P. Hoyle, J. B. Robertson, R. S. Edgar, A. D. Beale, S. Y. Peak-Chew, J. Day, A. S. H. Costa, C. Frezza, H. C. Causton, Eukaryotic cell biology is temporally coordinated to support the energetic demands of protein homeostasis. Nat Commun 11, 4706 (2020).

153. M. O. Moreira, Y. Qu, J. J. Wiens, Large-scale evolution of body temperatures in land vertebrates. Evol. Lett. 5, 484–494 (2021).

154. A. Lewden, C. M. Bishop, G. N. Askew, How birds dissipate heat before, during and after flight. J. R. Soc. Interface 20, 20230442 (2023).

155. I. Kelava, I. Chiaradia, L. Pellegrini, A. T. Kalinka, M. A. Lancaster, Androgens increase excitatory neurogenic potential in human brain organoids. Nature 602, 112–116 (2022).

156. F. W. Lindhout, F. M. Krienen, K. S. Pollard, M. A. Lancaster, A molecular and cellular perspective on human brain evolution and tempo. Nature 630, 596–608 (2024).

157. D. P. Iyer, H. H. Khoei, V. A. van der Weijden, H. Kagawa, S. J. Pradhan, M. Novatchkova, A. McCarthy, T. Rayon, C. S. Simon, I. Dunkel, S. E. Wamaitha, K. Elder, P. Snell, L. Christie, E. G. Schulz, K. K. Niakan, N. Rivron, A. Bulut-Karslioğlu, mTOR activity paces human blastocyst stage developmental progression. Cell 187, 6566–6583.e22 (2024).

158. R. Iwata, P. Vanderhaeghen, Metabolic mechanisms of species-specific developmental tempo. Dev. Cell 59, 1628–1639 (2024).

159. S. A. Brown, F. Fleury-Olela, E. Nagoshi, C. Hauser, C. Juge, C. A. Meier, R. Chicheportiche, J.-M. Dayer, U. Albrecht, U. Schibler, The Period Length of Fibroblast Circadian Gene Expression Varies Widely among Human Individuals. PLoS Biol. 3, e338 (2005).

160. J. M. Lotthammer, J. Hernández-García, D. Griffith, D. Weijers, A. S. Holehouse, R. J. Emenecker, Metapredict enables accurate disorder prediction across the Tree of Life. bioRxiv, 2024.11.05.622168 (2024).

161. A. Seluanov, A. Vaidya, V. Gorbunova, Establishing Primary Adult Fibroblast Cultures From Rodents. J Vis Exp, doi: 10.3791/2033 (2010).

162. J. Drouin-Ouellet, S. Lau, P. L. Brattås, D. R. Ottosson, K. Pircs, D. A. Grassi, L. M. Collins, R. Vuono, A. A. Sjöland, G. Westergren-Thorsson, C. Graff, L. Minthon, H. Toresson, R. A. Barker, J. Jakobsson, M. Parmar, REST suppression mediates neural conversion of adult human fibroblasts via microRNA-dependent and -independent pathways. EMBO Mol. Medicine 9, 1117–1131 (2017).

163. U. K. Valekunja, R. S. Edgar, M. Oklejewicz, G. T. J. van der Horst, J. S. O’Neill, F. Tamanini, D. J. Turner, A. B. Reddy, Histone methyltransferase MLL3 contributes to genome-scale circadian transcription. Proc National Acad Sci 110, 1554–1559 (2013).

164. K. Labun, T. G. Montague, J. A. Gagnon, S. B. Thyme, E. Valen, CHOPCHOP v2: a web tool for the next generation of CRISPR genome engineering. Nucleic Acids Res 44, W272–W276 (2016).

165. A. Zeng, J. S. O’Neill, “Using ALLIGATORs to Capture Circadian Bioluminescence” in Circadian Regulation (2022)vol. 2482 of Methods in Molecular Biology, pp. 125–135.

166. P. Crosby, N. P. Hoyle, J. S. O’Neill, Flexible Measurement of Bioluminescent Reporters Using an Automated Longitudinal Luciferase Imaging Gas- and Temperature-optimized Recorder (ALLIGATOR). J Vis Exp, doi: 10.3791/56623 (2017).

167. P. Stothard, The Sequence Manipulation Suite: JavaScript Programs for Analyzing and Formatting Protein and DNA Sequences. BioTechniques 28, 1102–1104 (2000).

168. Y. Nakamura, T. Gojobori, T. Ikemura, Codon usage tabulated from international DNA sequence databases: status for the year 2000. Nucleic Acids Res. 28, 292–292 (2000).

169. A. R. Gruber, R. Lorenz, S. H. Bernhart, R. Neuböck, I. L. Hofacker, The Vienna RNA websuite. Nucleic Acids Res 36, W70–4 (2008).

170. H. L. Sladitschek, P. A. Neveu, MXS-Chaining: A Highly Efficient Cloning Platform for Imaging and Flow Cytometry Approaches in Mammalian Systems. Plos One 10, e0124958 (2015).

171. R. Cencic, J. Pelletier, Hippuristanol - A potent steroid inhibitor of eukaryotic initiation factor 4A. Translation 4, e1137381 (2016).

172. S. Yamazaki, R. Numano, M. Abe, A. Hida, R. Takahashi, M. Ueda, G. D. Block, Y. Sakaki, M. Menaker, H. Tei, Resetting Central and Peripheral Circadian Oscillators in Transgenic Rats. Science 288, 682–685 (2000).

173. T. Yoshikawa, S. Yamazaki, M. Menaker, Effects of Preparation Time on Phase of Cultured Tissues Reveal Complexity of Circadian Organization. J Biol Rhythm 20, 500–512 (2005).

174. T. Noguchi, M. Ikeda, Y. Ohmiya, Y. Nakajima, A Dual-Color Luciferase Assay System Reveals Circadian Resetting of Cultured Fibroblasts by Co-Cultured Adrenal Glands. Plos One 7, e37093 (2012).

175. M. H. Hastings, A. B. Reddy, D. G. McMahon, E. S. Maywood, Analysis of Circadian Mechanisms in the Suprachiasmatic Nucleus by Transgenesis and Biolistic Transfection. Methods Enzymol 393, 579–592 (2005).

176. O. Wagih, N. Sugiyama, Y. Ishihama, P. Beltrao, Uncovering Phosphorylation-Based Specificities through Functional Interaction Networks*. Mol Cell Proteomics 15, 236–245 (2016).

177. J. Ou, H. Liu, N. K. Nirala, A. Stukalov, U. Acharya, M. R. Green, L. J. Zhu, dagLogo: An R/Bioconductor package for identifying and visualizing differential amino acid group usage in proteomics data. Plos One 15, e0242030 (2020).

178. A. Kowalczyk, W. K. Meyer, R. Partha, W. Mao, N. L. Clark, M. Chikina, RERconverge: an R package for associating evolutionary rates with convergent traits. Bioinformatics 35, 4815– 4817 (2019).

179. J. Reimand, M. Kull, H. Peterson, J. Hansen, J. Vilo, g:Profiler—a web-based toolset for functional profiling of gene lists from large-scale experiments. Nucleic Acids Res. 35, W193– W200 (2007).

180. C. Bárcena, P. M. Quirós, S. Durand, P. Mayoral, F. Rodríguez, X. M. Caravia, G. Mariño, C. Garabaya, M. T. Fernández-García, G. Kroemer, J. M. P. Freije, C. López-Otín, Methionine Restriction Extends Lifespan in Progeroid Mice and Alters Lipid and Bile Acid Metabolism. Cell Rep. 24, 2392–2403 (2018).

181. M. Comas, R. A. Hut, Twilight and Photoperiod Affect Behavioral Entrainment in the House Mouse (Mus musculus). J. Biol. Rhythm. 24, 403–412 (2009).

182. J. Ponten, E. Saksela, Two established in vitro cell lines from human mesenchymal tumours. Int J Cancer 2, 434–447 (1967).

183. K. A. Lamia, U. M. Sachdeva, L. DiTacchio, E. C. Williams, J. G. Alvarez, D. F. Egan, D. S. Vasquez, H. Juguilon, S. Panda, R. J. Shaw, C. B. Thompson, R. M. Evans, AMPK Regulates the Circadian Clock by Cryptochrome Phosphorylation and Degradation. Science 326, 437–440 (2009).

184. C. H. Gabriel, M. del Olmo, A. Zehtabian, M. Jäger, S. Reischl, H. van Dijk, C. Ulbricht, A. Rakhymzhan, T. Korte, B. Koller, A. Grudziecki, B. Maier, A. Herrmann, R. Niesner, T. Zemojtel, H. Ewers, A. E. Granada, H. Herzel, A. Kramer, Live-cell imaging of circadian clock protein dynamics in CRISPR-generated knock-in cells. Nat Commun 12, 3796 (2021).

185. M. Zhou, T. Wang, J. Fu, G. Xiao, Y. Liu, Nonoptimal codon usage influences protein structure in intrinsically disordered regions. Mol Microbiol 97, 974–87 (2015).

186. J. F. Pelham, J. C. Dunlap, J. M. Hurley, Intrinsic disorder is an essential characteristic of components in the conserved circadian circuit. Cell Commun Signal 18, 181 (2020).

187. Y. Liu, A code within the genetic code: codon usage regulates co-translational protein folding. Cell Commun Signal 18, 145 (2020).

188. K. M. Sakamoto, K. B. Kim, A. Kumagai, F. Mercurio, C. M. Crews, R. J. Deshaies, Protacs: Chimeric molecules that target proteins to the Skp1–Cullin–F box complex for ubiquitination and degradation. Proc. National Acad. Sci. 98, 8554–8559 (2001).

189. M. Békés, D. R. Langley, C. M. Crews, PROTAC targeted protein degraders: the past is prologue. Nat. Rev. Drug Discov. 21, 181–200 (2022).

190. C. S. Pittendrigh, Circadian Rhythms and the Circadian Organization of Living Systems. Cold Spring Harb Sym 25, 159--184 (1960).

191. J. Aschoff, Exogenous and Endogenous Components in Circadian Rhythms. Cold Spring Harb Sym 25, 11–28 (1960).

192. R. Partha, A. Kowalczyk, N. L. Clark, M. Chikina, Robust Method for Detecting Convergent Shifts in Evolutionary Rates. Mol. Biol. Evol. 36, 1817–1830 (2019).

193. K. Ninomiya, N. Kataoka, M. Hagiwara, Stress-responsive maturation of Clk1/4 pre-mRNAs promotes phosphorylation of SR splicing factor. J Cell Biol 195, 27–40 (2011).

194. K. Düvel, J. L. Yecies, S. Menon, P. Raman, A. I. Lipovsky, A. L. Souza, E. Triantafellow, Q. Ma, R. Gorski, S. Cleaver, M. G. V. Heiden, J. P. MacKeigan, P. M. Finan, C. B. Clish, L. O. Murphy, B. D. Manning, Activation of a Metabolic Gene Regulatory Network Downstream of mTOR Complex 1. Mol. Cell 39, 171–183 (2010).

195. I. Seyrling, P. W. Dierkes, A. L. Burger, Diurnal and Nocturnal Behaviour of Cheetahs (Acinonyx jubatus) and Lions (Panthera leo) in Zoos. Anim.: Open Access J. MDPI 12, 2367 (2022).

196. K. Rafiq, N. R. Jordan, K. Golabek, J. W. McNutt, A. Wilson, B. Abrahms, Increasing ambient temperatures trigger shifts in activity patterns and temporal partitioning in a large carnivore guild. Proc. R. Soc. B 290, 20231938 (2023).

197. K. E. Jones, J. Bielby, M. Cardillo, S. A. Fritz, J. O’Dell, C. D. L. Orme, K. Safi, W. Sechrest, E. H. Boakes, C. Carbone, C. Connolly, M. J. Cutts, J. K. Foster, R. Grenyer, M. Habib, C. A. Plaster, S. A. Price, E. A. Rigby, J. Rist, A. Teacher, O. R. P. Bininda-Emonds, J. L. Gittleman, G. M. Mace, A. Purvis, PanTHERIA: a species-level database of life history, ecology, and geography of extant and recently extinct mammals. Ecology 90, 2648–2648 (2009).

198. D. E. Wilson, R. A. Mittermeier, Eds., Handbook of the Mammals of the World, Vol. 1: Carnivores (Lynx Edicions, 2009)vol. 1.

199. K. A. Bugler, J. G. Ross, A. M. Paterson, Activity Patterns of Captive Red Panda (Ailurus fulgens). Animals 13, 846 (2023).

200. R. M. Nowak, Ed., Walker’s Mammals of the World (ed. 6, 1999).

201. D. J. Chivers, ON THE DAILY BEHAVIOUR AND SPACING OF HOWLING MONKEY GROUPS. Folia Primatol. 10, 48–102 (1969).

202. A. M. Prpi, P. Ganevi, T. Safner, K. Kavi, K. Jerina, N. prem, Activity patterns of aoudad (Ammotragus lervia) in a Mediterranean habitat. J. Vertebr. Biol. 69, 1–8 (2020).

203. J. J. Bennie, J. P. Duffy, R. Inger, K. J. Gaston, Biogeography of time partitioning in mammals. Proc. Natl. Acad. Sci. 111, 13727–13732 (2014).

204. P. F. Jones, T. A. Hurly, C. Jensen, K. Zimmer, A. Jakes, Diel and Monthly Movement Rates by Migratory and Resident Female Pronghorn. The Prairie Naturalist 49, 3–12 (2017).

205. T. Ikeda, J. A. Autio, A. Kawasaki, C. Takeda, T. Ose, M. Takada, D. C. V. Essen, M. F. Glasser, T. Hayashi, Cortical adaptation of the night monkey to a nocturnal niche environment: a comparative non-invasive T1w/T2w myelin study. Brain Struct. Funct. 228, 1107–1123 (2023).

206. S. R. Anderson, J. J. Wiens, Out of the dark: 350 million years of conservatism and evolution in diel activity patterns in vertebrates. Evolution 71, 1944–1959 (2017).

207. J. Muñoz-Delgado, B. Fuentes-Pardo, A. E. Baum, N. Lanzagorta, R. Arenas-Rosas, A. M. Santillán-Doherty, M. A. Guevara, M. Corsi-Cabrera, Presence of a circadian rhythm in the spider monkey’s (Ateles geoffroyi) motor activity. Biol. Rhythm Res. 36, 115–122 (2005).

208. C. D. Soria, M. Pacifici, M. D. Marco, S. M. Stephen, C. Rondinini, COMBINE: a coalesced mammal database of intrinsic and extrinsic traits. Ecology 102, e03344 (2021).

209. D. Wang, Q. Li, L. Hou, X. Su, X. Lian, Spatiotemporal overlap among snow leopard, bharal, and free-ranging livestock: Suggestions on mitigating human-snow leopard conflict. Glob. Ecol. Conserv. 53, e03029 (2024).

210. C. Giannetto, G. Piccione, Daily rhythms of 25 physiological variables in Bos taurus maintained under natural conditions. J. Appl. Biomed. 7, 55–61 (2009).

211. D. E. Wilson, R. A. Mittermeier, Eds., Handbook of the Mammals of the World, Vol. 2: Hoofed Mammals (2011)vol. 2.

212. H. G. Erkert, B. Nagel, I. Stephani, Light and social effects on the free-running circadian activity rhythm in common marmosets (Callithrix jacchus; Primates): social masking, pseudo-splitting, and relative coordination. Behav. Ecol. Sociobiol. 18, 443–452 (1986).

213. Y. Xue, D. Li, W. Xiao, F. Liu, Y. Zhang, X. Wang, H. Jia, Activity patterns of wild Bactrian camels (Camelus bactrianus) in the northern piedmont of the Altun Mountains, China. Anim. Biol. 65, 209–217 (2015).

214. H. Farsi, D. Harti, M. R. Achaâban, M. Piro, M. Ouassat, E. Challet, P. Pévet, K. E. Allali, Seasonal variations in locomotor activity rhythm and diurnal activity in the dromedary camel (Camelus dromedarius) under mesic semi-natural conditions. Chronobiol. Int. 39, 129–150 (2022).

215. B. Nasanbat, F. Ceacero, S. Ravchig, A small neighborhood well-organized: seasonal and daily activity patterns of the community of large and mid-sized mammals around waterholes in the Gobi Desert, Mongolia. Front. Zoöl. 18, 25 (2021).

216. Y. Xue, J. Li, G. Sagen, Y. Zhang, Y. Dai, D. Li, Activity patterns and resource partitioning: seven species at watering sites in the Altun Mountains, China. J. Arid Land 10, 959–967 (2018).

217. M. C. Nicholson, T. P. Husband, Diurnal Behavior of the Agrimi, Capra aegagrus. J. Mammal. 73, 135–142 (1992).

218. H. Farsi, D. Harti, M. R. Achaâban, M. Piro, V. Raverot, B. Bothorel, M. Ouassat, E. Challet, P. Pévet, K. E. Allali, Melatonin rhythm and other outputs of the master circadian clock in the desert goat (Capra hircus) are entrained by daily cycles of ambient temperature. J. Pineal Res. 68, e12634 (2020).

219. A. B. Taber, C. P. Doncaster, N. N. Neris, F. H. Colman, Ranging Behavior and Population Dynamics of the Chacoan Peccary, Catagonus wagneri. J. Mammal. 74, 443–454 (1993).

220. C. Künzl, N. Sachser, The Behavioral Endocrinology of Domestication: A Comparison between the Domestic Guinea Pig (Cavia apereaf.porcellus) and Its Wild Ancestor, the Cavy (Cavia aperea). Horm. Behav. 35, 28–37 (1999).

221. M. Akita, K. Ishii, M. Kuwahara, H. Tsubone, The Daily Pattern of Heart Rate, Body Temperature, and Locomotor Activity in Guinea Pigs. Exp. Anim. 50, 409 (2003).

222. M. Quispe-López, S. Barreda, D. Marcelo-Carranza, R. Mejía, C. Santana, D. W. Ramirez, Patrones de actividad diaria y lunar de Cavia tschudii (Rodentia) en un humedal costero tropical. Mastozoología Neotropical 29, 001–010 (2022).

223. T. D. Lambert, R. W. Kays, P. A. Jansen, E. Aliaga-Rossel, M. Wikelski, Nocturnal activity by the primarily diurnal Central American agouti (Dasyprocta punctata) in relation to environmental conditions, resource abundance and predation risk. J. Trop. Ecol. 25, 211–215 (2008).

224. A. Gatica, A. C. Ochoa, A. M. Mangione, Potential predators of Dolichotis patagonum in the surroundings of its burrows, in Sierra de las Quijadas National Park, San Luis, Argentina. Mammalia 86, 13–21 (2022).

225. G. G. Esslinger, J. L. Bodkin, A. R. Breton, J. M. Burns, D. H. Monson, Temporal patterns in the foraging behavior of sea otters in Alaska. J. Wildl. Manag. 78, 689–700 (2014).

226. C. Bertolucci, C. Giannetto, F. Fazio, G. Piccione, Seasonal variations in daily rhythms of activity in athletic horses. Animal 2, 1055–1060 (2008).

227. A.-M. Martin, J. A. Elliott, P. Duffy, C. M. Blake, S. B. Attia, L. M. Katz, J. A. Browne, V. Gath, B. A. McGivney, E. W. Hill, B. A. Murphy, Circadian regulation of locomotor activity and skeletal muscle gene expression in the horse. J. Appl. Physiol. 109, 1328–1336 (2010).

228. A. Berger, K.-M. Scheibe, K. Eichhorn, A. Scheibe, J. Streich, Diurnal and ultradian rhythms of behaviour in a mare group of Przewalski horse (Equus ferus przewalskii), measured through one year under semi-reserve conditions. Appl. Anim. Behav. Sci. 64, 1–17 (1999).

229. J. Bray, D. R. Samson, C. L. Nunn, Activity patterns in seven captive lemur species: Evidence of cathemerality in Varecia and Lemur catta? Am J Primatol 79, e22648 (2017).

230. G. Donati, A. Lunardini, P. M. Kappeler, New Directions in Lemur Studies. 119–137 (1999).

231. S. Streicher, J. G. Boyles, M. K. Oosthuizen, N. C. Bennett, Body Temperature Patterns and Rhythmicity in Free-Ranging Subterranean Damaraland Mole-Rats, Fukomys damarensis. PLoS ONE 6, e26346 (2011).

232. M. Clauss, M. Scriba, J. Kioko, J. U. Ganzhorn, C. Kiffner, Camera-trap data do not indicate scaling of diel activity and cathemerality with body mass in an East African mammal assemblage. Ecol. Evol. 11, 13846–13861 (2021).

233. C. M. Francis, A Guide to the Mammals of Southeast Asia (Princeton University Press, 2008).

234. C. J. R. Alho, N. L. Rondon, Habitats, population densities, and social structure of capybaras (Hydrochaeris Hydrochaeris, Rodentia) in the Pantanal, Brazil. Rev. Bras. Zoöl. 4, 139–149 (1987).

235. G. S. A. Rasmussen, D. W. Macdonald, Masking of the zeitgeber: African wild dogs mitigate persecution by balancing time. J. Zoöl. 286, 232–242 (2012).

236. Z. M, F. L, R. AM, A. T, A. M, Daily Activity and Foraging Patterns of Adult Golden Marmots in Pup-Rearing Burrows in Relation to Habitat Disturbance in Karakorum Range Pakistan. J Ethol & Animal Sci 4, 000124 (2022).

237. U. Roll, T. Dayan, N. Kronfeld-Schor, On the role of phylogeny in determining activity patterns of rodents. Evol. Ecol. 20, 479–490 (2006).

238. F. Cayetanot, E. J. W. V. Someren, M. Perret, F. Aujard, Shortened Seasonal Photoperiodic Cycles Accelerate Aging of the Diurnal and Circadian Locomotor Activity Rhythms in a Primate. J. Biol. Rhythm. 20, 461–469 (2005).

239. S. B. C. Bisceglia, J. A. Pereira, P. Teta, R. D. Quintana, Rodent selection by Geoffroy’s cats in a semi-arid scrubland of central Argentina. J. Arid Environ. 75, 1024–1028 (2011).

240. C. A. Nichols, K. Alexander, Creeping in the night: What might ecologists be missing? PLoS ONE 13, e0198277 (2018).

241. A. T. Smith, Y. Xie, R. S. Hoffmann, D. Lunde, J. MacKinnon, D. E. Wilson, W. C. Wozencraft, Eds., A Guide to the Mammals of China (Princeton University Press, 2010).

242. J. F. Redman, “Effect of light on the circadian activity rhythm of the slow loris, Nycticebus coucang,” thesis, University of the Pacific (1979).

243. R. Zhou, R. Hua, Z. Tang, L. Hua, Daily and Seasonal Activity Patterns of Plateau Pikas (Ochotona curzoniae) on the Qinghai–Tibet Plateau, China, and Their Relationship with Weather Condition. Animals 13, 1689 (2023).

244. M. J. H. Kas, D. M. Edgar, A Nonphotic Stimulus Inverts the Diurnal–Nocturnal Phase Preference in Octodon degus. J. Neurosci. 19, 328–333 (1999).

245. J. Hart, T. Hart, Tracking the rainforest giraffe. Animal Kingdom 91, 26–32 (1988).

246. V. R. Squires, Ecology and behaviour of domestic sheep (Ovis aries): a review. Mammal Rev. 5, 35–57 (1975).

247. C. A. Stockwell, G. C. Bateman, J. Berger, Conflicts in national parks: A case study of helicopters and bighorn sheep time budgets at the grand canyon. Biol. Conserv. 56, 317–328 (1991).

248. J. A. Alderman, P. R. Krausman, B. D. Leopold, Diel Activity of Female Desert Bighorn Sheep in Western Arizona. J. Wildl. Manag. 53, 264 (1989).

249. H. Yang, S. Han, B. Xie, P. Mou, X. Kou, T. Wang, J. Ge, L. Feng, Do prey availability, human disturbance and habitat structure drive the daily activity patterns of Amur tigers (Panthera tigris altaica)? J. Zoöl. 307, 131–140 (2019).

250. Y. Luo, L. Wang, L. Yang, M. Tan, Y. Wu, Y. Li, Z. Li, Puppet resting behavior in the Tibetan antelope (Pantholops hodgsonii). PLoS ONE 13, e0204379 (2018).

251. P. Withers, Ecology of a small mammal community on a rocky outcrop in the Namib Desert. Madoqua 2, 229–245 (1979).

252. J. R. Speakman, The function of daylight flying in British bats. J. Zoöl. 220, 101–113 (1990).

253. M. Ilan, Y. Yom-Tov, Diel Activity Pattern of a Diurnal Desert Rodent, Psammomys obesus. J. Mammal. 71, 66–69 (1990).

254. C. Bilu, H. Einat, P. Zimmet, N. Kronfeld-Schor, Circadian rhythms-related disorders in diurnal fat sand rats under modern lifestyle conditions: A review. Front. Physiol. 13, 963449 (2022).

255. C. Leuchtenberger, C. A. Zucco, C. Ribas, W. Magnusson, G. Mourão, Activity patterns of giant otters recorded by telemetry and camera traps. Ethol. Ecol. Evol. 26, 19–28 (2014).

256. P. Beier, D. Choate, R. H. Barrett, Movement Patterns of Mountain Lions during Different Behaviors. J. Mammal. 76, 1056–1070 (1995).

257. B. E. H. van Oort, N. J. C. Tyler, M. P. Gerkema, L. Folkow, A. S. Blix, K.-A. Stokkan, Circadian organization in reindeer. Nature 438, 1095–1096 (2005).

258. S. A. Meier, M. Furrer, N. Nowak, R. Zenobi, M. A. Sundset, R. Huber, S. A. Brown, G. Wagner, Uncoupling of behavioral and metabolic 24-h rhythms in reindeer. Curr. Biol. 34, 1596–1603.e4 (2024).

259. L.-Y. Shuai, C.-L. Ren, C. Cao, Y.-L. Song, Z.-G. Zeng, Shifts in activity patterns of Microtus gregalis: a role of competition or temperature? J. Mammal. 95, 960–967 (2014).

260. F. Johann, M. Handschuh, P. Linderoth, C. F. Dormann, J. Arnold, Adaptation of wild boar (Sus scrofa) activity in a human-dominated landscape. BMC Ecol. 20, 4 (2020).

261. P. Cruz, A. Paviolo, R. F. Bó, J. J. Thompson, M. S. D. Bitetti, Daily activity patterns and habitat use of the lowland tapir (Tapirus terrestris) in the Atlantic Forest. Mamm. Biol. 79, 376– 383 (2014).

262. M. M. Dimanico, A.-L. Klaassen, J. Wang, M. Kaeser, M. Harvey, B. Rasch, G. Rainer, Aspects of tree shrew consolidated sleep structure resemble human sleep. Commun. Biol. 4, 722 (2021).

263. J. H. Meijer, S. Daan, G. J. F. Overkamp, P. M. Hermann, The Two-Oscillator Circadian System of Tree Shrews (Tupaia belangeri) and Its Response to Light and Dark Pulses. J. Biol. Rhythm. 5, 1–16 (1990).

264. J. V. Ware, K. D. Rode, C. T. Robbins, T. Leise, C. R. Weil, H. T. Jansen, The Clock Keeps Ticking: Circadian Rhythms of Free-Ranging Polar Bears. J. Biol. Rhythm. 35, 180–194 (2020).

265. P. T. Matthews, J. Barwick, A. K. Doughty, E. K. Doyle, C. L. Morton, W. Y. Brown, Alpaca Field Behaviour When Cohabitating with Lambing Ewes. Animals 10, 1605 (2020).

266. M. Scantlebury, M. Danek-Gontard, P. W. Bateman, N. C. Bennett, M. B. Manjerovic, M.-B. Manjerovic, K. E. Joubert, J. M. Waterman, Seasonal Patterns of Body Temperature Daily Rhythms in Group-Living Cape Ground Squirrels Xerus inauris. PLoS ONE 7, e36053 (2012).

