## Supplementary material for "A cellular basis for the mammalian nocturnal-diurnal switch": Table S2

Diurnal and nocturnal activity scoring for species included in the analysis of evolutionary rate.

| assembly | code | species | Order | Family | common name | Phenotype | Reference |
| --- | --- | --- | --- | --- | --- | --- | --- |
| vs_HLaciJub2 | aciJub2 | Acinonyx_jubatus | Carnivora | Felidae | Cheetah | DIU | (*195*, *196*) |
| vs_HLacoCah1 | acoCah1 | Acomys_cahirinus | Rodentia | Muridae | cairo spiny mouse | NOC | (*197*) |
| vs_HLailMel2 | ailMel2 | Ailuropoda_melanoleuca | Carnivora | Ursidae | giant panda | CAT | (*198*) |
| vs_HLailFul2 | ailFul2 | Ailurus_fulgens | Carnivora | Ailuridae | lesser panda | NOC | (*199*) |
| vs_HLallBul1 | allBul1 | Allactaga_bullata | Rodentia | Dipodidae | gobi jerboa | NOC | (*200*) |
| vs_HLaloPal1 | aloPal1 | Alouatta_palliata | Primates | Atelidae | mantled howler | DIU | (*201*) |
| vs_HLammLer1 | ammLer1 | Ammotragus_lervia | Cetartiodactyla | Bovidae | aoudad | CRE | (*202*) |
| vs_HLanoCau1 | anoCau1 | Anoura_caudifer | Chiroptera | Phyllostomidae | tailed tailless bat | NOC | (*203*) |
| vs_HLantAme1 | antAme1 | Antilocapra_americana | Cetartiodactyla | Antilocapridae | pronghorn | DIU | (*204*) |
| vs_aotNan1 | aotNan1 | Aotus_nancymaae | Primates | Aotidae | nancy ma’a night monkey | NOC | (*205*) |
| vs_HLaplRuf1 | aplRuf1 | Aplodontia_rufa | Rodentia | Aplodontiidae | mountain beaver | NOC | (*206*) |
| vs_HLartJam2 | artJam2 | Artibeus_jamaicensis | Chiroptera | Phyllostomidae | jamacian fruit-eating bat | NOC | (*206*) |
| vs_HLateGeo1 | ateGeo1 | Ateles_geoffroyi | Primates | Atelidae | geoffroy's spider monkey | DIU | (*207*) |
| vs_HLbeaHun1 | beaHun1 | Beatragus_hunteri | Cetartiodactyla | Bovidae | hirola | CRE | (*208*) |
| vs_bisBis1 | bisBis1 | Bison_bison_bison | Cetartiodactyla | Bovidae | american bison | CAT | (*200*) |
| vs_HLbosInd2 | bosInd2 | Bos_indicus | Cetartiodactyla | Bovidae | zebu cattle | CAT | (*200*) |
| vs_HLbosMut2 | bosMut2 | Bos_mutus | Cetartiodactyla | Bovidae | wild yak | DIU | (*209*) |
| vs_bosTau9 | bosTau9 | Bos_taurus | Cetartiodactyla | Bovidae | cattle | DIU | (*210*) |
| vs_HLbubBub2 | bubBub2 | Bubalus_bubalis | Cetartiodactyla | Bovidae | water buffalo | CAT | (*211*) |
| vs_HLcalJac4 | calJac4 | Callithrix_jacchus | Primates | Cebidae | common marmoset | DIU | (*212*) |
| vs_HLcamBac1 | camBac1 | Camelus_bactrianus | Cetartiodactyla | Camelidae | bactrian camel | DIU | (*213*) |
| vs_HLcamDro2 | camDro2 | Camelus_dromedarius | Cetartiodactyla | Camelidae | arabian camel | DIU | (*214*) |
| vs_HLcamFer3 | camFer3 | Camelus_ferus | Cetartiodactyla | Camelidae | wild bactrian camel | DIU | (*215*, *216*) |
| vs_HLcapAeg1 | capAeg1 | Capra_aegagrus | Cetartiodactyla | Bovidae | wild goat | CRE | (*217*) |
| vs_HLcapHir2 | capHir2 | Capra_hircus | Cetartiodactyla | Bovidae | goat | DIU | (*218*) |
| vs_HLcarPer3 | carPer3 | Carollia_perspicillata | Chiroptera | Phyllostomidae | seba's short-tailed bat | NOC | (*203*) |
| vs_HLcasCan3 | casCan3 | Castor_canadensis | Rodentia | Castoridae | north american beaver | NOC | (*200*) |
| vs_HLcatWag1 | catWag1 | Catagonus_wagneri | Cetartiodactyla | Tayassuidae | chacoan peccary | DIU | (*219*) |
| vs_cavApe1 | cavApe1 | Cavia_aperea | Rodentia | Caviidae | brazilian guinea pig | CAT | (*220*, *221*) |
| vs_HLcavTsc1 | cavTsc1 | Cavia_tschudii | Rodentia | Caviidae | montane guinea pig | NOC | (*222*) |
| vs_HLcebAlb1 | cebAlb1 | Cebus_albifrons | Primates | Cebidae | white-fronted capuchin | DIU | (*63*) |
| vs_cebCap1 | cebCap1 | Cebus_capucinus_imitator | Primates | Cebidae | white-faced sapajou | DIU | (*63*) |
| vs_HLcerSimCot1 | cerSimCot1 | Ceratotherium_simum_cottoni | Perissodactyla | Rhinocerotidae | northern white rhinoceros | CRE | (*200*) |
| vs_cerSim1 | cerSim1 | Ceratotherium_simum_simum | Perissodactyla | Rhinocerotidae | southern white rhinoceros | CRE | (*200*) |
| vs_cerAty1 | cerAty1 | Cercocebus_atys | Primates | Cercopithecidae | sooty mangabey | DIU | (*197*) |
| vs_HLcerNeg1 | cerNeg1 | Cercopithecus_neglectus | Primates | Cercopithecidae | de brazza's monkey | DIU | (*63*) |
| vs_HLcheMed1 | cheMed1 | Cheirogaleus_medius | Primates | Cheirogaleidae | fat-tailed dwarf lemur | NOC | (*63*) |
| vs_chiLan1 | chiLan1 | Chinchilla_lanigera | Rodentia | Chinchillidae | long-tailed chinchilla | NOC | (*206*) |
| vs_chlSab2 | chlSab2 | Chlorocebus_sabaeus | Primates | Cercopithecidae | green monkey | DIU | (*63*) |
| vs_colAng1 | colAng1 | Colobus_angolensis_palliatus | Primates | Cercopithecidae | angolan colobus | DIU | (*63*) |
| vs_conCri1 | conCri1 | Condylura_cristata | Eulipotyphla | Talpidae | star-nosed mole | CAT | (*200*) |
| vs_HLcraTho1 | craTho1 | Craseonycteris_thonglongyai | Chiroptera | Craseonycteridae | bumblebee bat | NOC | (*200*) |
| vs_HLcriGam1 | criGam1 | Cricetomys_gambianus | Rodentia | Nesomyidae | gambian pouched rat | NOC | (*203*) |
| vs_HLcryFer2 | cryFer2 | Cryptoprocta_ferox | Carnivora | Eupleridae | fossa | CAT | (*198*) |
| vs_HLcteGun1 | cteGun1 | Ctenodactylus_gundi | Rodentia | Ctenodactylidae | common gundi | DIU | (*208*) |
| vs_HLdasPun1 | dasPun1 | Dasyprocta_punctata | Rodentia | Dasyproctidae | central american agouti | DIU | (*223*) |
| vs_HLdauMad1 | dauMad1 | Daubentonia_madagascariensis | Primates | Daubentoniidae | aye-aye | NOC | (*200*) |
| vs_HLdesRot2 | desRot2 | Desmodus_rotundus | Chiroptera | Phyllostomidae | common vampire bat | NOC | (*203*) |
| vs_HLdicSum1 | dicSum1 | Dicerorhinus_sumatrensis_sumatrensis | Perissodactyla | Rhinocerotidae | sumatran rhinoceros | NOC | (*200*) |
| vs_HLdicBic1 | dicBic1 | Diceros_bicornis | Perissodactyla | Rhinocerotidae | black rhinoceros | NOC | (*200*) |
| vs_HLdinBra1 | dinBra1 | Dinomys_branickii | Rodentia | Dinomyidae | pacarana | NOC | (*200*) |
| vs_dipOrd2 | dipOrd2 | Dipodomys_ordii | Rodentia | Heteromyidae | ord's kangaroo rat | NOC | (*197*) |
| vs_HLdipSte1 | dipSte1 | Dipodomys_stephensi | Rodentia | Heteromyidae | stephen's kangaroo rat | NOC | (*200*) |
| vs_HLdolPat1 | dolPat1 | Dolichotis_patagonum | Rodentia | Caviidae | patagonian mara | DIU | (*224*) |
| vs_HLeidHel2 | eidHel2 | Eidolon_helvum | Chiroptera | Pteropodidae | straw-colored fruit bat | NOC | (*200*) |
| vs_HLelaDav1 | elaDav1 | Elaphurus_davidianus | Cetartiodactyla | Cervidae | pere david's deer | DIU | (*211*) |
| vs_enhLutNer1 | enhLutNer1 | Enhydra_lutris_kenyoni | Carnivora | Mustelidae | sea otter | DIU | (*225*) |
| vs_eptFus1 | eptFus1 | Eptesicus_fuscus | Chiroptera | Vespertilionidae | big brown bat | NOC | (*206*) |
| vs_equCab3 | equCab3 | Equus_caballus | Perissodactyla | Equidae | horse | CAT | (*226*, *227*) |
| vs_equPrz1 | equPrz1 | Equus_przewalskii | Perissodactyla | Equidae | przewalski's horse | DIU | (*228*) |
| vs_eriEur2 | eriEur2 | Erinaceus_europaeus | Eulipotyphla | Erinaceidae | western european hedgehog | NOC | (*200*) |
| vs_HLeryPat1 | eryPat1 | Erythrocebus_patas | Primates | Cercopithecidae | patas monkey | DIU | (*63*) |
| vs_HLeulFla1 | eulFla1 | Eulemur_flavifrons | Primates | Lemuridae | sclater's lemur | CAT | (*229*) |
| vs_HLeulFul1 | eulFul1 | Eulemur_fulvus | Primates | Lemuridae | common brown lemur | CAT | (*63*, *230*) |
| vs_felCat9 | felCat9 | Felis_catus | Carnivora | Felidae | domestic cat | CAT | (*208*) |
| vs_HLfelNig1 | felNig1 | Felis_nigripes | Carnivora | Felidae | black-footed cat | NOC | (*198*) |
| vs_HLfukDam2 | fukDam2 | Fukomys_damarensis | Rodentia | Bathyergidae | damara mole-rat | CAT | (*231*) |
| vs_HLgirTip1 | girTip1 | Giraffa_tippelskirchi | Cetartiodactyla | Giraffidae | giraffe | CRE | (*200*) |
| vs_HLgliGli1 | gliGli1 | Glis_glis | Rodentia | Gliridae | edible dormouse | NOC | (*208*) |
| vs_gorGor6 | gorGor6 | Gorilla_gorilla_gorilla | Primates | Hominidae | western lowland gorilla | DIU | (*63*) |
| vs_HLgraMur1 | graMur1 | Graphiurus_murinus | Rodentia | Gliridae | woodland doormouse | NOC | (*200*) |
| vs_HLhelPar1 | helPar1 | Helogale_parvula | Carnivora | Herpestidae | dwarf mongoose | DIU | (*232*) |
| vs_HLhemHyl1 | hemHyl1 | Hemitragus_hylocrius | Cetartiodactyla | Bovidae | nilgiri tahr | NOC | (*200*) |
| vs_HLhipArm1 | hipArm1 | Hipposideros_armiger | Chiroptera | Hipposideridae | great roundleaf bat | NOC | (*233*) |
| vs_HLhipGal1 | hipGal1 | Hipposideros_galeritus | Chiroptera | Hipposideridae | cantor's leaf-nosed bat | NOC | (*203*) |
| vs_HLhyaHya1 | hyaHya1 | Hyaena_hyaena | Carnivora | Hyaenidae | striped hyena | NOC | (*198*) |
| vs_HLhydHyd1 | hydHyd1 | Hydrochoerus_hydrochaeris | Rodentia | Caviidae | capybara | DIU | (*234*) |
| vs_HLhysCri1 | hysCri1 | Hystrix_cristata | Rodentia | Hystricidae | crested porcupine | NOC | (*200*) |
| vs_HLindInd1 | indInd1 | Indri_indri | Primates | Indridae | indri | DIU | (*63*) |
| vs_jacJac1 | jacJac1 | Jaculus_jaculus | Rodentia | Dipodidae | lesser egyptian jerboa | NOC | (*206*) |
| vs_HLlasBor1 | lasBor1 | Lasiurus_borealis | Chiroptera | Vespertilionidae | eastern red bat | NOC | (*203*) |
| vs_HLlemCat1 | lemCat1 | Lemur_catta | Primates | Lemuridae | ring tailed lemur | CAT | (*63*) |
| vs_HLlepAme1 | lepAme1 | Lepus_americanus | Lagomorpha | Leporidae | snowshoe hare | NOC | (*197*) |
| vs_HLlycPic3 | lycPic3 | Lycaon_pictus | Carnivora | Canidae | african hunting dog | CRE | (*198*, *235*) |
| vs_HLmacFas6 | macFas6 | Macaca_fascicularis | Primates | Cercopithecidae | crab-eating macaque | DIU | (*63*) |
| vs_macNem1 | macNem1 | Macaca_nemestrina | Primates | Cercopithecidae | southern pig-tailed macaque | DIU | (*63*) |
| vs_HLmacSob1 | macSob1 | Macroglossus_sobrinus | Chiroptera | Pteropodidae | long-tongued fruit bat | NOC | (*233*) |
| vs_manLeu1 | manLeu1 | Mandrillus_leucophaeus | Primates | Cercopithecidae | drill | DIU | (*63*) |
| vs_HLmanJav2 | manJav2 | Manis_javanica | Pholidota | Manidae | malayan pangolin | NOC | (*203*) |
| vs_HLmanPen2 | manPen2 | Manis_pentadactyla | Pholidota | Manidae | chinese pangolin | NOC | (*206*) |
| vs_HLmarMar1 | marMar1 | Marmota_marmota_marmota | Rodentia | Sciuridae | alpine marmot | DIU | (*236*) |
| vs_HLmegLyr2 | megLyr2 | Megaderma_lyra | Chiroptera | Megadermatidae | greater false vampire bat | NOC | (*200*) |
| vs_HLmelCap1 | melCap1 | Mellivora_capensis | Carnivora | Mustelidae | honey badger | NOC | (*198*) |
| vs_HLmerUng1 | merUng1 | Meriones_unguiculatus | Rodentia | Muridae | mongolian jird | CAT | (*200*) |
| vs_mesAur1 | mesAur1 | Mesocricetus_auratus | Rodentia | Cricetidae | golden hamster | NOC | (*237*) |
| vs_micMur3 | micMur3 | Microcebus_murinus | Primates | Cheirogaleidae | gray mouse lemur | NOC | (*238*) |
| vs_HLmicHir1 | micHir1 | Micronycteris_hirsuta | Chiroptera | Phyllostomidae | hairy big-eared bat | NOC | (*200*) |
| vs_micOch1 | micOch1 | Microtus_ochrogaster | Rodentia | Cricetidae | prairie vole | CAT | (*239*) |
| vs_HLminNat1 | minNat1 | Miniopterus_natalensis | Chiroptera | Vespertilionidae | atal long-fingered bat | NOC | Chiroptera - assumed nocturnal unless otherwise described |
| vs_HLminSch1 | minSch1 | Miniopterus_schreibersii | Chiroptera | Vespertilionidae | common bent-wing bat | NOC | Chiroptera - assumed nocturnal unless otherwise described |
| vs_HLmirCoq1 | mirCoq1 | Mirza_coquereli | Primates | Cheirogaleidae | coquerel's giant mouse lemur | NOC | (*63*) |
| vs_HLmorBla1 | morBla1 | Mormoops_blainvillei | Chiroptera | Mormoopidae | ghost-faced bat | NOC | (*203*) |
| vs_HLmosMos1 | mosMos1 | Moschus_moschiferus | Cetartiodactyla | Moschidae | siberian musk deer | NOC | (*211*) |
| vs_HLmunMug1 | munMug1 | Mungos_mungo | Carnivora | Herpestidae | south african banded mongoose | DIU | (*240*) |
| vs_HLmurAurFea1 | murAurFea1 | Murina_aurata_feae | Chiroptera | Vespertilionidae | ashy-gray tube-nosed bat | NOC | (*200*) |
| vs_HLmusCar1 | musCar1 | Mus_caroli | Rodentia | Muridae | ryukyu mouse | NOC | (*208*) |
| vs_mm39 | mm39 | Mus_musculus | Rodentia | Muridae | house mouse | NOC | (*208*) |
| vs_HLmusPah1 | musPah1 | Mus_pahari | Rodentia | Muridae | shrew mouse | NOC | (*208*) |
| vs_HLmusSpr1 | musSpr1 | Mus_spretus | Rodentia | Muridae | western mediterranean mouse | NOC | (*200*) |
| vs_HLmusAve1 | musAve1 | Muscardinus_avellanarius | Rodentia | Gliridae | hazel dormouse | NOC | (*208*) |
| vs_HLmusFur2 | musFur2 | Mustela_putorius_furo | Carnivora | Mustelidae | domestic ferret | NOC | (*198*) |
| vs_HLmyoCoy1 | myoCoy1 | Myocastor_coypus | Rodentia | Myocastoridae | coypu | NOC | (*206*) |
| vs_myoBra1 | myoBra1 | Myotis_brandtii | Chiroptera | Vespertilionidae | brandt's bat | NOC | (*200*) |
| vs_myoDav1 | myoDav1 | Myotis_davidii | Chiroptera | Vespertilionidae | david's myotis | NOC | (*241*) |
| vs_HLmyoLuc1 | myoLuc1 | Myotis_lucifugus | Chiroptera | Vespertilionidae | little brown bat | NOC | (*206*) |
| vs_HLmyoMyo6 | myoMyo6 | Myotis_myotis | Chiroptera | Vespertilionidae | greater mouse-eared bat | NOC | (*203*) |
| vs_nasLar1 | nasLar1 | Nasalis_larvatus | Primates | Cercopithecidae | proboscis monkey | DIU | (*63*) |
| vs_HLnocLep1 | nocLep1 | Noctilio_leporinus | Chiroptera | Noctilionidae | greater bulldog bat | NOC | (*208*) |
| vs_HLnomLeu4 | nomLeu4 | Nomascus_leucogenys | Primates | Hylobatidae | northern white-cheeked gibbon | DIU | (*63*) |
| vs_HLnycCou1 | nycCou1 | Nycticebus_coucang | Primates | Lorisidae | sunda slow loris | NOC | (*63*, *242*) |
| vs_ochPri3 | ochPri3 | Ochotona_princeps | Lagomorpha | Ochotonidae | american pika | DIU | (*243*) |
| vs_octDeg1 | octDeg1 | Octodon_degus | Rodentia | Octodontidae | degu | DIU | (*244*) |
| vs_HLodoVir3 | odoVir3 | Odocoileus_virginianus_texanus | Cetartiodactyla | Cervidae | white-tailed deer | CRE | (*211*) |
| vs_HLokaJoh2 | okaJoh2 | Okapia_johnstoni | Cetartiodactyla | Giraffidae | okapi | DIU | (*245*) |
| vs_HLondZib1 | ondZib1 | Ondatra_zibethicus | Rodentia | Cricetidae | muskrat | NOC | (*200*) |
| vs_HLonyTor1 | onyTor1 | Onychomys_torridus | Rodentia | Cricetidae | scorpion mouse | NOC | (*200*) |
| vs_HLoryCunCun4 | oryCunCun4 | Oryctolagus_cuniculus_cuniculus | Lagomorpha | Leporidae | rabbit | CAT | (*208*) |
| vs_otoGar3 | otoGar3 | Otolemur_garnettii | Primates | Galagidae | small-eared galago | NOC | (*63*) |
| vs_HLoviAri5 | oviAri5 | Ovis_aries | Cetartiodactyla | Bovidae | sheep | DIU | (*246*) |
| vs_HLoviCan1 | oviCan1 | Ovis_canadensis_canadensis | Cetartiodactyla | Bovidae | peninsular bighorn sheep | CRE | (*247*, *248*) |
| vs_panPan3 | panPan3 | Pan_paniscus | Primates | Hominidae | bonobo | DIU | (*63*) |
| vs_panTro6 | panTro6 | Pan_troglodytes | Primates | Hominidae | chimpanzee | DIU | (*63*) |
| vs_HLpanOnc2 | panOnc2 | Panthera_onca | Carnivora | Felidae | jaguar | CAT | (*198*) |
| vs_HLpanPar1 | panPar1 | Panthera_pardus | Carnivora | Felidae | leopard | NOC | (*198*) |
| vs_panTig1 | panTig1 | Panthera_tigris_altaica | Carnivora | Felidae | amur tiger | CRE | (*249*) |
| vs_panHod1 | panHod1 | Pantholops_hodgsonii | Cetartiodactyla | Bovidae | chiru | DIU | (*250*) |
| vs_HLpapAnu5 | papAnu5 | Papio_anubis | Primates | Cercopithecidae | olive baboon | DIU | (*63*) |
| vs_HLparHer1 | parHer1 | Paradoxurus_hermaphroditus | Carnivora | Viverridae | asian palm civet | NOC | (*203*) |
| vs_HLperLonPac1 | perLonPac1 | Perognathus_longimembris_pacificus | Rodentia | Heteromyidae | pacific pocket mouse | NOC | (*200*) |
| vs_HLperManBai2 | perManBai2 | Peromyscus_maniculatus_bairdii | Rodentia | Cricetidae | prairie deer mouse | NOC | (*200*) |
| vs_HLpetTyp1 | petTyp1 | Petromus_typicus | Rodentia | Petromuridae | dassie rat | DIU | (*251*) |
| vs_HLpilTep2 | pilTep2 | Piliocolobus_tephrosceles | Primates | Cercopithecidae | ugandan red colobus | DIU | (*63*) |
| vs_HLpipPip2 | pipPip2 | Pipistrellus_pipistrellus | Chiroptera | Vespertilionidae | common pipistrelle | NOC | (*252*) |
| vs_HLpitPit1 | pitPit1 | Pithecia_pithecia | Primates | Pitheciidae | white-faced saki | DIU | (*63*) |
| vs_HLpleDon1 | pleDon1 | Plecturocebus_donacophilus | Primates | Pitheciidae | white-eared titi | DIU | (*63*) |
| vs_ponAbe3 | ponAbe3 | Pongo_abelii | Primates | Hominidae | sumatran orangutan | DIU | (*63*) |
| vs_proCoq1 | proCoq1 | Propithecus_coquereli | Primates | Indridae | coquerel's sifaka | DIU | (*63*) |
| vs_HLpsaObe1 | psaObe1 | Psammomys_obesus | Rodentia | Muridae | fat sand rat | DIU | (*253*, *254*) |
| vs_ptePar1 | ptePar1 | Pteronotus_parnellii | Chiroptera | Mormoopidae | parnell's mustached bat | NOC | (*206*) |
| vs_HLpteBra2 | pteBra2 | Pteronura_brasiliensis | Carnivora | Mustelidae | giant otter | DIU | (*255*) |
| vs_pteAle1 | pteAle1 | Pteropus_alecto | Chiroptera | Pteropodidae | black flying fox | NOC | (*208*) |
| vs_HLpteVam2 | pteVam2 | Pteropus_vampyrus | Chiroptera | Pteropodidae | large flying fox | NOC | (*203*) |
| vs_HLpumCon1 | pumCon1 | Puma_concolor | Carnivora | Felidae | puma | CRE | (*256*) |
| vs_HLpygNem1 | pygNem1 | Pygathrix_nemaeus | Primates | Cercopithecidae | red-shanked douc langur | DIU | (*63*) |
| vs_HLranTar1 | ranTar1 | Rangifer_tarandus | Cetartiodactyla | Cervidae | siberian reindeer | CAT | (*257*, *258*) |
| vs_HLratNor7 | ratNor7 | Rattus_norvegicus | Rodentia | Muridae | brown rat | NOC | (*203*) |
| vs_HLrhiSin1 | rhiSin1 | Rhinolophus_sinicus | Chiroptera | Rhinolophidae | chinese rufous horseshoe bat | NOC | (*233*) |
| vs_rhiBie1 | rhiBie1 | Rhinopithecus_bieti | Primates | Cercopithecidae | black snub-nosed monkey | DIU | (*63*) |
| vs_HLrhiRox2 | rhiRox2 | Rhinopithecus_roxellana | Primates | Cercopithecidae | golden snub-nosed monkey | DIU | (*63*) |
| vs_HLrouAeg4 | rouAeg4 | Rousettus_aegyptiacus | Chiroptera | Pteropodidae | egyptian fruit bat | NOC | (*206*) |
| vs_HLsagImp1 | sagImp1 | Saguinus_imperator | Primates | Cebidae | emperor tamarin | DIU | (*63*) |
| vs_HLsaiTat1 | saiTat1 | Saiga_tatarica | Cetartiodactyla | Bovidae | steppe saiga | DIU | (*200*) |
| vs_HLsaiBol1 | saiBol1 | Saimiri_boliviensis_boliviensis | Primates | Cebidae | bolivian squirrel monkey | DIU | (*63*) |
| vs_HLscaAqu1 | scaAqu1 | Scalopus_aquaticus | Eulipotyphla | Talpidae | eastern mole | CAT | (*200*) |
| vs_HLsemEnt1 | semEnt1 | Semnopithecus_entellus | Primates | Cercopithecidae | northern plains gray langur | DIU | (*63*) |
| vs_HLsigHis1 | sigHis1 | Sigmodon_hispidus | Rodentia | Cricetidae | hispid cotton rat | CAT | (*200*) |
| vs_HLsolPar1 | solPar1 | Solenodon_paradoxus | Eulipotyphla | Solenodontidae | hispaniolan solenodon | NOC | (*200*) |
| vs_sorAra2 | sorAra2 | Sorex_araneus | Eulipotyphla | Soricidae | common shrew | CAT | (*200*) |
| vs_HLspeDau1 | speDau1 | Spermophilus_dauricus | Rodentia | Sciuridae | daurian ground squirrel | DIU | (*259*) |
| vs_HLspiGra1 | spiGra1 | Spilogale_gracilis | Carnivora | Mephitidae | western spotted skunk | NOC | (*198*) |
| vs_HLsurSur2 | surSur2 | Suricata_suricatta | Carnivora | Herpestidae | meerkat | DIU | (*211*) |
| vs_susScr11 | susScr11 | Sus_scrofa | Cetartiodactyla | Suidae | pig | CAT | (*260*) |
| vs_HLtadBra1 | tadBra1 | Tadarida_brasiliensis | Chiroptera | Molossidae | mexican free-tailed bat | NOC | (*206*) |
| vs_HLtapInd2 | tapInd2 | Tapirus_indicus | Perissodactyla | Tapiridae | malayan tapir | NOC | (*211*) |
| vs_HLtapTer1 | tapTer1 | Tapirus_terrestris | Perissodactyla | Tapiridae | south american tapir | NOC | (*261*) |
| vs_HLthrSwi1 | thrSwi1 | Thryonomys_swinderianus | Rodentia | Thryonomyidae | greater cane rat | NOC | (*200*) |
| vs_HLtonSau1 | tonSau1 | Tonatia_saurophila | Chiroptera | Phyllostomidae | stripe-headed round-eared bat | NOC | (*200*) |
| vs_HLtraJav1 | traJav1 | Tragulus_javanicus | Cetartiodactyla | Tragulidae | java lesser chevrotain | NOC | (*200*) |
| vs_tupChi1 | tupChi1 | Tupaia_chinensis | Scandentia | Tupaiidae | northern tree shrew | DIU | (*262*) |
| vs_tupBel1 | tupBel1 | Tupaia_tana | Scandentia | Tupaiidae | large treeshrew | DIU | (*263*) |
| vs_HLuroGra1 | uroGra1 | Uropsilus_gracilis | Eulipotyphla | Talpidae | gracile shrew mole | NOC | (*208*) |
| vs_ursMar1 | ursMar1 | Ursus_maritimus | Carnivora | Ursidae | polar bear | DIU | (*264*) |
| vs_vicPac2 | vicPac2 | Vicugna_pacos_huacaya | Cetartiodactyla | Camelidae | alpaca | DIU | (*265*) |
| vs_HLvulLag1 | vulLag1 | Vulpes_lagopus | Carnivora | Canidae | arctic fox | NOC | (*203*) |
| vs_HLxerIna1 | xerIna1 | Xerus_inauris | Rodentia | Sciuridae | cape ground squirrel | DIU | (*266*) |
| vs_HLzapHud1 | zapHud1 | Zapus_hudsonius | Rodentia | Dipodidae | meadow jumping mouse | NOC | (*237*) |
