## Supplementary Materials for "A cellular basis for the mammalian nocturnal-diurnal switch"

**The PDF file includes:**

Materials and Methods
Figs. S1 to S11

**Other Supplementary Materials for this manuscript include the following:**

Tables S1 to S3 as a separate files

### Materials and Methods

Cell culture

Primary fibroblasts from mice (*Mus musculus*, wild-type C57BL/6), humans (*Homo sapiens*), common marmoset (*Callithrix jacchus*), ring tailed lemur (*Lemur catta*), common sheep (*Ovis aries*), rat (*Rattus norvegicus*), and four-striped grass mouse (*Rhabdomys pumilio*) were obtained as following. Tissue biopsies were taken from skin (human, mouse, rat and striped mouse), lung (mouse), heart (lemur, sheep) or muscle (marmoset, gibbon), and primary fibroblasts were isolated following standard culture (*161*). Adult dermal fibroblasts were obtained from healthy volunteers at the John van Geest Centre for Brain Repair (Cambridge, UK), under local ethical approval (REC 09/H0311/88) using a previously described method (*162*). Fibroblasts were kept in Dulbecco's modified Eagle medium (DMEM) high glucose, GlutaMAX(TM), pyruvate medium (Gibco, #31966) supplemented with 10% fetal bovine serum (FBS, Gibco, #10437028) or HyClone FetalClone III serum (Cytiva Life Sciences, #SH30109) and 1% penicillin/streptomycin (P/S, final concentration: 100 units/mL of penicillin and 100 µg/mL of streptomycin, Gibco, #15140122) and passaged when they reached 80–90% confluency. All lines were cultured to at least passage 2, tested for mycoplasma using MycoAlert™ mycoplasma detection kit (Lonza, #LT07), and maintained for routine culture in DMEM supplemented with 10-20% FetalClone III serum and P/S at 37°C, 5 % $O_2$, 5% $CO_2$. For creation of bioluminescent lines, cells were transduced with *pLV6-Bmal-luc* lentivirus (gift of Steven Brown, also found at Addgene plasmid #68833) and selected for stable cultures by blasticidin (blasticidin-HCL, Gibco) selection at 4 µg/ml. Selection was performed for between 1 and 2 weeks until non-transduced control cells were dead, and colonies were established in transduced lines. Lines were subsequently maintained at 2 µg/ml blasticidin in DMEM containing 10% FetalClone III serum and penicillin/streptomycin at 37°C, 5% $CO_2$. Experiments were performed at between 1 and 5 passages post transduction, omitting blasticidin in the experimental culture medium.

Immortalized lung fibroblasts were obtained from lung tissue of adult PER2-LUC mice (*67*) and immortalized by serial passage as described in Crosby et al. (*29*). Immortalized fibroblast lines were cultured as described previously (*163*), cultured in DMEM/10% FetalClone III/P/S as above for primary lines, but maintained at atmospheric $O_2$.

Human U2OS (also known as U-2 OS) cells, purchased from ATCC, were cultured in DMEM (Gibco, #31966) supplemented with 10% HyClone FetalClone II serum (Cytiva Life Sciences, #SH30066) and P/S at 5% $CO_2$. For creation of stably transfected lines, cells were transfected with linearized plasmid DNA with Genejuice (Merck, #70967) at 3 µl/µg DNA according to manufacturer's protocol. Between 24 and 48 h after transfection, cells were selected for stable transfection by supplementing medium with puromycin (puromycin dihydrochloride, Gibco, #12122530) at 4 µg/ml.

U2OS cells endogenously expressing the PER2-LUCIFERASE or PER2-Halotag fusion proteins were obtained as following. Guide RNAs for Cas9 editing at the stop codon of *PER2* were designed using a combination of CHOPCHOP v2 (*164*) and ATUM gRNA design tool (atum.bio/eCommerce/cas9/input) and default parameters. Guide RNA sequences, lacking the PAM sequence were inserted as annealed complementary single-stranded oligos into BbsI-digested pSpCas9-2A-GFP (PX458, Addgene). HDR templates, inserting the coding region lacking the initial methionine codon of human codon-optimized firefly *luciferase* (fig. S2A) or human codon-optimized *HaloTag* (Promega, fig. S6A) immediately 5' of the endogenous stop codon, were created by NEB Hifi assembly or direct gene synthesis respectively. Homology arms of ~1000 bp were included either side of the insert coding sequence. U2OS cells, purchased from ATCC, were transfected with a 3:1 molar ratio of linearized HDR template:Cas9 sgRNA using PEI at 3ug/ug DNA. On the third day after transfection, GFP+ve cells were single-cell sorted and expanded as clonal cell lines in 96-well plates. Clonal lines were expanded until confluent, changed into medium containing 1mM firefly luciferin (Biosynth, #L-8220) and placed in an ALLIGATOR Luminescence System ("Alligator", Cairn) at 37°C and 5% $CO_2$ for screening of positive colonies (*165*, *166*). Positive clones were identified by presence of luminescence, which oscillated in brightness with a period of approximately 24 h in constant conditions. Positive clones were validated by PCR (fig. S2B) and functional assays (fig. S2, C to E).

Plasmids

Constitutive expression constructs for mouse PER2-LUC, human PER2-LUC and human PER2_codon optimized-LUC (PER2_CO) were generated as following. PER2 ORF from mouse (GenBank NM_011066.3) or human (NM_022817) was synthesized by Twist Bioscience gene synthesis. Codon optimized human PER2 was synthesized by Twist Bioscience gene synthesis, and codons were optimized using Twist Bioscience Codon Optimization tool. This tool limits the selection of codons for an amino acid with a cutoff frequency of 8%. Codon usage frequency was assessed using Codon Plot from the Sequence Manipulation Suite (https://www.bioinformatics.org/sms2/codon_plot.html, (*167*)) using codon frequencies for Homo sapiens from the Codon Usage Database (https://www.kazusa.or.jp/codon/, (*168*)). mRNA structure was assessed for the short translational ramp of 30 amino acids (*79*) using RNAFold (*169*) web server (http://rna.tbi.univie.ac.at//cgi-bin/RNAWebSuite/RNAfold.cgi) using minimum free energy (MFE) function settings. ORFs were synthesised flanked with SpeI and NotI sites and with no stop codon. ORF-LUC fusions were generated in a step wise subcloning protocol, following MXS chaining assembly protocol (*170*) as following. ORFs were cloned into a backbone containing SV40 promoter. Luciferase, plus SV40pA sequence, was subcloned into this backbone with NotI and SalI to generate an in-frame fusion of ORF and luciferase, under the control of SV40 promoter, in a vector containing puromycin resistance. Constitutive LUC was generated similarly, omitting the ORF subcloning step.

Experimental cell culture - static

For static culture bioluminescent experiments, cells were plated at confluence in transparent bottom black 96-well plates (Greiner, µClear #655096) at 100 µl/well in standard culture medium: DMEM, 10% serum (FetalClone II for U2OS, FetalClone III for fibroblasts), 1X P/S. The following day, medium was replaced with experimental culture medium which was identical to the standard culture medium except for the addition of D-luciferin potassium salt (Biosynth, #L-8220) at 0.037 mM to 1 mM (to deconvolve luciferase signal into steady state and nascent (*69*) , and serum concentrations from 0% to 10%, both as described in figure legends. For examining the effect of amino acid restriction on cellular rhythms, DMEM (Gibco, #31966) was diluted to 10% concentration with EBSS (Gibco, #24010), and then supplemented with sodium bicarbonate (final concentration 44 mM), pyruvate (final concentration 1 mM), glucose (final concentration 25 mM), P/S (final concentration 1X) and serum (FetalClone III, final concentration 1%). For bioluminescence recording, luciferin was added to final concentration of

0.3 mM. Osmolarity of experimental medium was measured with the Osmomat 3000 freezing-point-depression osmometer (Gonotec) and adjusted to 330 ± 10 mOsm/kg using NaCl. Osmolarity of media was controlled in all conditions to remain isosmotic before and after treatment, except where change in extracellular osmolarity was the treatment variable. Here, osmolarity was adjusted with sucrose.

Entrainment was performed as described in figure legends and described below. For temperature entrainment, cells were placed in humidified incubators at 5% CO2 which cycled 37°C 12h: 32°C 12h. At the transition from 32 °C to 37 °C or 37°C to 32 °C, plates were transferred isothermally into dark, light-tight, humified and 5% $CO_2$ Alligator at 37 °C or 32°C respectively. Where indicated, temperature entrainment took place in humified and 5% $CO_2$ Alligators (cycling 37°C 12h: 32°C 12h; 37°C 12h: 35.5°C 12h; or 33°C 12h: 28°C 12h) to enable recording of bioluminescence under cycling conditions. In the latter case, cells were allowed to adapt to the lower temperature of 33°C for 5 days before cycling commenced. In cases were entrainment took place in the Alligator, cells were not disturbed at the transition from cycling to constant conditions.

For dexamethasone or serum synchronizations, after plating, the following day medium was replaced with experimental culture medium containing 10% serum ± 100 nM dexamethasone. Serum synchronization and dexamethasone treatment sets mouse and human cells to identical phases (fig. S1A). Cells were then moved isothermally into humified and 5% $CO_2$ Alligator at 37 °C. The effect on phase of acute translation initiation inhibition under constant temperature conditions was determined by treating with a 10X spike of inhibitor (rocaglamide A or Hippuristanol) isothermally at the trough of PER2 expression. Plates were gently mixed before placing back in Alligators. For re-entrainment, cells were given a final medium change containing 10% serum at ZT0, ZT6 or ZT12, where ZT0 indicates the start of the hot cycle. Cells were transferred isothermally into a humified and 5% $CO_2$ Alligator, which was programmed to cycle 37°C 12h: 32°C 12h for 7 days before maintaining constant 37 °C for the remainder of the recording. For re-entrainment, drug treatments were given at the final medium change before entering temperature cycles or constant conditions as indicated. In all cases, bioluminescence was recorded at half-hourly intervals.

The following drugs were used: Apigenin (Cayman, #10010275), H89 (Apex Bio, #B2190), Hippuristanol (lab of Jerry Pelletier, (*171*)), INK128 (Selleckchem, #S2811), LY3214996 (Selleckchem, #8534), Radicicol (Tocris, #1589), Ravoxertinib (Selleckchem, #S7554), Rocaglamide (Sigma, #SML0656), SCH772984 (Selleckchem, #S7101), SR3029 (Medchem Express, #HY100011), TG003 (Apex Bio, #B1431), Torin1 (Selleckchem, #S2827), TTP22 (Tocris, #4432), VER144008 (Selleckchem, #S7751).

Experimental cell culture - flow

For bioluminescent experiments under continuous flow, cells were plated at confluence in transparent glass bottom 6-channel slides (μ-Slide VI 0.5 Glass Bottom, ibidi) in standard culture medium (DMEM Gibco #31966, 10% FetalClone III serum, 1X P/S) supplemented with 0.3 mM luciferin (Biosynth, #L-8220) at 40 μl/channel and placed in humidified incubators at 5% CO2 and 37°C. After cell attachment, channels were topped up with 120 μl/channel of the same medium. After 24-72h, channel slides were connected in series to a pressurized microfluidics flow system (Elveflow OB1 Mk3), with thermal flow rate sensor feedback control (Elveflow MFS 0-80 μl/min), inside an Alligator bioluminescence incubator set to 37°C and 5% $CO_2$.

"Flow medium", a modified version of the static culture experimental medium (DMEM #31966 Gibco, 1% FetalClone III, 1X P/S, 0.5X Mycozap-plus PR Lonza, 0.3mM luciferin), was delivered to cells via a distribution valve (10:1 MUX bidirectional distributer) at 9-11 µl/min, which was programmed to select medium from pressurized microfluidic reservoirs for durations and intervals as indicated in figure legends. FGF2-containing medium contained 20 ng/ml fibroblast growth factor-basic (FGF2; PeproTech, #450-33). Hyperosmotic medium (+ 50 mOsm) contained 50 mM raffinose (D-(+)-Raffinose pentahydrate; Sigma, #R0514). Bioluminescence was monitored throughout.

<u>Animals and organotypic culture</u>

C57BL/6 and PER2-LUC mice (originally supplied by Joe Takahashi, University of Texas Southwestern) were housed in a specified pathogen free barrier facility. For husbandry and non-experimental housing, mice were group-housed with environmental enrichment under 12:12 light:dark cycles with lights on at 7am. For experiments, mice were singly housed in individual cages containing environmental enrichment and equipped with running wheels to enable monitoring throughout the experiment. All animal experiments were licenced under 1986 Home Office Animal Procedures Act (UK) and carried out in accordance with local animal welfare committee guidelines.

For organotypic culture of pituitary, adrenal glands and lungs, adult mice, aged between 8-12 weeks, 8 females, 16 males, were housed in individual cages placed in LD (lights on 10am, lights off 10pm, 12 mice, 4 females, 8 males) or DL (lights on 10pm, lights off 10am, 12 mice, 4 females, 8 males). Mice were euthanized by cervical dislocation and exsanguination at ZT 0.5 (LD) or ZT 23.5 (DL), ensuring that mice did not experience a light-to-dark or dark-to-light transition before euthanizing. Tissues were dissected from the mice in 1X HBSS on ice following Yamazaki et al. (*172*), Abe et al. (*115*), and Yoshikawa et al. (*173*). Whole pituitary, paired adrenal glands and lung tissue were cut into 1-3 mm pieces in HBSS with a surgical knife following Noguchi et al. (*174*). Diced tissue was cultured on Millicell culture membranes (Millipore, #PICMORG50) in 35 mm dishes with 1.2 ml of medium consisting of DMEM (#31966, Gibco), 5% FBS, 100 IU/ml and 100 µg/ml penicillin/streptomycin, 300 µM D-luciferin potassium salt (Biosynth, #L-8220), and 1 µM INK128 (N=6 mice) or DMSO control (N=6 mice).

SCN slices were prepared as previously described (*175*). Briefly, brains were removed from male and female mice aged between 9-10 weeks and transferred to ice-cold dissection medium: GBSS (Sigma, #G9779), containing 5 mg/mL glucose, and 100 nM MK801 (Sigma, #M-107), 3 mM $MgCl_2$ and 0.05 mM APV (Sigma, #A5282) to block excitotoxicity. After trimming, 300 µm coronal brain slices were prepared on a McIlwain tissue chopper. Non-SCN tissue was removed from slices, before they were transferred to Millicell culture membranes (Millipore, #PICMORG50) placed on top of tissue culture medium containing luciferin (50% Eagle's basal medium, 25% EBSS, 25% heat inactivated horse serum (Invitrogen, USA), 5 mg/mL D-glucose (Sigma), 25 µg/mL Penicillin/Streptomycin, 1% GlutaMAX (Invitrogen), 300 µM luciferin) adjusted to pH7.2 and osmolarity 315-320 mOsm, and supplemented with 100 nM MK801, 3 mM $MgCl_2$ and 0.05 mM APV. After 2-4h, slices were transferred into 35 mm dishes containing fresh culture medium excluding MK801, $MgCl_2$ and APV, for experiments.

Once prepared, tissues were placed in constant conditions (37°C, 5% $CO_2$) or temperature cycling (12h:12h, 32°C:37°C) humidified Alligator for 3-7 days before release into constant conditions of 37°C. Bioluminescence was recorded throughout temperature cycling and constant conditions at half-hourly intervals.

Temperature shift experiments

For temperature shift experiments (described in Fig. 3A and fig. S7A), cells were plated at confluence and maintained at 'origin' temperature of either 32°C or 37°C in incubators humified and at 5% $CO_2$ for one week. 24h before temperature switch, at t=-24h, cells were re-synchronized with 100 nM dexamethasone, to ensure all cells were in the same circadian phase. At t=0h, cells were either switched to the 'destination' temperature of 37°C or 32°C respectively, or maintained at the original constant temperature, for four conditions: constant 'high' of 37°C, 'shift up', 'shift down' or constant 'low' of 32°C. Parallel bioluminescent recordings were made in Alligators, humidified and at 5% CO2, at 32°C or 37°C and parallel plates remained in Alligators or switched between them to match the above conditions. Cells were recorded for at least 4 days post shift to allow assessment of phase and period of the circadian rhythm. For phase response curves, 6 x 96-well plates were consecutively shifted between temperatures in Alligators every 4h for 24h.

Puromycin labelling

Puromycin dihydrochloride (#12122530, Gibco) was added directly to cells in culture medium as 10x bolus to a final concentration of 10 μg/ml puromycin. Labelling proceeded for 30min, after which cells were lysed and puromycin detected by Western blotting using anti-puromycin (PMY-2A4-2 from Developmental Studies Hybridoma bank, at 1:1000. Fold change puromycin incorporation was calculated as $^{\text{incorporation in temperature shifted}}/_{\text{incorporation in constant temperature control}}$.

Protein extraction

For each condition, cells were washed twice in isosmotic ice-cold PBS plus phosphatase and protease inhibitors at 1X (PhosSTOP, Roche, #04906837001 and cOmplete, Roche #4693116001,) and then lysed at room temperature in a urea/thiourea buffer (UTS, 7 M urea, 2 M thiourea, 1 % Na deoxycholate, 20 mM Tris-HCl, 5 mM TCEP), including phosphatase and protease inhibitors. Lysis (500 μl for 15 cm dish for (phospho)proteomics, 150 μl for 6-well plate for puromycin immunoblot), proceeded for 30 min at room temperature. The lysis buffer was prepared the day before sampling began and frozen in 1 ml aliquots. After lysis, the cells were scraped, transferred into Protein LoBind tubes (#0030108116, Eppendorf) and sonicated for 3 min with 30s on/ 30s off sonication (Bioruptor Plus, Diagenode). Samples were then spun at 15,000 rpm at 4°C for 20 min and the supernatant was taken to a fresh tube. Supernatant was flash-frozen in liquid nitrogen and stored at −80°C for subsequent analysis and processing.

For tissues, on dry ice, brain and liver were chopped into ~5 mm pieces, weighed, and 100 mg transferred to 2 ml tubes containing CK14 ceramic beads (Bertin Instruments) with 1 ml UTS buffer (7 M urea, 1% sodium deoxycholate, 20 mM tris pH 8, 5 mM TCEP, 5 mM DTT). Tissues were homogenized using a Precellys 24 Tissue Homogenizer (Bertin Instruments) for 3 x 15 s at 5,000 RPM with 30 s breaks. Lysates were centrifuged at 20,000 x g for 1 min at 4°C, sonicated, centrifuged at 20,000 x g for 30 minutes at 4°C, and supernatants collected.

Protein concentration was measured on thawed samples using Pierce 660 nm Protein Assay Kit (#22662, Thermo Scientific) in U-bottom 96-well plates (Costar) using a Tecan Spark 10M microplate reader, diluting samples 1 in 4 with dH2O to reduce sodium deoxycholate to compatible levels.

Proteomics and phosphoproteomics

Protein samples for (phospho)proteomics were processed as Seinkmane et al. (*150*) with slight modifications.

*Digestion*

Protein samples (150 μg) were reduced with 5 mM DTT at 37°C for 45 min and alkylated with 10 mM iodoacetamide in the dark at room temperature for 30 min. Samples were then diluted to 2.7M urea and digested with Lys-C (Promega) for 4 h at 25°C. Next, samples were further diluted to 1.5 M urea and digested with trypsin (Promega) overnight at 30°C. After digestion, samples were acidified with formic acid (FA) to a final concentration of 0.5% to precipitate SDC, mixed by shaking for 1 min and centrifuged at 12000 x *g* for 10 min. The supernatants were transferred to new tubes and desalted using home-made C18 stage tips (3M Empore) packed with porous R3 resin (Thermo Scientific). The stage tips were equilibrated with 80% acetonitrile (MeCN) / 0.5% FA, followed by 0.5% FA. Bound peptides were eluted with 30-80% MeCN / 0.5% FA and lyophilized.

*Tandem mass tag (TMT) labelling*

The lyophilized peptides from each sample were resuspended in 40 μl of 200 mM Hepes, pH 8.5. 20 μl (500 μg) TMTpro 12plex reagent (Thermo Fisher Scientific), reconstituted in anhydrous MeCN according to manufacturer's instructions was added. Peptides from each time point were labelled with a distinct TMT tag for 60 min at room temperature. The labelling reaction was quenched by incubation with 4 μl 5% hydroxylamine for 30 min. Each set of 12 labelled peptides (4 conditions, triplicates) for human and mouse were combined into two single samples, one per species, and partially dried to remove MeCN in a SpeedVac (Savant). Sample was then acidified and centrifuged at 16000 x *g* for 10 min. Supernatant was desalted using the same stage tips method as above. Bound peptides were eluted and lyophilized.

*Titanium dioxide ($TiO_2$) – Enrichment of phosphopeptides*

Phosphopeptides were enriched using $TiO_2$ titansphere-chromatography (GL Science Inc. Japan). Dried peptides were resolubilized in 50% MeCN containing 2 M lactic acid (loading buffer) and incubated with TiO2 beads (1: 5, peptides: TiO2, w/w) that were prewashed with loading buffer. After 30 min, the TiO2 beads were centrifuged at 9000 x *g* for 2 min, the supernatant was transferred to fresh TiO2 beads for a second round of enrichment. After second incubation, TiO2 beads were loaded onto C8 stage tips (3M Empore) and washed twice with loading buffer and once with 50% MeCN/0.5% FA. Phosphopeptides were eluted sequentially with 100 mM $K_2HPO_4$, pH10; 50% MeCN/50 mM $K_2HPO_4$, pH10 and 50% MeCN/0.5%FA. The eluates were acidified, partially dried in a SpeedVac, desalted with C18 stage tip (3M Empore) same as above and lyophilized.

*Off-line high pH reverse-phase peptides fractionation*

200 µg of the labelled peptides were separated on an off-line, high pressure liquid chromatography (HPLC). The experiment was carried out using XBridge BEH130 C18, 5 µm, 2.1 x 150 mm column (Waters), connected to an Ultimate 3000 analytical HPLC (Thermo Fisher Scientific). Peptides were separated with a gradient of 1-90% buffer A and B (A: 5% MeCN, 10 mM ammonium bicarbonate, pH8; B: MeCN, 10 mM ammonium bicarbonate, pH8, [9:1]) for 60 min at a flow rate of 250 µl/min. A total of 54 fractions were collected, which were then combined into 18 fractions and lyophilized. Dried peptides were resuspended in 1% MeCN / 0.5% FA, and desalted using C18 stage tips, ready for mass spectrometry analysis.

*LC-MS/MS*

The fractionated peptides were analyzed by LC-MS/MS using a fully automated Ultimate 3000 RSLC nano System fitted with a 100 µm x 2 cm PepMap100 C18 nano trap column (Thermo Fisher Scientific) and a 75 µm×25 cm, nanoEase M/Z HSS C18 T3 column (Waters). Peptides were separated using a binary gradient consisting of buffer A (0.1% FA) and buffer B (80% MeCN, 0.1% FA) at 300 nL/min, column temperature of 40 $^{0}$C. Eluted peptides were introduced directly via a nanoFlex ion source into an Orbitrap Eclipse mass spectrometer (Thermo Fisher Scientific). The mass spectrometer was operated in real-time database search (RTS) with synchronous-precursor selection (SPS) -MS3 analysis for reporter ion quantification. MS1 spectra were acquired with a resolution of 120K; mass range = 400-1400 m/z; AGC target = 4e5; maximum injection time (MaxIT) = 50 ms; dynamic exclusion was set at 60 s. MS2 analysis were carried out with collision induced dissociation (CID) activation; ion trap detection; AGC = 1e4; MaxIT = 50 ms and isolation window = 0.7 m/z. RTS-SPS was set up to search Uniprot *Mus musculus* or *Homo sapiens* proteomes (2021), with fixed modifications cysteine carbamidomethylation. TMTpro at Lys and peptide N-terminal, and methionine oxidation were set as dynamic modifications. In MS3 scans, the selected precursors were fragmented by higher-energy collision dissociation (HCD) with collision energy = 55% and analyzed using the orbitrap; resolution = 50K; scan range = 110-500 m/z and MaxIT = 150ms. For phosphoproteomcs, the mass spectrometer was operated in DDA mode, performed MS1 scan events using the same parameters as above. MS2 spectra were acquired with a 50K resolution; MaxIT= 200ms; AGC = 1e5; collision energy =35% and isolation width = 0.7 m/z.

*Raw MS data processing*

The acquired raw files from LC-MS/MS were processed using MaxQuant (Cox & Mann, 2008) with the integrated Andromeda search engine (v1.6.17.0). MS/MS spectra were quantified with reporter ion MS3 from TMTpro experiments and searched against UniProt *Mus musculus* or *Homo sapiens* Reviewed (Nov 2020) Fasta databases. Carbamidomethylation of cysteines was set as a fixed modification, while methionine oxidation, protein N-terminal acetylation and phosphorylation (STY) (for phosphoproteomic group only) were set as variable modifications. Protein quantification requirements were set at 1 unique and razor peptide. In the identification tab, second peptides and match between runs were not selected. Other parameters in MaxQuant were set to default values. The MaxQuant output file was then processed with Perseus (v1.6.6.0). Reporter ion intensities for protein group table were uploaded to Perseus. The data was filtered: identifications from the reverse database, only identified by site, potential contaminants were removed. Then all columns with an intensity "less or equal to zero" were converted to "NAN" (not a number) and exported. The MaxQuant output file with phospho (STY) sites table was also processed with Perseus software (v1.6.15.0). The data was filtered: identifications from the reverse database, potential contaminants were removed, and we only considered phosphopeptides with localization probability ≥ 0.75. Then all columns with intensity "less or equal to zero" were converted to "NAN" and exported.

Proteomics data analysis

After the MaxQuant search, all subsequent proteomics data processing and analysis was performed in R (v4.3.2) with R Studio (2023.09.1+494). The custom scripts are available via a github repository, at https://github.com/andrewbeale. Peptide level information from MaxQuant) was used as a starting point. Contaminants and reverse hits were removed. Sample loading normalization was performed on proteomic and phosphoproteomic data, applying a scaling factor to equalize total summed intensity across individual TMT channels for the proteome, and using the same scaling factor for the phosphoproteome. In this way, real and biological differences in total phosphorylation between samples was maintained, while correcting for sample loading. Peptides and phosphopeptides were filtered to leave only those detected in all conditions. To assess the proteins and phosphopeptides that changed significantly between conditions, the data were processed in the following manner. For temperature steps, the raw intensity data were converted to fold change (FC) by matched replicate (i.e. step up FC = intensity at 32 °C to 37 °C "up"/ intensity at 32 °C "low"; step down FC = intensity at 37 °C to 32 °C/ intensity at 37 °C). These fold changes were $\log_2$ transformed (upfc*#rep*_log2 or downfc*#rep*_log2) before performing repeated measures two-way ANOVA (TWA) on matched pairs of conditions. For adaptation comparisons (i.e. 32 °C vs 37 °C), raw data were $\log_2$ transformed before performing repeated measures two-way ANOVA (TWA) on matched pairs of conditions. Multiple comparisons were corrected for by controlling the false discovery rate by the two-stage step-up method of Benjamini, Krieger and Yekutieli, with $Q = 0.01$. Figures show mean $\log_2$ fold changes, plotted for each protein/phosphoprotein. The hypothesis that mTOR- and WNK-related phosphosites were differentially responsive to acute temperature change or adapted temperature change in the two species was tested by TWA or t-test respectively for each phosphosite. For acute temperature response, species and fold change upon step (up or down) and their interaction were variables; the interaction p value presented above each subpanel. For adapted temperature response, fold change (high/low) was tested against species.

Phosphopeptide motif analysis was performed with motif-x R package (*176*) and enrichment-depletion sequence logos were created using dagLogo R package (*177*), with foreground and background datasets, and a cutoff of $p < 0.05$. Foreground datasets were created by subsetting total data into subsets that changed in a proportional (log fold change up > 0 AND log fold change down < 0) or inversely proportional (log fold change up < 0 AND log fold change down > 0) direction. Background datasets were created for each direction comprising the total dataset. Enrichment-depletion sequence logos were separated to obtain enrichment motifs (Fig. 3D) and depletion motifs (fig. S7G) for each direction.

To compare the magnitude of proteome and phosphoproteome change, vectors were calculated for each protein or phosphosite comprising their $\log_2$ fold change under shift up and shift down conditions as illustrated (fig. S7B). Vector magnitude for all proteins was compared with that for phosphosites by Wilcoxon signed-rank test.

Immunoblotting

Samples for denaturing polyacrylamide gel electrophoresis (SDS-PAGE) were prepared by diluting lysates with reduced NuPage™ LDS sample buffer and heating at 70°C for 10 min. Samples were run on NuPage™ Novex™ 4-12% Bis-Tris protein gels in MES buffer. For Western blotting for Halotag or puromycin, chemiluminescence detection was used. Proteins were transferred from the gels to nitrocellulose membranes using an iBlot system (ThermoFisher). Membranes were stained by Ponceau as control for total protein loading, then washed, blocked, and incubated with primary antibody in the blocking buffer at 4°C overnight. Mouse anti-puromycin (PMY-2A4-2 from Developmental Studies Hybridoma bank, at 1:1000), rabbit anti-Halotag (Promega, #G9221, 1:3000), rabbit anti-S6K (Abcam, #ab32359, 1:1000), rabbit anti-S6K pT389 (Cell Signaling, #9234S, 1:1000), anti-4EBP1 (Cell Signaling, #9644S, 1:1000), rabbit anti-4EBP1 pT37/T46 (Cell Signaling, #2855S, 1:1000), or anti-β-actin (SCBT, #SC-47778 (C4), 1:1000) were used with 5% milk in TBST blocking buffer, and an anti-mouse or anti-rabbit HRP-conjugated secondary antibody. Immobilon HRP substrate reagents (Millipore, #WBKLS0500) were used to detect chemiluminescence (ChemiDoc MP; Bio-Rad). Gels were stained with coomassie blue (Severn Biotech, #30-38-50). Images were analyzed by densitometry in ImageLab v4.1 (BioRad).

Zoonomia analysis

We used RERconverge (*178*) to identify genes whose evolutionary rates significantly correlate with diurnality in mammals. Genes with low relative evolutionary rates in diurnal mammals compared to nocturnal mammals are likely to be under greater evolutionary constraint in diurnal species. Higher gene relative evolutionary rates in diurnal versus nocturnal mammals may be due to relaxed evolutionary constraint (implying decreased functional importance) or positive selection (implying divergent functional importance). Filtered alignments for gene amino acid sequences were used to estimate tree branch lengths (*104*), with 16,209 total genes or 11259 ubiquitous genes as defined by expression in fibroblasts within 1.5 SD of the mean expression across 68 tissues and cell lines from RNA-Seq data from the Genotype-Tissue Expression (GTEx) Project. Median RNA-Seq data was accessed from the GTEx Portal on 12 Nov 2024 (GTEx_Analysis_v10_RNASeQCv2.4.2_gene_median_tpm.gct.gz).

We ran RERconverge using default parameters on a set of 186 mammal species for which phenotypes could be defined and had genome information. This included a foreground set of 77 species defined as diurnal or crepuscular and the remaining 109 species defined as nocturnal (table S2). Relative evolutionary rates were calculated using 'getAllResiduals' with the gene trees as input. The binary trait tree was defined using the 'foreground2Tree' function. We set the transition argument to 'transition = "unidirectional"' allow only transitions to the foreground state. We set the clade argument to "clade=all", where maximum parsimony is used to infer where transitions to diurnality occur on the tree and set those branches, along with all daughter lineages, to foreground. Paths were then generated from the trait tree using the 'tree2paths' function. We performed correlation analyses between the paths and the relative evolutionary rate of each gene using the correlateWithBinaryPhenotype function, with the minimum number of species in a gene tree for inclusion in the analysis set to 20 ('min.sp=20'), the minimum number of foreground species set to 5 ('min.pos=5'). P values were adjusted for multiple comparisons using the Benjamini-Hochberg procedure. The hypothesis that mTOR- and WNK-related genes were targets for evolutionary change was tested by filtering the total dataset for genes in these pathways and repeating multiple comparisons adjustment using the Benjamini-Hochberg procedure. To account for any phylogenetic signal that may lead to significant associations that are independent from the diurnality trait we conducted phylogeny-aware permutations ('permulations') using the getPermsBinary function with default settings and 1,000 permutations. We considered any genes with both a adjusted p value < 0.05 from the correlation tests and a permuted p value < 0.05 to be significantly associated with being a diurnal mammal (table S3).

Gene Ontology and KEGG pathway analysis was performed using g:Profiler (*179*). Redundant GO:Process terms were removed using REVIGO. Top terms were retained and plotted according to whether they were conserved (Rho < 0) or evolved (Rho > 0) in diurnal mammals.

WNK1 protein disorder was calculated per residue per species (fig. S8G) using Metapredict v3 via a Google Colab notebook (*160*). The activity phenotype-WNK1 disorder relationship was assessed by calculating median ensemble disorder score of full-length WNK1 for each species, and comparing the median disorder score of diurnal vs nocturnal species by Wilcoxon signed-rank test (Fig. 3L).

*In vivo* mTOR activity reduction

To assess the effect of reduced mTOR activity on diurnal activity pattern, mice were fed a methionine-reduced diet (methionine content 14% of control) as formulated in (*180*)(Research Diets Inc, methionine reduced: #A11051301Bi, control: #A11051302Bi). 23 mice (12 males, 11 females) were singly housed in running-wheel cages with environmental enrichment under long day lighting conditions with natural twilights (*181*) corresponding to the following lighting schedule: 2h light increasing:14h light:2h light decreasing:6h dark. Behavioral wheel running was monitored throughout. Mice were fed *ad libitum* with standard diet (Teklad global 18% protein, Inotiv) for 1 week to acclimatize to single housing. After 1 week, mice were switched onto control or methionine reduced diet (N=5-6/sex/group) *ad libitum* for 4 weeks and monitored by behavioral analysis under LD conditions. After 4 weeks, mice were switched into constant darkness, continuing with control or methionine reduced diet, *ad libitum* for 1 week with behavioral monitoring. At the end of this period, at CT6 on day 42, all mice were culled by cervical dislocation and exsanguination, with whole brain and liver tissue taken and snap frozen in liquid nitrogen for subsequent protein extraction and immunoblotting. Mice were weighed weekly throughout to monitor weight loss. Mouse work was overseen by the Animal Welfare and Ethical Review Body of the MRC Laboratory of Molecular Biology.

For work for food (WFF), data was extracted from previously published work (*125*) for reanalysis, with experiments described as in (*125*, *127*). Experiments were approved by the University of Groningen ethical committee (DEC 5011, 5114, 5454) and carried out following national animal welfare regulations. Adult male mice (n = 24, CBA/CaJ) were kept singly housed in individual cages containing environmental enrichment, water *ad libitum* and equipped with running wheels to enable monitoring throughout the experiment. Mice were implanted under isoflurane (Abbott Laboratories Ltd., Kent, UK) inhalation anesthesia (2%) with customized temperature loggers (Thermochron iButton, DS1922L, Maxim Integrated Products Inc., Sunnyvale CA, USA) into their abdomen to enable core body temperature monitoring. Cages were placed in 12:12 light:dark cycles for the duration of the experiment (48 days). Mice were split into two groups: control, fed *ad libitum* for the duration, and work for food, where food availability was linked to wheel running and delivered after a set number of wheel revolutions. On days 1-10, 100 revolutions were required per food pellet, equivalent to an *ad libitum* food condition. At day 10, workload was increased daily by an extra 10–20 revolutions per pellet up to 260–330 revolutions per pellet by day 40. 260-300 rev/pellet was maintained for one week during days 40-47 during the test period. Throughout the experiment body mass was closely monitored at least every 3rd day at different times of day and high workload levels were individually titrated to keep the mice above 18 grams and 75% of their initial body mass.

To create an average daily activity, activity (running wheel counts) was grouped into 24 x 1h time bins for each day and time bins averaged across 7 days of the baseline (WFF: days 2 to 9) and test (WFF: days 40 to 47; methionine restriction: days 21 to 28) periods of the experiment. Averaged counts per time bin were subsequently normalized by the total counts per day, to give hourly activity as a proportion of the total activity in each day. Acrophase was determined by curve fitting (as described below) to normalised activity (counts/sum of counts during time window) and body temperature (WFF only) during baseline (WFF only) and test periods of the experiment. Phase was calculated as described below, where the peak of the curve fit represents the timing of the highest activity or body temperature. Activity onset was determined using automated onset detection in ClockLab (Actimetrics Inc.). To assess diurnality index (WFF only, for activity), normalized activity in daylight hours (6am to 6pm) of the average day was summed and expressed as a % of total activity occurring during the day.

<u>Statistics</u>

For bioluminescence data, all raw data was detrended using a 24h moving average and normalized (where 0 = average detrended luminescence and 1 = maximum detrended luminescence). For curve fitting, the following equation was used: $y=(mx + c) + ae^{kx}\cos(2\pi x - r/p)$ where m is the baseline, c is the offset from 0 in y-axis, a is the amplitude, k is the damping rate, r is the phase (in radians), and p is the period (*165*). Curve fitting was performed for constant condition data at least 12h after the last transition in temperature, or treatment, until 96h in constant conditions. Circadian phase was calculated from the timing of peaks of the fitted curve (designated as > 10% above baseline) using the area under the curve test on Prism. The time in hours of the first peak after entry to constant conditions was converted into circadian phase in hours by dividing by circadian period, and is given relative to the last transition between cold and hot such that the start of the hot cycle = 0h. Phase shift for the phase response curve was calculated as the difference in phase between cells that changed temperature and cells that stayed at constant temperature, and was given by $phase\ shift = phase_{shifted} - phase_{control}$. Circadian phases and circadian periods between species ± conditions were compared by either Watson-Williams test for homogeneity of means (R package “circular”: watson.williams.test), Correlation Coefficient for Angular Variables (R package “circular”: cor.circular with permutations), or two-way ANOVA (TWA) with Šídák’s post-hoc test for individual comparisons as stated in figure legends.

Mean ± SEM are reported throughout unless otherwise stated in figure legends. Error bars are not displayed if they are smaller than the size of the point to which they relate, due to a restriction in GraphPad Prism (v10, GraphPad Software). Statistical tests were performed using GraphPad Prism (v10) and R v4 and are indicated in figure legends. *P* values are either reported in figures directly or annotated with asterisks: * $p \leq 0.05$; ** $p \leq 0.01$; *** $p \leq 0.001$; **** $p \leq 0.0001$; n.s. not significant, $p>0.05$. Number of replicates are reported as n or N (for technical and biological, respectively) in the figures; error bars represent standard error (SEM) unless stated otherwise.

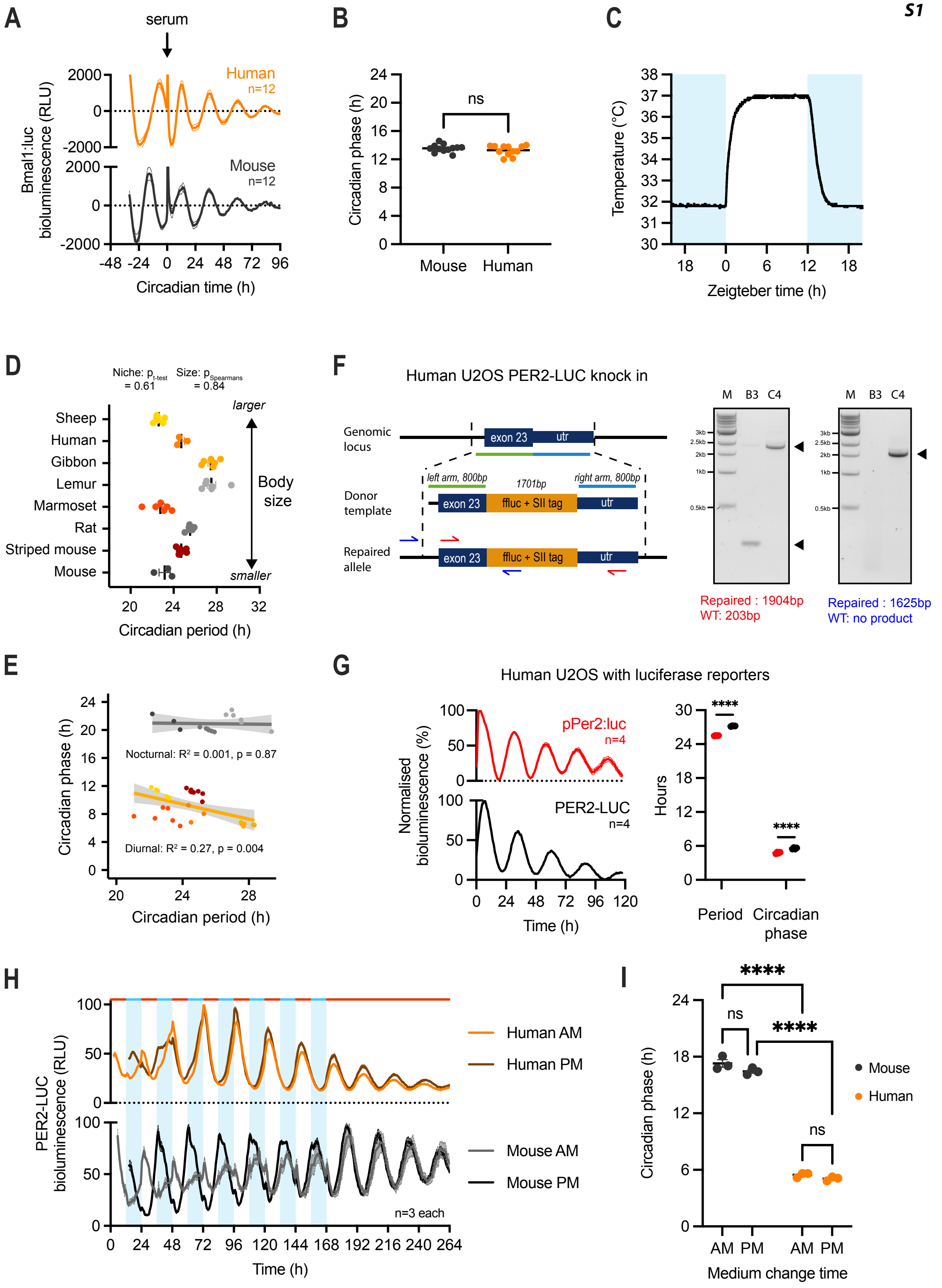
A
serum
Human
n=12
Mouse
n=12
Bmal1:luc bioluminescence (RLU)
Circadian time (h)
B
ns
Circadian phase (h)
Mouse
Human
C
Temperature (°C)
Zeigteber time (h)
D
Niche: p t-test = 0.61
Size: p Spearmans = 0.84
larger
smaller
Body size
Sheep
Human
Gibbon
Lemur
Marmoset
Rat
Striped mouse
Mouse
Circadian period (h)
E
Nocturnal: R² = 0.001, p = 0.87
Diurnal: R² = 0.27, p = 0.004
Circadian phase (h)
Circadian period (h)
F
Human U2OS PER2-LUC knock in
Genomic locus
Donor template
Repaired allele
exon 23
utr
left arm, 800bp
1701bp
right arm, 800bp
ffluc + SII tag
M B3 C4
Repaired : 1904bp
WT: 203bp
Repaired : 1625bp
WT: no product
G
Human U2OS with luciferase reporters
pPer2:luc
n=4
PER2-LUC
n=4
Normalised bioluminescence (%)
Time (h)
Hours
Period
Circadian phase
H
Human AM
Human PM
Mouse AM
Mouse PM
n=3 each
PER2-LUC bioluminescence (RLU)
Time (h)
I
ns
****
Mouse
Human
Circadian phase (h)
AM PM AM PM
Medium change time

**Fig. S1.**

**(A)** Primary fibroblasts from mouse and human, stably expressing *Bmal1:luc*, were given a medium change containing 10% serum at t = 0h at constant temperature. Following medium change, *Bmal1:luc* oscillated in the same phase. Detrended *Bmal1:luc* bioluminescence shown. **(B)** Circadian phase, calculated from (A). **(C)** Temperature profile of a 37 °C : 32 °C cycling incubator, used in temperature entrainment experiments. Incubators take approximately 3-4h to reach set temperature. Temperature profile is given against "Zeitgeber" time, where 0h is equivalent to the transition from 32 °C to 37 °C, at start of the 37 °C period. In all subsequent phase data, phase = 0h is equivalent to the start of the 37 °C period. **(D)** Circadian period of mammalian cell lines used in Fig. 1. Period was compared between the diurnal and nocturnal niche by t-test, and against ranked body size by Spearman's correlation. **(E)** Circadian period against circadian phase of mammalian cell lines. Regression lines for nocturnal (grey) and diurnal (yellow) are plotted. A small but significant association was observed between phase and period for the diurnal cells. **(F)** To examine the human cellular circadian biology, we developed an endogenous hPER2-LUC reporter using CRISPR-Cas9 mediated HDR in U2OS cells (*182*), a well-established model for human cellular circadian rhythms (*28*, *183*, *184*). PER2-LUC homozygous knock-in was achieved in the clonal cell line, C4, as seen in agarose gel of HDR PCR (red) and external PCR (blue). **(G)** Knock-in did not affect circadian dynamics of U2OS cells when examined with an alternative (transcriptional) reporter, pPer2:luc (red), after temperature cycles for 6 days before release into constant temperature, as previously observed for PER2 knock ins with fluorescent proteins (*184*). Only modest effects on circadian period and phase were seen. Statistics: Two-way ANOVA followed by Holm-Šídák's post-hoc test. **(H)** Human (U2OS) and mouse (fibroblasts, immortalized) PER2-LUC cells were synchronized by 10% serum-containing medium change at t=0h (AM) and t=12h (PM) and subject to temperature cycles (12 h 37 °C: 12 h 32 °C) starting at t=0h. Medium change in the morning (human) and afternoon (mouse) matches the applied temperature cycle. Medium change at the converse time of day is in conflict to the new timing cue, and human and mouse cells entrain within 7 days. Once entrained, cells exhibit a stable and opposite entrained phase in constant conditions **(I)**.

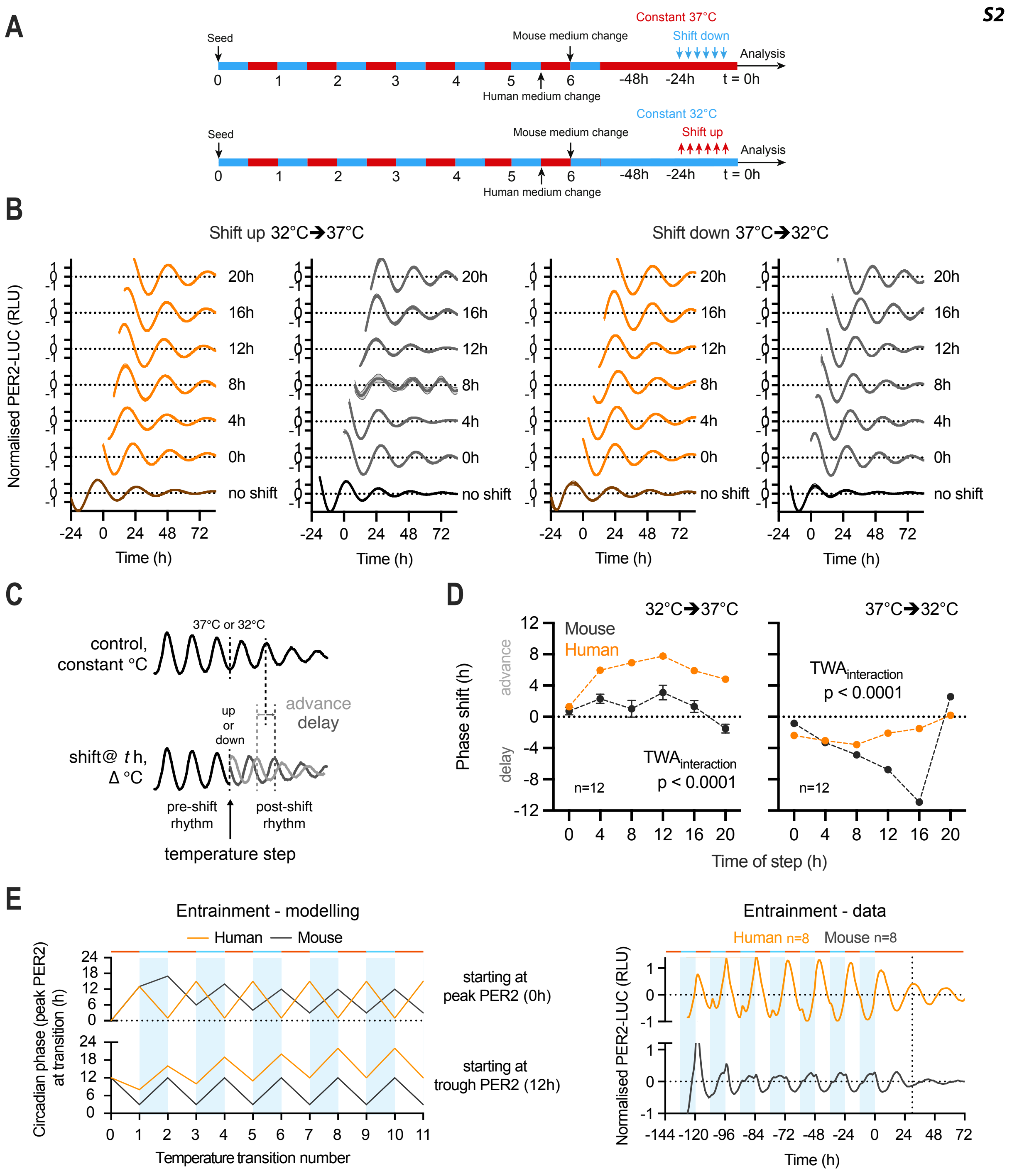


**Fig. S2.**

**(A)** Schematic of phase response curve experiment. Cells were plated on day 0 and entrained in temperature cycles. After 5 days, cells were given a medium change containing luciferin at a coherent phase when serum response matches the temperature response (fig. S1H). After 6.5/7 days, cells entered constant conditions of either 37°C or 32°C. At t=-24h cells were stepped between temperatures every 4h. Analysis of data began 24h after stepping.

**(B)** Detrended bioluminescence data from phase response curve experiment with step times given relative to peak PER2 (at 0h). Human (orange) and mouse (grey), n=4 each timepoint and cell line.

**(C)** Schematic of quantification of phase response curve experiment. Illustrative rhythms are shown for phase advance (grey) or delay (black) in the shifted condition relative to the control.

**(D)** Phase shift (phase shift = $phase_{stepped}$ - $phase_{control}$) is plotted against time of temperature step, either up (left) or down (right), for mouse and human cell lines expressing PER2-LUC, raw data is displayed in fig S3B. Statistics: TWA, interaction step up - species x time $F(6, 132) = 13.52$, $p < 0.0001$, interaction step down - species x time $F(6, 134) = 132.0$, $p < 0.0001$.

**(E)** Under a temperature cycle, cells receive repeated temperature steps (transitions) from cold to hot and hot to cold. Due to the phase shifts described in the PRC (D), cells do not encounter at these repeated transitions at a constant circadian phase. Instead, the phase of cells gradually changes in response to the previous transition, leading to the next transition occurring at a different circadian phase. (Left) Mathematical modelling of circadian phase (given by peak PER2 expression) in human and mouse cells at transition points, encompassing data from (D), reveals stable (and opposite) entrainment after 1-8 transitions depending on starting phase and cell type. This agrees with observed data (Right, data extended from Fig. 1E). and is equivalent to 0.5 to 3.5 days of 12h:12h temperature cycles.

**S3**

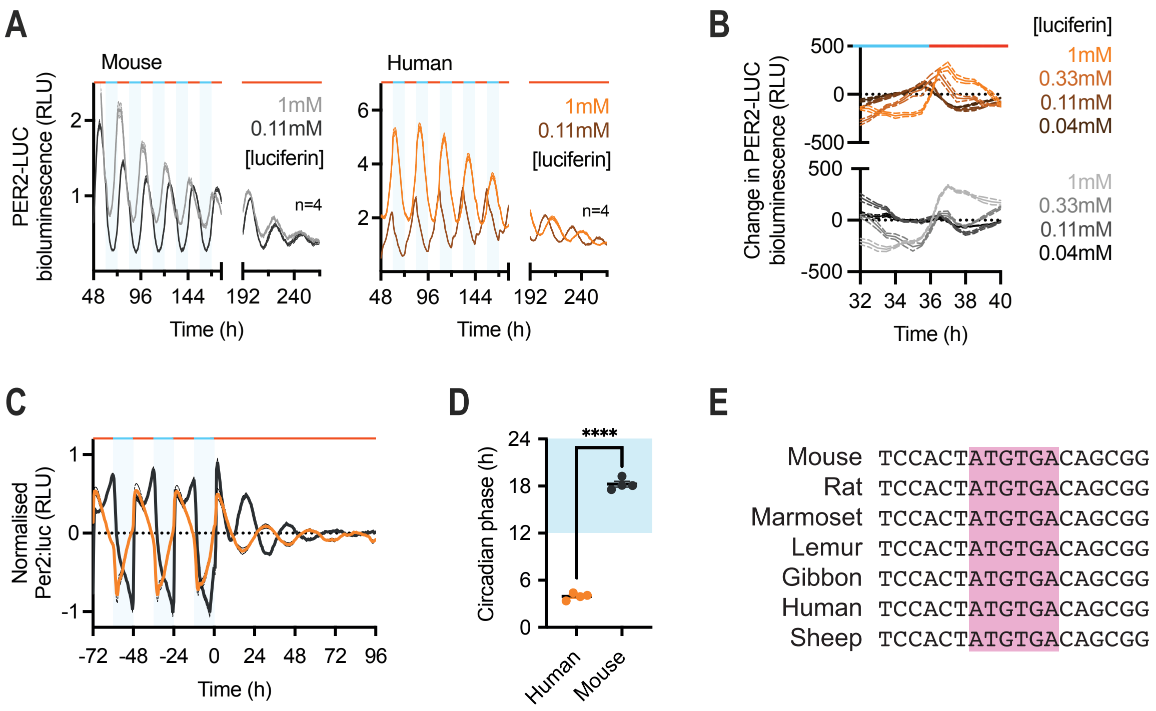


**Fig. S3.**

**(A)** Rhythmic expression of luciferase from the PER2 locus in human cells (orange) was similar to that from the well-characterized mouse PER2-LUC fibroblast cells (grey), including the relationship of the phase of the reporter to luciferin concentration after 7 days of 37°C : 32°C temperature cycles in the presence of different concentrations of luciferin (*69*).

**(B)** Under temperature cycling conditions, high concentrations of luciferin reveal temperature-dependent synthesis of PER2-LUC, here shown for a step up from 32 °C to 37 °C.

**(C)** Expression of luciferase under the control of the Per2 promoter (*Per2:luc* transcriptional reporter) in Human (U2OS, orange) and mouse (NIH 3t3, grey) can be deconvolved into acute temperature-responsive and circadian-driven in 37°C : 32°C temperature cycles at high concentrations of luciferin. Initially, Per2 transcription in both cells showed a similar circadian phenotype to other reporters in these two species (e.g. PER2-LUC and *Bmal:luc*). In temperature cycles, the reporter of Per2 transcription in both cells initially responds similarly to changes in temperature, increasing with temperature increase and decrease with temperature decrease, agreeing with recent reports on temperature response of Per2 (*52*). In the following hours, as the signal from the acute temperature response diminishes due to rapid catalytic inactivation of luciferase (*69*), the circadian-driven component of the signal is visible. Since the circadian rhythm underlying is opposite in the two species, this is seen as a gradual decay or increase upon transition to 37°C in mouse and human respectively, and vice versa upon transition to 32°C.

**(D)** As with other reporters (PER2-LUC and *Bmal:luc*), opposite entrainment to temperature cycles is observed between the two species.

**(E)** *Per2* uORF, first identified as an important regulator of temperature-dependent Per2 protein synthesis in (*52*), was compared between the diurnal and nocturnal mammals studied. *Per2* uORF is conserved in all species (*52*) including *Lemur catta*. Genomic information for Lemur catta was extracted from NC_059135.1 and Genome Data Viewer surrounding predicted *Per2* locus, with Blastn conducted to identify uORF location.

***S4***

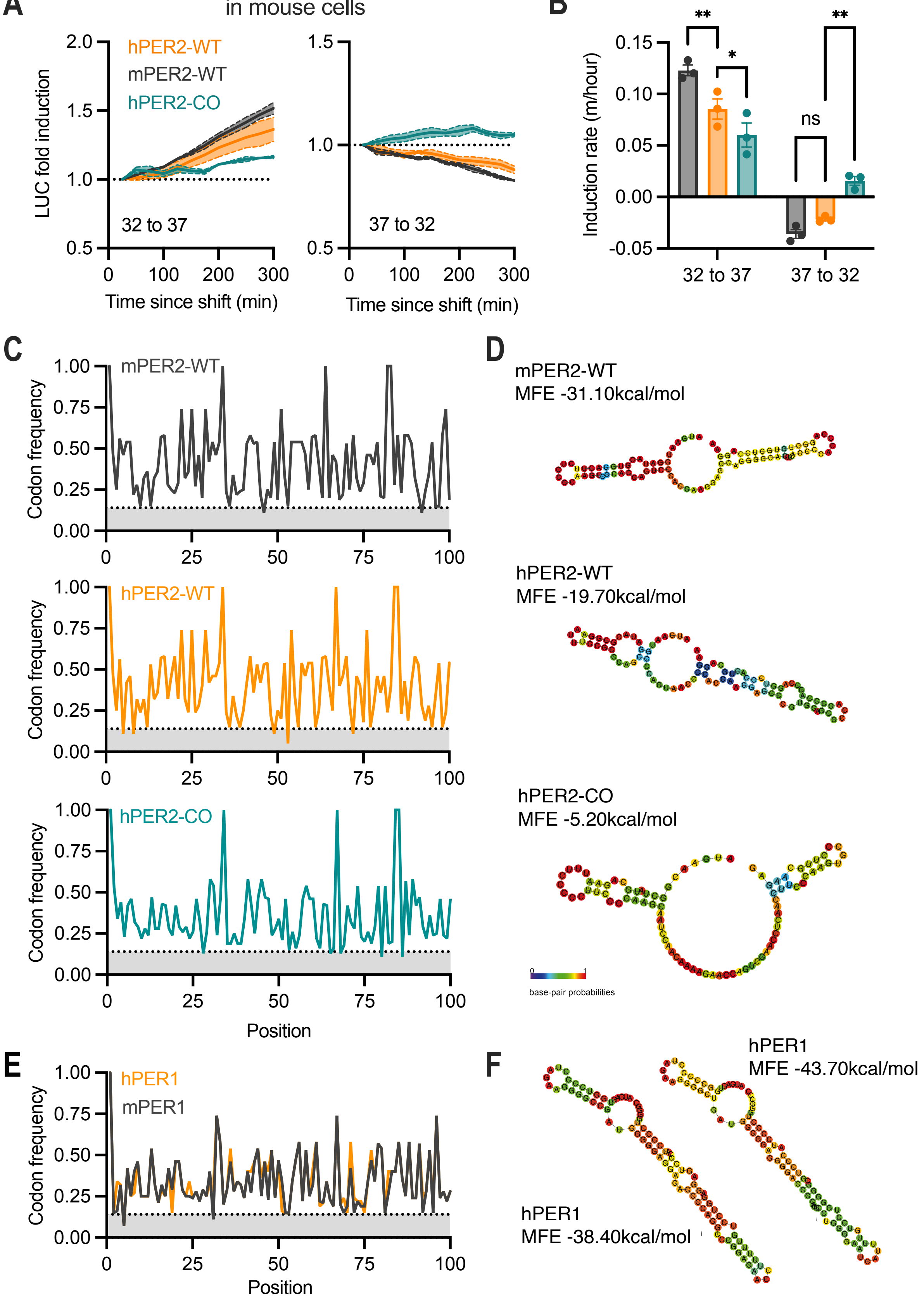
A
in mouse cells
hPER2-WT
mPER2-WT
hPER2-CO
LUC fold induction
32 to 37
37 to 32
Time since shift (min)
B
Induction rate (m/hour)
**
*
ns
**
32 to 37
37 to 32
C
Codon frequency
mPER2-WT
hPER2-WT
hPER2-CO
Position
D
mPER2-WT
MFE -31.10kcal/mol
hPER2-WT
MFE -19.70kcal/mol
hPER2-CO
MFE -5.20kcal/mol
base-pair probabilities
E
hPER1
mPER1
Codon frequency
Position
F
hPER1
MFE -43.70kcal/mol
hPER1
MFE -38.40kcal/mol

**Fig. S4.**

**(A)** Mouse NIH3t3 cells stably expressing constitutive PER2-LUC fusions (human PER2-LUC (orange), mouse PER2-LUC (grey) or codon-optimized human PER2-LUC (teal)) were kept for 3 days at constant temperature of either 32 °C or 37 °C before shifting temperature at t = 0 min. Fold induction of luminescence is shown relative to t = 0 min.
**(B)** Rate of induction was quantified as the gradient, m, of the straight line fit, non-linear regression (Prism). Induction rates were compared between reporters. Statistics: Two-way ANOVA (TWA) followed by Holm-Šídák's post-hoc test.
**(C)** Rare codon usage is important for *Drosophila* Per function (*80*) and is a common functional feature of clock proteins which contain intrinsically disordered regions, like PER2 and FRQ (*185*, *186*). Codon usage impacts translation dynamics (*75*) and mRNA stability (*76*, *77*) through its effect on translation rate, protein folding and mRNA stability (reviewed in (*187*)). Codon frequency is plotted for the first 100 amino acids for each of the constitutively expressed PER2 proteins used in this study. Dashed line and grey shading indicates frequency less than 14%. Codon optimized human PER2 (hPER2_CO) contains fewer rare codons and a smaller range of codon frequency ($SD_{CO}$ 0.197) than wild type human PER2 (hPER2_WT, $SD_{WT}$ 0.215), and wild type mouse PER2 (mPER2_WT, $SD_{WT}$ 0.214).
**(D)** RNAFold predicts the initial 90 nucleotides of the mRNA encoding hPER2_CO contains less secondary structure (minimum free energy prediction, MFE, is less negative) than hPER2_WT, which contains less secondary structure than mPER2_WT. As translation initiation and early elongation phases dictate the efficiency of protein synthesis (*79*), these features within the first 30-100 amino acids of PER2 of codon usage and RNA structure suggest that these features contribute to the temperature sensitivity of PER2 translation and the loss of temperature sensitivity of the PER2_CO.
**(E and F)** PER1 is highly similar between mouse and human. (E) Codon frequency for the first 100 amino acids for human and mouse PER1. The range of codon frequency is the same ($SD_{human}$ 0.163, $SD_{mouse}$ 0.163). (F) mRNA structure prediction from RNAFold for the initial 90 nucleotides of mouse and human PER1.

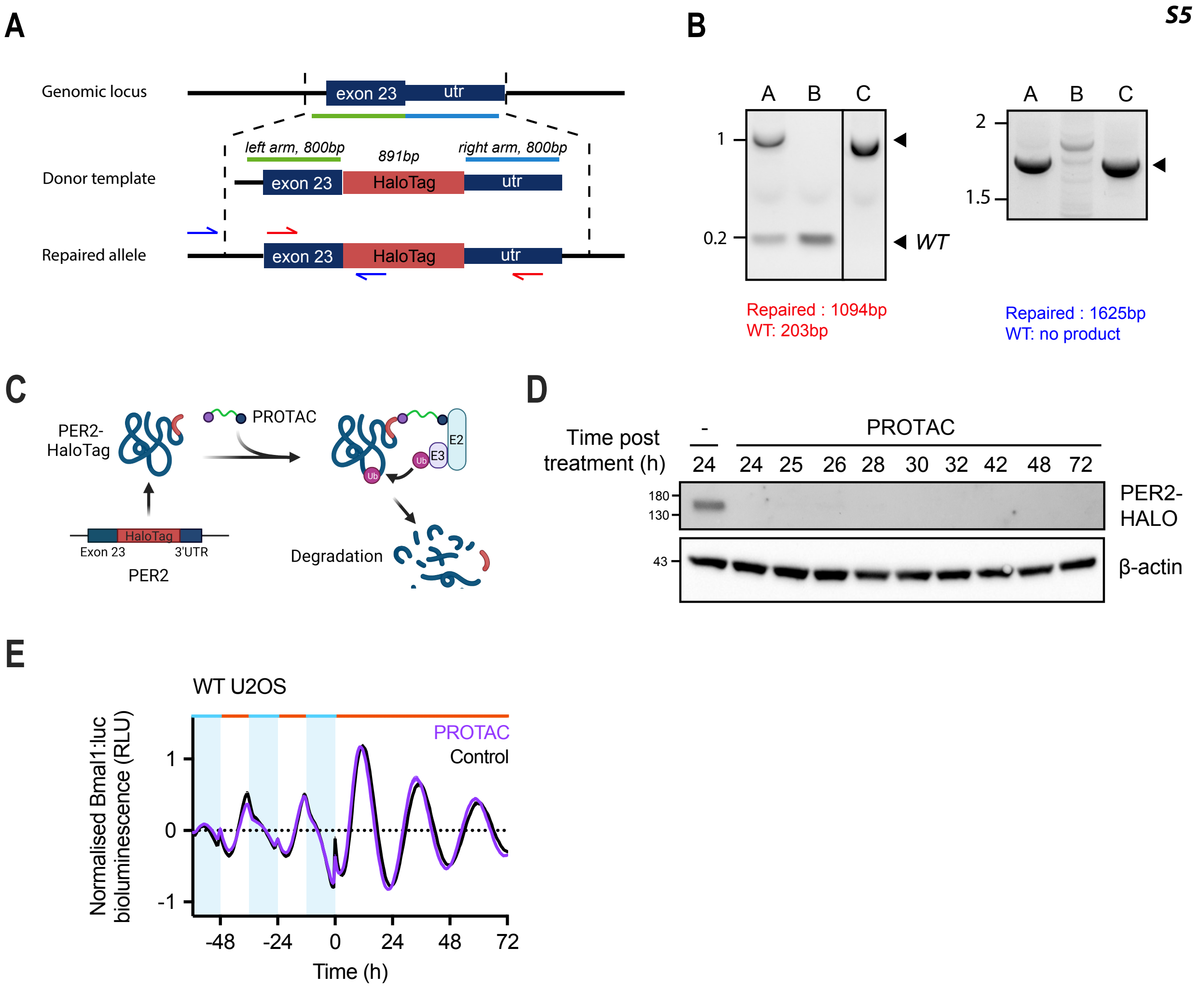


**Fig. S5**

**(A)** PER2-HaloTag knock in human U2OS cells were created using CRISPR-Cas9 mediated HDR. PER2-HaloTag heterozygous and homozygous knock in was achieved in the clonal cell lines, C and A respectively, as seen in agarose gel **(B)** of HDR PCR (red) and external PCR (blue).

**(C)** A proteolysis targeting chimera (PROTAC) specific to the HaloTag (HaloPROTAC3) specifically binds to and degrades HaloTag and its fusion partner PER2 via E3 ligase (VHL) and the proteasome, allowing precise control of PER2 levels in cells (*188*, *189*).

**(D)** PROTAC efficiently and stably reduces PER2 levels in cells. PER2-HaloTag cells were incubated with 500 nM HaloPROTAC3 (or DMSO control) for 24 h before sampling at the indicated times. Immunoblots of anti-Halo or anti-β-actin are shown.

**(E)** Wild type U2OS cells were cultured in temperature cycles ± 1 μM HaloPROTAC3 (n=4 each condition). At 72h after the start of temperature cycles, cells were kept in constant temperature. Bioluminescence from *Bmal1:luc* was recorded throughout.

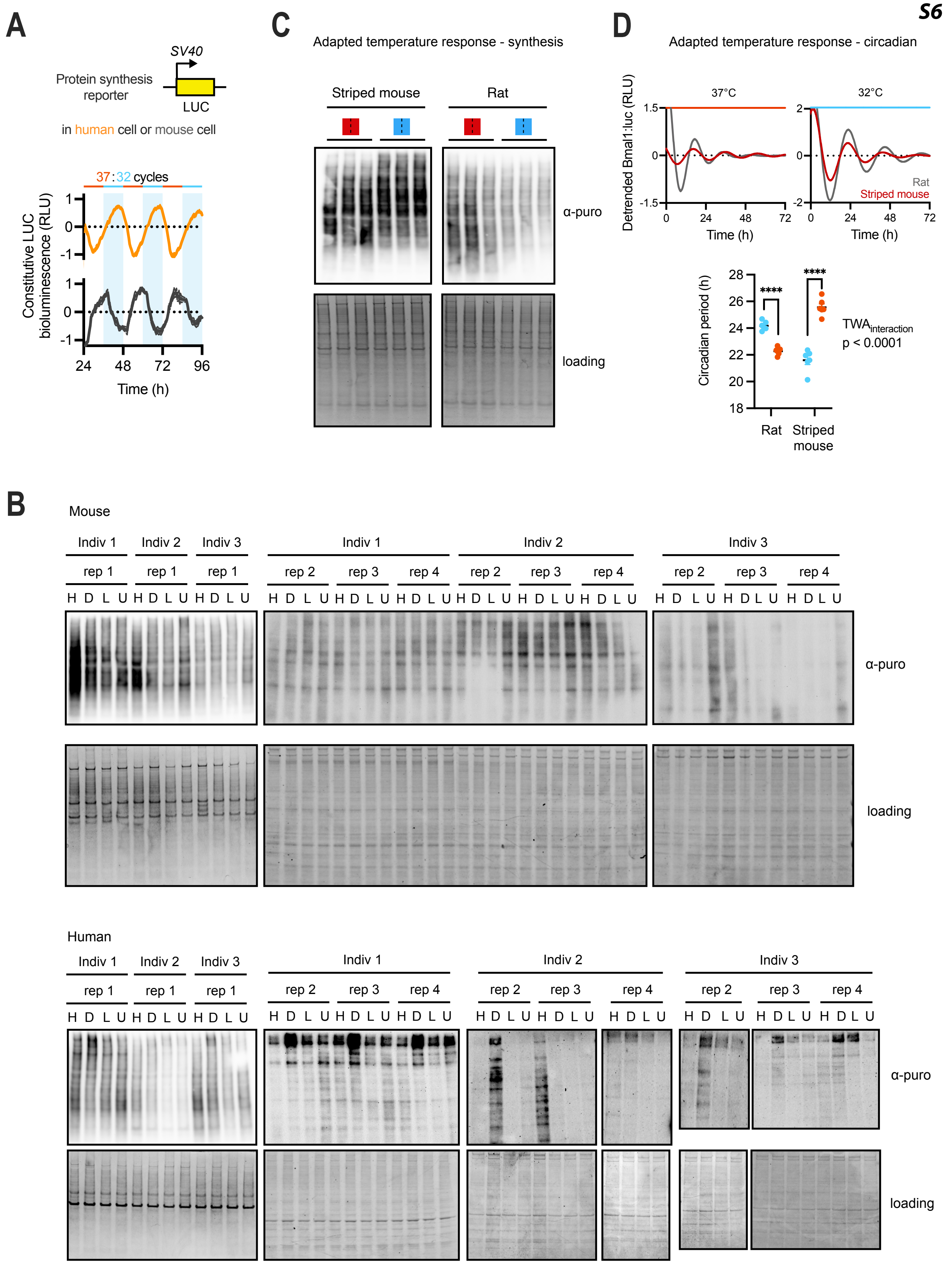
A
SV40
Protein synthesis reporter
LUC
in human cell or mouse cell
37:32 cycles
Constitutive LUC bioluminescence (RLU)
Time (h)
C
Adapted temperature response - synthesis
Striped mouse
Rat
α-puro
loading
D
Adapted temperature response - circadian
37°C
32°C
Detrended Bmal1:luc (RLU)
Time (h)
Rat
Striped mouse
Circadian period (h)
TWA interaction
p < 0.0001
Rat
Striped mouse
B
Mouse
Indiv 1
Indiv 2
Indiv 3
rep 1
rep 2
rep 3
rep 4
H D L U
α-puro
loading
Human

**Fig. S6.**

**(A)** Human and mouse cells expressing constitutive LUC (SV40:LUC), a proxy for protein synthesis, were cultured in temperature cycles as Fig. 1B. Protein synthesis oscillates in antiphase in the two species.

**(B)** Anti-puromycin (top) and coomassie loading control (bottom) for all replicates for temperature shift and adaptation experiment in Fig 2, H and I. 4 replicate cell lines from 3 individuals were used. Multiple separate gels are indicated by black boundaries – quantification (fold change) was performed relative to samples on the same gel.

**(C)** Anti-puromycin (top) and coomassie loading control (bottom) for all replicates of temperature-adapted puromycin incorporation in multiple species, quantified in Fig. 2I, n=3 per species.

**(D)** Under constant temperatures, temperature compensation of the circadian rhythms of rat and striped mouse primary fibroblasts (n=6) was examined. *Bmal1:luc* bioluminescence was recorded under constant 32°C (blue) or constant 37°C (red) conditions. Cells were kept at the indicated constant temperature for 7 days prior to the recording. 24h prior to recording, cells were given a complete medium change containing 10% serum. Raw bioluminescence data was baseline detrended and converted to circadian time. As shown in Fig. 2J, whilst the period of circadian rhythms in all species exhibit $Q_{10} \sim 1$, indicating temperature compensation (*84*), there are nocturnal-diurnal differences in the temperature dependence of circadian period. Nocturnal mammals are overcompensated i.e. circadian period ($\tau$) of the rhythm is shorter at 37°C than 32°C and $Q_{10} < 1$, while nocturnal mammals are undercompensated, $Q_{10} > 1$. This invites analogies with Aschoff's Rule (*190*) on the relationship between circadian period and light intensity. Circadian period is dependent on light intensity, but the direction of the relationship is opposite in nocturnal and diurnal species: for nocturnal $\tau_{LL} > \tau_{DD}$ whereas for diurnal $\tau_{LL} < \tau_{DD}$ (*191*). Our results suggest a version of Aschoff's Rule related to temperature: for nocturnal $\tau_{32} > \tau_{37}$ whereas for diurnal $\tau_{32} < \tau_{37}$.

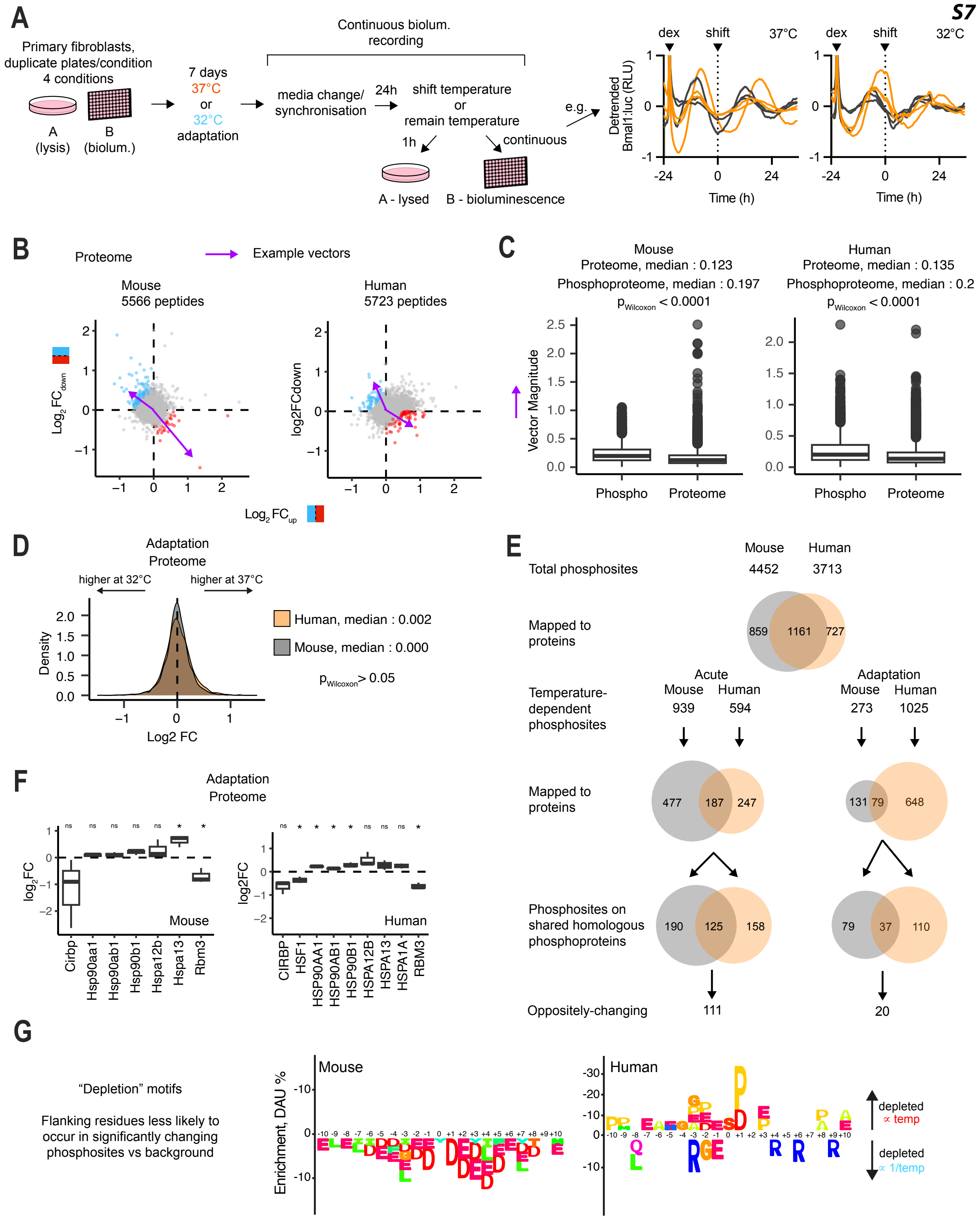


700

**Fig. S7.**

**(A)** Schematic of experiment. Primary fibroblasts of mouse and human cells were plated for four conditions (constant 37°C, shift down from 37°C to 32°C, constant 32°C and shift up from 32°C

to 37°C) in duplicate plates, one for lysis (plate A) and a parallel plate for bioluminescence monitoring (plate B), to ensure phase alignment between species at treatment (temperature shift) and lysis timepoints. Cells were adapted to 37°C or 32°C for one week, before a synchronizing pulse of dexamethasone to bring the cells back into phase alignment. Biological replicate cell lines exhibit different free running periods as has been described previously (*159*), which, over the course of one week, results in cells from different individuals drifting out of phase. After 24h, cells were either shifted temperatures, or remained at the adapted temperature, before lysis (for plate A) or continuing bioluminescence monitoring (plate B). Data for the 37°C and 32°C constant temperature bioluminescence recordings are shown, demonstrating general phase alignment between the species at the point of temperature shift after dexamethasone synchronization (dex).
**(B)** As Fig. 3B, proteome changes under shift conditions.
**(C)** Under shift conditions, in contrast to the phosphoproteome, the proteome does not change as much. Magnitude of phosphosite (Fig. 3B) and protein (A) vector lengths is compared for mouse (left) and human (right).
**(D)** Probability density distribution of fold change upon temperature adaptation for every detected protein. Mouse (grey) and human (orange). Statistics: Mann-Whitney, p-value shown. Under temperature adaptation, the mouse and human proteomes do not differ in response. Fold change.
**(E)** Comparison of cellular phosphoproteome temperature responses between mouse (grey) and human (orange). 4452 and 3713 individual phosphosites were quantified for mouse and human respectively after filtering for missing values (see Materials and Methods). These phosphosites mapped to 2020 and 1888 (phospho)proteins in mouse and human respectively. Of these quantified phosphoproteins, 1161 overlapped between human and mouse. Between 7% and 23% of quantified phosphosites were determined to be temperature dependent in acute and adapted conditions, between 70% and 77% of which mapped to individual (phospho)proteins. Of the temperature dependent phosphoproteins, the identity of a proportion was shared between mouse and humans: 187 phosphoproteins in acute conditions and 79 phosphoproteins in adapted conditions. These shared phosphoproteins had multiple phosphosites – however, only a proportion of these were homologous phosphosites in mouse and human. Of the homologous phosphosites, 111 and 20 phosphosites in acute and adapted conditions respectively changed in opposite directions.
**(F)** $Log_2FC$ for selected proteins of the heat shock and cold response pathways. Positive values indicate higher abundance at 37°C than 32°C. Negative values indicate higher abundance at 32°C than 37°C. Statistics shown are adjusted p-values from the entire proteome dataset. Overall, direction of response in these pathways is conserved between mouse (left) and human (right).
**(G)** Depletion motifs, centered around the phosphoacceptor at position 0, for proportional and inversely proportional direction for mouse and human, corresponding to Fig. 3D. AA residues displayed are significantly depleted in these direction changes relative to the background.

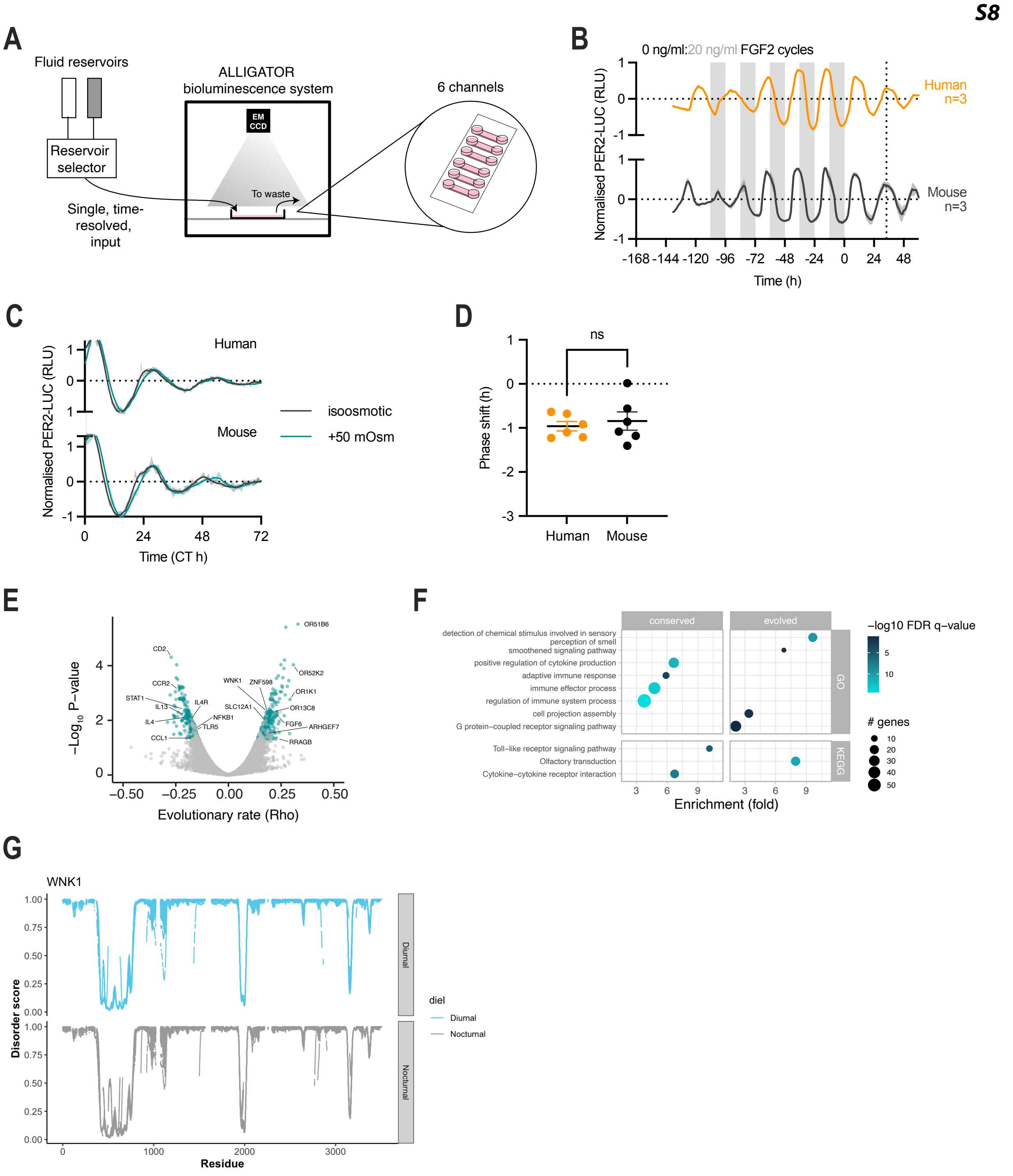


**Fig. S8**

**(A)** Schematic of bioluminescent experiments under continuous flow. A distribution valve selects medium from pressurized microfluidic reservoirs for durations and intervals as indicated in figure legends to produce a single, time-resolved input condition. This medium flows through cells on channel slides, which are situated inside an Alligator bioluminescence incubator to continuously monitor bioluminescence. In this way, physiologically-relevant and daily variation in extracellular conditions can be closely modelled.

**(B)** Human U2OS PER2-LUC and immortalized mouse fibroblast PER2::LUC cells were exposed to cycles of FGF2-containing medium. Cells were placed under flow at -144h in regular flow medium without supplementary FGF2. After 36h, FGF2-containing medium (grey) was flowed over cells for 12h before switching to FGF2-free medium (white) for 12h. In this way, five 0 ng/ml (white) : 20 ng/ml (grey) FGF2 cycles were performed, before release into 0 ng/ml FGF2 medium for 60h. Unlike temperature cycles or osmotic cycles, FGF2 cycles entrain mouse and human cells to the same circadian phase.

**(C)** Human U2OS PER2-LUC and immortalized mouse fibroblast PER2::LUC cells were given a medium change at t = 0h at constant temperature. Medium was either isosmotic or hyperosmotic (+50 mOsm, adjusted using sucrose) to the medium the cells were previously adapted to. Detrended bioluminescence shown.

**(D)** Circadian phase, calculated from (F). There was no significant difference in the phase shift in rhythm after increasing medium osmolarity by 50 mOsm in cells from human and mice.

**(E)** RERconverge, which tests for associations between relative evolutionary (substitution) rates of genes and the evolution of traits (*178*, *192*), identified 478 of 16209 genes whose evolutionary rates significantly correlate with diurnality in mammals (table S3). To account for any phylogenetic signal that may lead to significant associations that are independent from the diurnality trait across the mammals, for example in the close Family-level grouping of many diurnal or nocturnal species, we conducted phylogeny-aware permutations (pFDR < 0.05). Of the significant genes, 187 genes and 291 genes were identified as evolving slower (implying conservation) or faster (implying selection or relaxed evolutionary constraint) respectively in diurnal relative to nocturnal mammals.

**(F)** Genes with a significant rate change in diurnal mammals vs nocturnal mammals were processed for Gene Ontology and KEGG pathway analysis using g:Profiler. Redundant GO:Process terms were removed using REVIGO. Top terms were retained and plotted according to whether they were conserved (Rho < 0) or evolved (Rho > 0) in diurnal mammals. Conserved genes were enriched for gene ontology (GO) terms related to immune response and the most enriched KEGG term was for Toll-like receptor signaling. Faster-evolving genes were nominally enriched in gene sets related to signal transduction, in particular olfactory signaling, as may be expected from the diverse sensory challenges between the highly different environmental niches occupied by diurnal and nocturnal mammals.

**(G)** We asked whether diurnal WNK1 tended towards less disorder than nocturnal WNK1 across the mammalian family tree. WNK1 protein disorder was calculated per residue per species using Metapredict v3 via a Google Colab notebook (*160*) on amino acid sequences and alignments from the Zoonomia resource (*103*, *104*). Median disorder score at each aligned amino acid position for diurnal and nocturnal species is presented. No clear difference in specific regions of WNK1 was observed; however ensemble disorder (calculated across the whole protein) was significantly different between nocturnal and diurnal species (Fig. 3L).

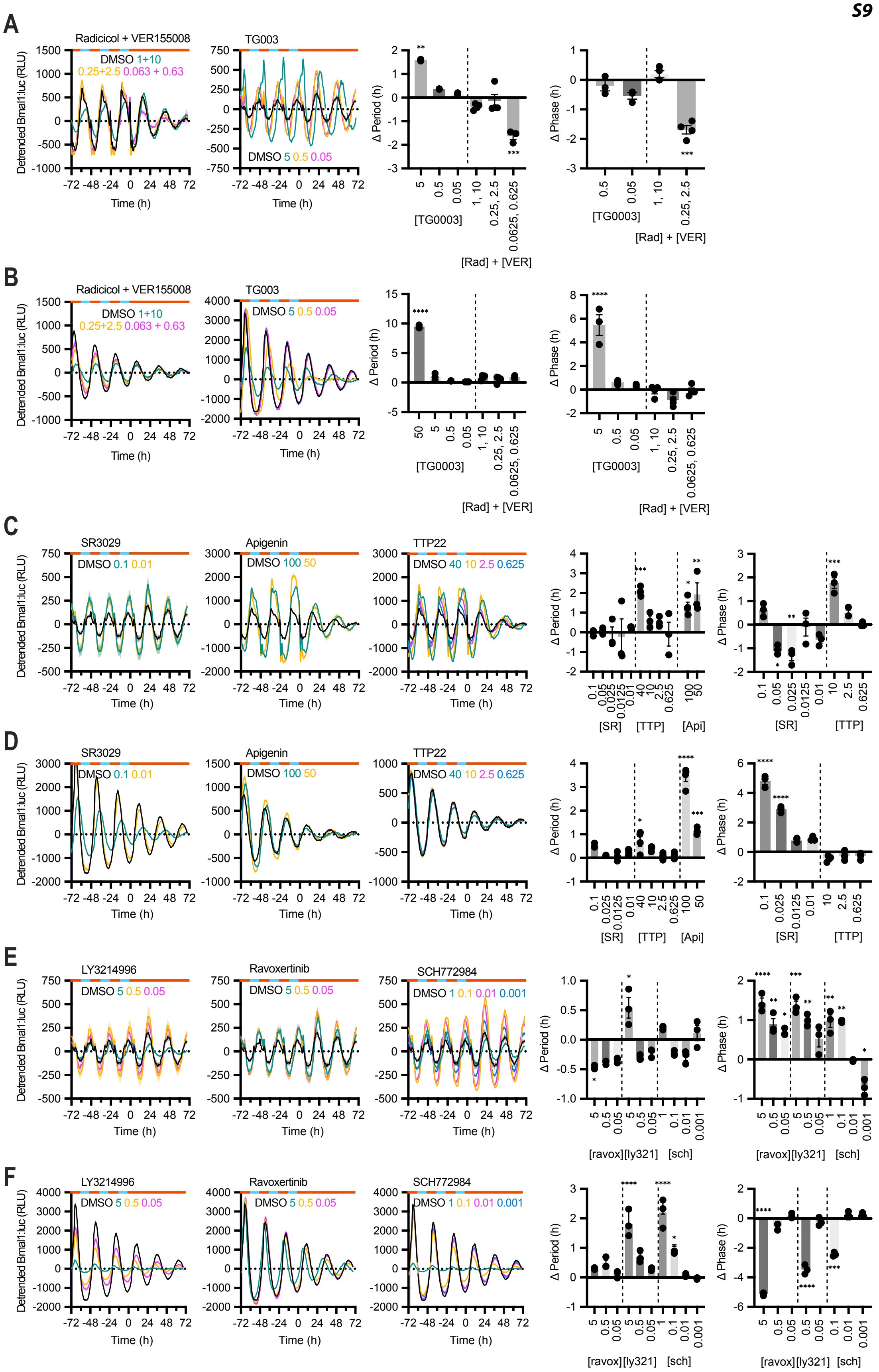
A
Radicicol + VER155008
DMSO 1+10
0.25+2.5 0.063 + 0.63
TG003
DMSO 5 0.5 0.05
Detrended Bmal1:luc (RLU)
Time (h)
Δ Period (h)
Δ Phase (h)
[TG0003]
[Rad] + [VER]
B
C
SR3029
DMSO 0.1 0.01
Apigenin
DMSO 100 50
TTP22
DMSO 40 10 2.5 0.625
[SR]
[TTP]
[Api]
D
E
LY3214996
Ravoxertinib
SCH772984
DMSO 1 0.1 0.01 0.001
[ravox][ly321]
[sch]
F

**Fig S9**

The heat-shock pathway and CLK kinases have been previously suggested to play a role in temperature entrainment and body temperature sensing (*46*, *51*). However, functional inhibition of HSF1 or CIRP does not completely inhibit temperature entrainment (*46*, *47*, *49*) suggestive of an alternative mediator(s) playing a role in temperature entrainment. Therefore, we examined the effects of a panel of small molecule inhibitors of proteins and kinases previously implicated in circadian post-translational regulation on entrainment to temperature cycles, to see if the activity of any kinase was responsible for the oppositely phased phenotype. Mouse and human PER2-LUC cells were synchronized by medium change at t=-186h, before re-entrainment in temperature cycles for 7 cycles. After 7 cycles, cells were released into constant conditions (at t=0). Cells were culture in the presence of inhibitors of HSP70/90, CLK, CK1, CK2, and ERK1/2 from the final medium change at t=-186h to test the effect of inhibitor on re-entrainment and entrained phase and bioluminescence monitored. A range of concentrations was used; phase of entrainment was only calculated for concentrations that did not affect circadian period, since entrainment phase is confounded by period length (Roenneberg et al., 2003).

**(A and B)** HSP70/90 inhibition (radicicol and VER15508) and CLK inhibition (TG003 (*51*, *193*)) in mouse (A) and human (B) cells. (Left) Bioluminescence data for mouse fibroblasts and (right) change in circadian period (ΔPeriod) relative to control for each condition. Inhibition of HSP70/90 or CLK1/4, at concentrations that did not affect period, only partly altered the entrained phases of human and mouse cells under daily temperature cycles, suggesting their function is not the primary cause of differential circadian temperature entrainment.

**(C and D)** CK1 inhibition (SR3029) and CK2 inhibition (Apigenin and TTP22) mouse (C) and human (D) cells. (Left) Bioluminescence data for mouse fibroblasts and (right) change in circadian period (ΔPeriod) relative to control for each condition. Whilst CK2 inhibition either changed period (Apigenin) or only partially affected phase (TTP22 10µM, mouse), CK1 inhibition had a larger and more consistent effect, dose-dependently delaying entrained phase in humans, and weakly advancing phase in mouse.

**(E and F)** ERK1/2 inhibition (LY3214886, Ravoxertinib, SCH772964) in mouse (E) and human (F) cells. (Left) Bioluminescence data for mouse fibroblasts and (right) change in circadian period (ΔPeriod) relative to control for each condition. In contrast to HSP, CLK, CK1 and CK2 inhibition, ERK1/2 inhibition had large and opposite effects upon human and mouse lines with respect to entrained phase: at the same concentrations, humans exhibited phase advances where mouse exhibited phase delays.

Statistics: Two-way ANOVA (TWA) followed by Holm-Šídák’s post-hoc test.

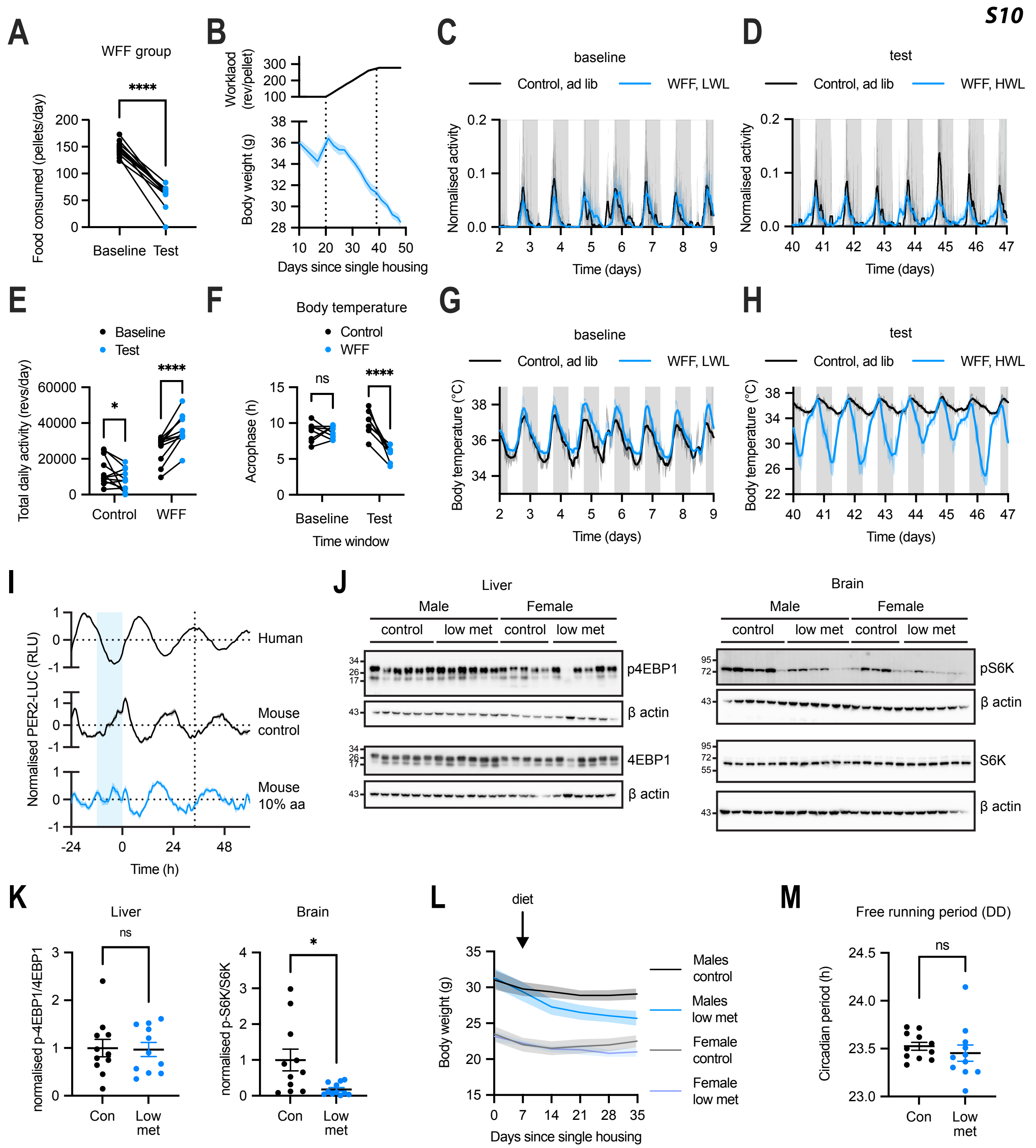


**Fig. S10.**

**(A to E)** WFF alters energy balance by providing a discrete number of calories (pellets) for gradually increasing energy expenditure without imposing any explicit feed:fast cycle (*125–127*). (A) In the test portion of the experiment (days 40-47, high work load, HWL), daily number of pellets is significantly less than at baseline (days 2-10, low work load, LWL). (B) Mice lose body weight as workload (revolutions of running wheel per pellet provided) increases. (C and D) Average proportion of total daily locomotor activity in 1h bins, plotted across the 7 days of each window of the experiment for control (n=10-11) and WFF groups (n=11-12). (C) baseline: *ad lib* feeding both groups; (D) test: WFF group, HWL; control group, *ad lib* feeding . Compared with control conditions (food *ad lib*) where mice are nocturnal, as energy balance becomes negative under WFF, mice progressively portion more of their activity to the light phase, like a diurnal mammal. (E) Under test condition, mice consume fewer pellets and lose body weight relative to the baseline condition. However, mice are running significantly more. There is a small but significant difference in wheel running in the *ad lib* control mice group after 39 days of single housing.

**(F to H)** As for (C and D) but presenting core body temperature, as recorded by implanted temperature sensors. (F) Acrophase was calculated by curve fitting to the 7 days of data in each time window. Statistics: TWA mixed-effects model, matched design. Interaction, baseline vs test x ad lib vs WFF $F(1, 17) = 123.5$, $p < 0.0001$, Šídák's post-hoc test reported.

For (C), (D), (G) and (H) mice were kept in 6am ON: 6pm OFF LD cycle throughout, and grey shading indicates dark phase.

**(I)** Mouse PER2-LUC fibroblasts were cultured in medium containing 10% total amino acids relative to control and entrained to 7 x 12h 37 °C: 12h 32 °C temperature cycles before constant 37 °C at 0h. Under amino acid restriction, mouse cells are phase-advanced relative to control.

**(J)** mTOR activity in liver (left) and brain (right) taken from mice after methionine restriction for 4 weeks, as indicated by immunoblot.

**(K)** Quantification of (A) and (B).

**(L)** Weights of mice (male and female) on diet (n=5 females control, n=6 males control, n=6 females low met diet, n=6 males low met diet).

**(M)** Weight loss (%) vs baseline (calculated between day 15 and 37) against acrophase (calculated as an average day between day 15 and 37, as Fig. 5I) for low met and control diet.

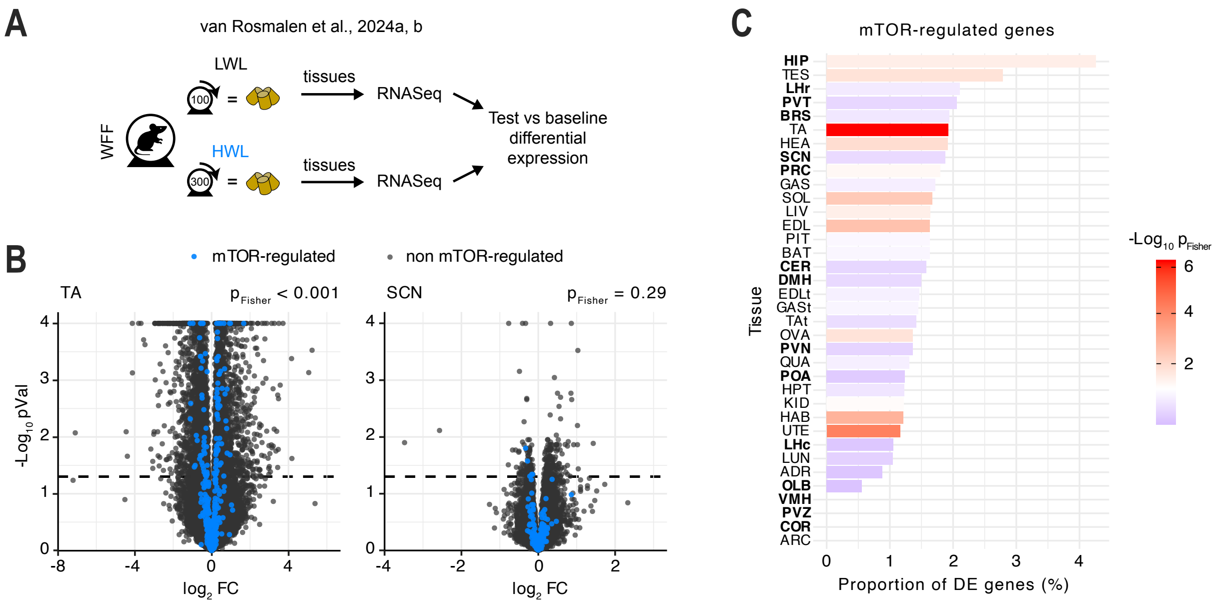


**Fig. S11. Differential gene expression analysis and mTOR pathway involvement across tissues in WFF mice**

**(A)** Tissue samples were collected from mice subjected to high workload (HWL) or low workload (LWL) conditions for RNA sequencing (RNA-seq), as reported by van Rosmalen et al. (*128*, *130*). Differential gene expression analysis was performed using DESeq2, comparing HWL to LWL within each tissue. Genes were classified as differentially expressed (DE) based on a Benjamini-Hochberg adjusted significance threshold of $p < 0.05$.

**(B)** Representative volcano plots from time- and sex-independent analyses illustrate gene expression changes in the (left) tibialis anterior muscle and (right) suprachiasmatic nucleus (SCN). mTOR-regulated genes (blue) were identified and annotated based on Düvel et al. (*194*). The proportion of mTOR-regulated DE genes was tested against the null hypothesis using Fisher's Exact Test, with $p$ values ($p_{Fisher}$) shown.

**(C)** Summary of $p_{Fisher}$ values and the proportion of mTOR-regulated genes among DE genes across all analyzed tissues from van Rosmalen et al. (*128*, *130*). Significance is color-coded: tissues where mTOR-regulated genes are significantly overrepresented ($p_{Fisher} < 0.05$) are shown in red, while non-significant results ($p_{Fisher} > 0.05$) are shown in blue.

**Table S1.**

Raw and processed data for human (A to D) and mouse (E to H) primary biological replicate (N=3 each) fibroblast proteomics (A, B, E and F) and phosphoproteomics (C, D, G and H) for adaptation conditions (A, C, E, G) and acute shift conditions (B, D, F, H). R scripts to generate tables from raw data exist at https://github.com/andrewbeale/temperature. 6809 and 6719 proteins were detected in mice and humans respectively. After filtering for proteins detected in all conditions, we quantified 5566 and 5723 proteins in mice and humans respectively. 22989 and 19236 phosphopeptides were detected in mice and humans respectively. After filtering for phosphosites detected in all conditions, we quantified 4399 and 3713 phosphosites in mice and humans respectively. For adaptation condition, log2 intensities under "high" or "low" conditions for each replicate are presented, alongside log2 fold change for each replicate, and FDR-corrected qVal. For acute condition, log2 fold change (up over low or down over high as illustrated in Fig 2B) is presented for each replicate with FDR-corrected qVal.

**Table S2.**

Diurnal and nocturnal activity scoring for species included in the analysis of evolutionary rate.

**Table S3.**

Correlation values (Rho) between relative evolutionary rates of 16209 genes and the binary phenotype of diurnal vs nocturnal mammals. Multiply adjusted p-value and phylogenetically-aware permutation p-value are shown.
